# Translational response to mitochondrial stresses is orchestrated by tRNA modifications

**DOI:** 10.1101/2024.02.14.580389

**Authors:** Sherif Rashad, Shadi Al-Mesitef, Abdulrahman Mousa, Yuan Zhou, Daisuke Ando, Guangxin Sun, Tomoko Fukuuchi, Yuko Iwasaki, Jingdong Xiang, Shane R Byrne, Jingjing Sun, Masamitsu Maekawa, Daisuke Saigusa, Thomas J Begley, Peter C Dedon, Kuniyasu Niizuma

## Abstract

Mitochondrial stress and dysfunction play important roles in many pathologies. However, how cells respond to mitochondrial stress is not fully understood. Here, we examined the translational response to electron transport chain (ETC) inhibition and arsenite induced mitochondrial stresses. Our analysis revealed that during mitochondrial stress, tRNA modifications (namely f5C, hm5C, queuosine and its derivatives, and mcm5U) dynamically change to fine tune codon decoding, usage, and optimality. These changes in codon optimality drive the translation of many pathways and gene sets, such as the ATF4 pathway and selenoproteins, involved in the cellular response to mitochondrial stress. We further examined several of these modifications using targeted approaches. ALKBH1 knockout (KO) abrogated f5C and hm5C levels and led to mitochondrial dysfunction, reduced proliferation, and impacted mRNA translation rates. Our analysis revealed that tRNA queuosine (tRNA-Q) is a master regulator of the mitochondrial stress response. KO of QTRT1 or QTRT2, the enzymes responsible for tRNA-Q synthesis, led to mitochondrial dysfunction, translational dysregulation, and metabolic alterations in mitochondria-related pathways, without altering cellular proliferation. In addition, our analysis revealed that tRNA-Q loss led to a domino effect on various tRNA modifications. Some of these changes could be explained by metabolic profiling. Our analysis also revealed that utilizing serum deprivation or alteration with Queuine supplementation to study tRNA-Q or stress response can introduce various confounding factors by altering many other tRNA modifications. In summary, our data show that tRNA modifications are master regulators of the mitochondrial stress response by driving changes in codon decoding.

## Introduction

Mitochondrial fitness and proper functioning are essential for a myriad of biological processes such as oxidative phosphorylation (OXPHOS), redox balance maintenance, cellular metabolism and signaling(Mick et al., 2020; Winter et al., 2022). Mitochondrial dysfunction is associated with a wide range of pathological conditions such as diabetes, aging, neurodegenerative diseases, and innate errors of metabolism(Mick et al., 2020; O’Malley et al., 2020; To et al., 2019; Wang et al., 2023). However, mitochondria do not possess the machinery to respond to electron transport chain (ETC) induced, and other forms of, mitochondrial dysfunction(Winter et al., 2022). Thus, in response to mitochondrial stress, the signal is relayed via the OMAI1-DELE1-HRI pathway in order to activate ATF4 and the integrated stress response (ISR)(Fessler et al., 2020; Guo et al., 2020). In addition to this, it became understood that the cellular responses and buffering mechanisms towards ETC inhibitors that act on different respiratory complexes are highly nuanced and diverse(Mick et al., 2020; To et al., 2019), a testament to the complexity of mitochondrial biology. While most of the previous work has focused on how the mitochondrial stress signal is relayed to the cytosol, and the consequent events leading to ISR and transcriptional changes, less work has been done in understanding how the cells translationally respond to mitochondrial stress. During oxidative stress, mRNA translation defects and dynamic changes in translation occur that can shape the cellular proteome(Ling and Söll, 2010). In addition, several translational phenomena that orchestrate how the cells respond to oxidative stress in order to shift the proteome towards generating antioxidant proteins are activated, one of the most important is the changes in the tRNA epitranscriptome(Chan et al., 2012; Chionh et al., 2016; Huber et al., 2022; Torrent et al., 2018).

The tRNA epitranscriptome, which refers herein to tRNA modifications, tRNA expression, tRNA derived small RNA fragments (tsRNAs), and tRNA aminoacylation, has emerged as a critical regulator of codon biased translation and maintenance of cellular proteostasis(Ando et al., 2023; Suzuki, 2021). During oxidative stress, tRNA modifications dynamically alter codon optimality and usage in order to promote translation of mRNAs encoding stress response genes and pathways(Chan et al., 2012; Huber et al., 2022). tRNA expression levels also change to achieve the same effect(Torrent et al., 2018). In addition to these mechanisms, tsRNAs are generated to induce translation repression and to interact with various RNA binding proteins (RBPs) to modulate the cellular response to stress(Rashad et al., 2020a; Rashad et al., 2020b; Rashad et al., 2021; Yu et al., 2021). Combined, these findings point to the important regulatory role of the tRNA epitranscriptome in modulating oxidative stress at various levels.

tRNA modifications, the focus of the work presented here, have been the subject of increasing interest in recent years. Aberrations in tRNA modifications were shown to impact diseases such as cancers(Dedon and Begley, 2022; Rosselló-Tortella et al., 2020), diabetes(Matsumura et al., 2023), and neurodegenerative diseases(Bento-Abreu et al., 2018). Anti-codon modifications specifically have been studied for their role in dictating codon decoding and optimality(Suzuki, 2021). For example, tissue specific patterns of tRNA modifications enrichment were shown to determine the codon usage, optimality, and decoding speed across these tissues(Ando et al., 2023). Dynamic shifts in tRNA modifications, such as queuosine, during oxidative stress were also shown to alter the codon recognition to drive the stress response to arsenite stress(Huber et al., 2022). However, despite these advances in our understanding of tRNA modifications, and the new tools available for their probing, there are gaps in understanding the specific regulatory processes governing diverse stress response pathways from the translational point of view. In particular, despite our knowledge of the importance of different mitochondrial tRNA modifications in regulating mitochondrial function(Asano et al., 2018; Suzuki et al., 2020), there are no links between mitochondrial stress response and tRNA modifications mediated translational adaptations.

In this work, we investigated whether tRNA modifications could dynamically change in response to mitochondrial stress induction. Our analysis revealed that mitochondrial stress induced strong translation dysregulation evident by a disconnect between transcriptional and translational changes in the cells. Further, the translational program was governed by a set of tRNA modifications changes that altered the codon decoding patterns to drive the stress response pathways. We further validated the biological function of these modifications, namely queuosine (tRNA-Q) and f5C and hm5C, and their role in maintaining mitochondrial stress and function. Our results reveal a central role of tRNA modifications in regulating the translational response to mitochondrial stress and function that was not previously explored, placing tRNA biology in the heart of mitochondrial diseases.

## Results

### Induction of mitochondrial stress leads to translational stress

Mitochondrial stress was induced via pharmacological agents to inhibit the respiratory complexes I (via Rotenone (Rot)) and III (using Antimycin A (AntiM)), or via ROS induced mitochondrial dysfunction (via sodium metaArsenite (AS)) [Figure 1a]. We adjusted the cell viability to be relatively comparable between stresses [Figure 1b]. Despite comparable cell death, arsenite led to much stronger translational repression compared to antimycin, while rotenone did not lead to significant translational repression as evidenced by a puromycin incorporation assay [Figure 1c]. Further, stress granules (SG) were only observable after arsenite stress, while only a negligible percentage of cells exposed to rotenone had evident SG (<1%). Antimycin did not induce SG assembly. We further investigated the markers for translational stress responses such as the Integrated stress response (ISR) and the ribotoxic stress response (RSR). Eukaryotic translation initiation factor 2α (eIF2α), a marker for ISR, showed increased phosphorylation significantly only after Arsenite stress [Figure 1e]. P38, a marker for RSR(Wu et al., 2020), also showed increased phosphorylation after Arsenite stress [Figure 1e]. The P70 S6 Kinase (P70S6K), which is downstream from mTOR and plays a role in RSR and in promoting mRNA translation(Snieckute et al., 2022), showed modest reduction in its protein levels. However, the phosphorylation of P70S6K was strongly downregulated after mitochondrial ETC inhibition (5-fold downregulation) [Figure 1e]. On the other hand, arsenite induced modest changes in the phosphorylation of P70S6K. Collectively, these results indicate that the regulation of translation and the induction of stress-associated translation repression is unique in ETC inhibition induced mitochondrial stresses compared to arsenite, which induces secondary mitochondrial dysfunction via general reactive oxygen species (ROS) induction.

**Figure 1:**
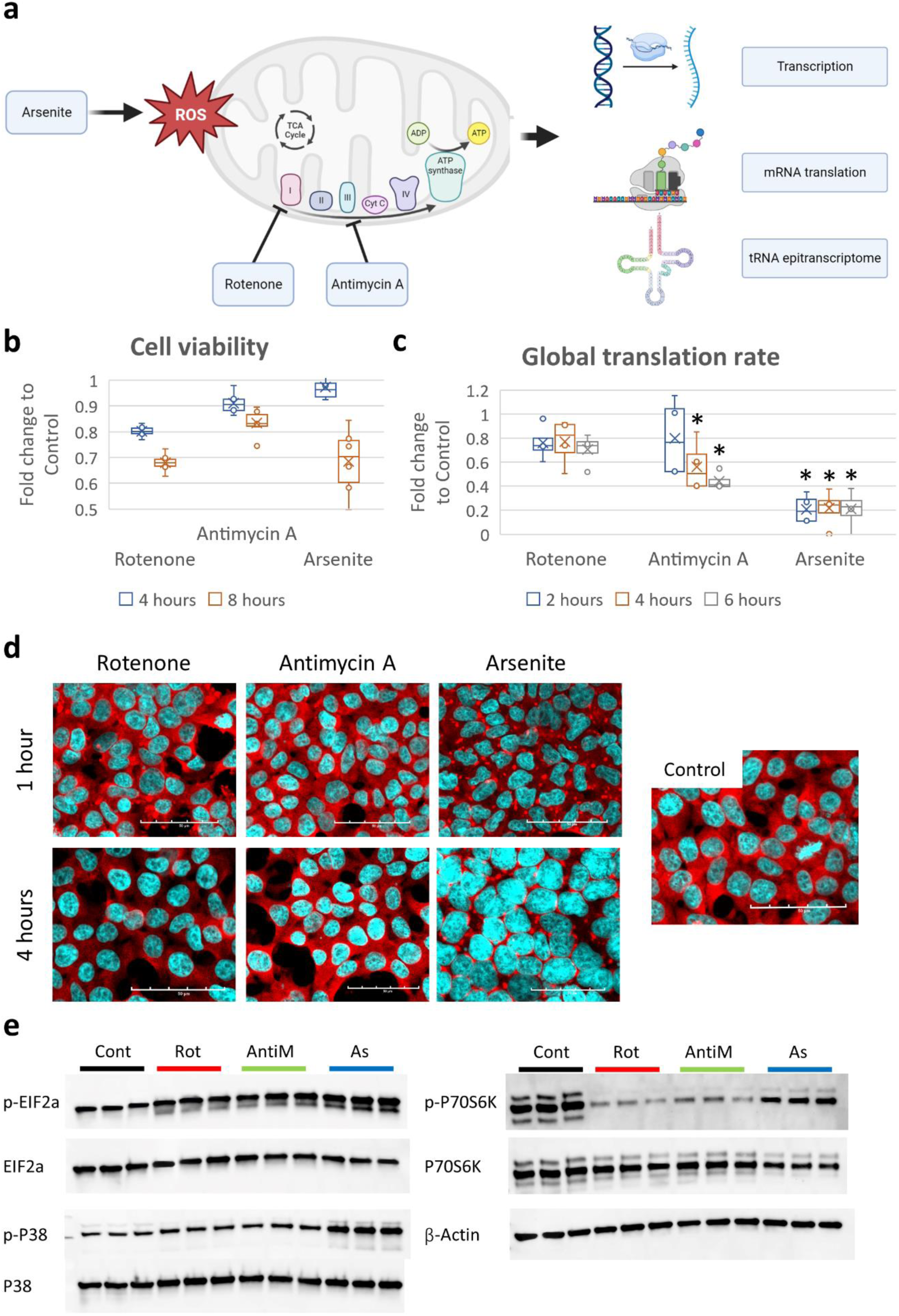
Induction of mitochondrial stress. **a:** Schematic of the study analysis. **b:** Cell viability analysis via MTT assay after exposure to 10μM Rotenone (Rot), 10μg/ml Antimycin A (AntiM), or 100μM sodium metaArsenite (As). **c:** Puromycin incorporation assay after stress exposure. **d:** Staining with anti-G3BP1 (in red) for analysis of stress granules (SG) assembly after stress. **e:** Western blot analysis of ISR and RSR markers after 8 hours of stress.

### Translation and transcription are decoupled during oxidative & mitochondrial stress

To understand how cells transcriptionally and translationally respond to each of these stresses, we conducted RNA-seq, and Ribo-seq. This yielded 3 datasets per stressor (Gene expression, translation efficiency (TE), and mRNA translation levels). At the level of the gene expression [Figure 2a-c], it was apparent that the transcriptional response to arsenite was drastically different from rotenone or antimycin [Figure 2d, Supplementary figure 1a]. Gene ontology biological processes (GOBP) activation matrix further revealed distinct pathways being activated or repressed in each stress [Supplementary figure 1b]. For example, unfolded protein response (UPR) and endoplasmic reticulum stress response (ER-stress) were activated in arsenite, while histone related pathways were activated in Rotenone. The same patterns were observable at the level of translation (i.e., Ribo-seq datasets) [Supplementary figure 1c-e], where arsenite stress showed different clustering pattern compared to respiratory complex inhibitors [Supplementary figure 1f]. GOBP activation matrix revealed distinct pathway signatures pertaining to each stressor [Supplementary figure 1g]. For example, various rRNA and translational pathways were downregulated in Antimycin stress. To further understand the complex relations and interactions between different stressors at multiple levels, we conducted Pearson’s correlation coefficient analysis on the full 9 datasets included in this analysis [Figure 2e]. First, we compared each level of analysis between different stresses. It was apparent that at different levels of analysis, rotenone and antimycin stress responses correlated much better than with arsenite stress response. Further, in each stress, RNA-seq data had low correlation with Ribo-seq data, indicating a divergence between transcription and translation during oxidative stress. However, the correlation between RNA-seq and Ribo-seq was highest in arsenite (*R =* 0.45), while it was < 0.4 in rotenone and antimycin. TE had good agreement with Ribo-seq (*R* > 0.7 in all stresses). Next, we examined the Ribo-seq datasets for translational changes during stress exposure. Metagene distribution plots at the coding sequence (CDS) start site revealed translational changes occurring in antimycin and arsenite stresses only, and not in rotenone [Supplementary figure 2a-i]. These changes were mostly evident at the translational start sites [Supplementary figure 2d-f]. These changes were also more pronounced in arsenite than in antimycin, much in agreement with the puromycin incorporation assay [Figure 1c]. We also examined ribosome pause sites in the E, P, and A sites [Figure 2f]. The A site ribosome pausing revealed a high degree of agreement between different stresses in terms of codons that are stalled vs codons that were more ready translated, with some inter-stress variations observed in some codons.

**Figure 2:**
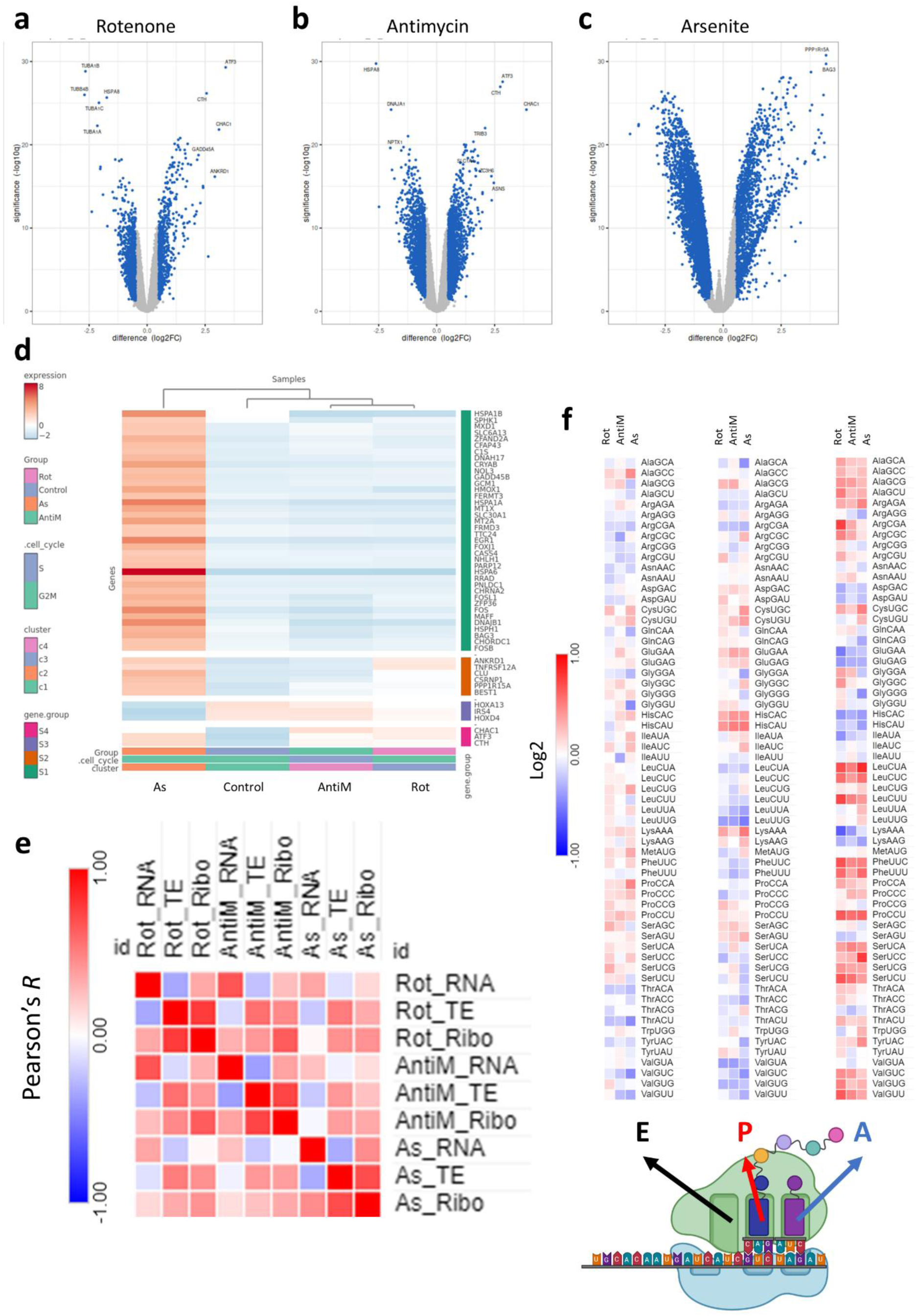
Mitochondrial stress induces transcriptional and translational dysregulation. **a-c:** Volcano plots for differentially expressed genes (DEGs) in the RNA-seq analysis after 8 hours of stress exposure. **d:** Cluster heatmap analysis of the RNA-seq datasets. **e:** Ribosome dwelling times after stress exposure across different ribosome sites. **f:** Pearson’s correlation analysis between stresses across different datasets.

### tRNA modifications dynamically change in response to mitochondrial stressors

The observation of changes at the translational level that were independent from transcription, as well as the changes in codon decoding efficiency by the ribosomes led us to examine tRNA modifications, as an important regulator of translation via altering codon usage and optimality during stress(Suzuki, 2021). We analyzed the dynamic changes in small RNA (>90% tRNA) modifications levels following exposure to the stressors in question using high throughput liquid chromatography tandem mass spectrometry (LC-MS/MS) [Figure 3a, Supplementary table 1]. Of the 44 modifications detected in our analysis, 5-formylcytidine (f5C), 5-hydroxymethylcytidine (hm5C), queuosine (Q) and its derivatives galactosyl-queuosine (galQ) and mannosyl-queuosine (manQ), and 5-methoxycarbonylmethyluridine (mcm5U) changed in a statistically significant manner [Figure 3a]. f5C, hm5C, and mcm5U were significant only in arsenite stress, while Q, manQ, and galQ were upregulated in all stresses. f5C and hm5C were downregulated after 8 hours of exposure to arsenite, while the other modifications were upregulated. We further explored the Ribo-seq datasets for the expression patterns of known tRNA and small RNA modifying enzymes, curated from the literature and from reference(Suzuki, 2021) [Figure 3b]. We did not observe overt changes in the tRNA modifying enzymes responsible for the detected modifications that can explain the changes observed.

**Figure 3:**
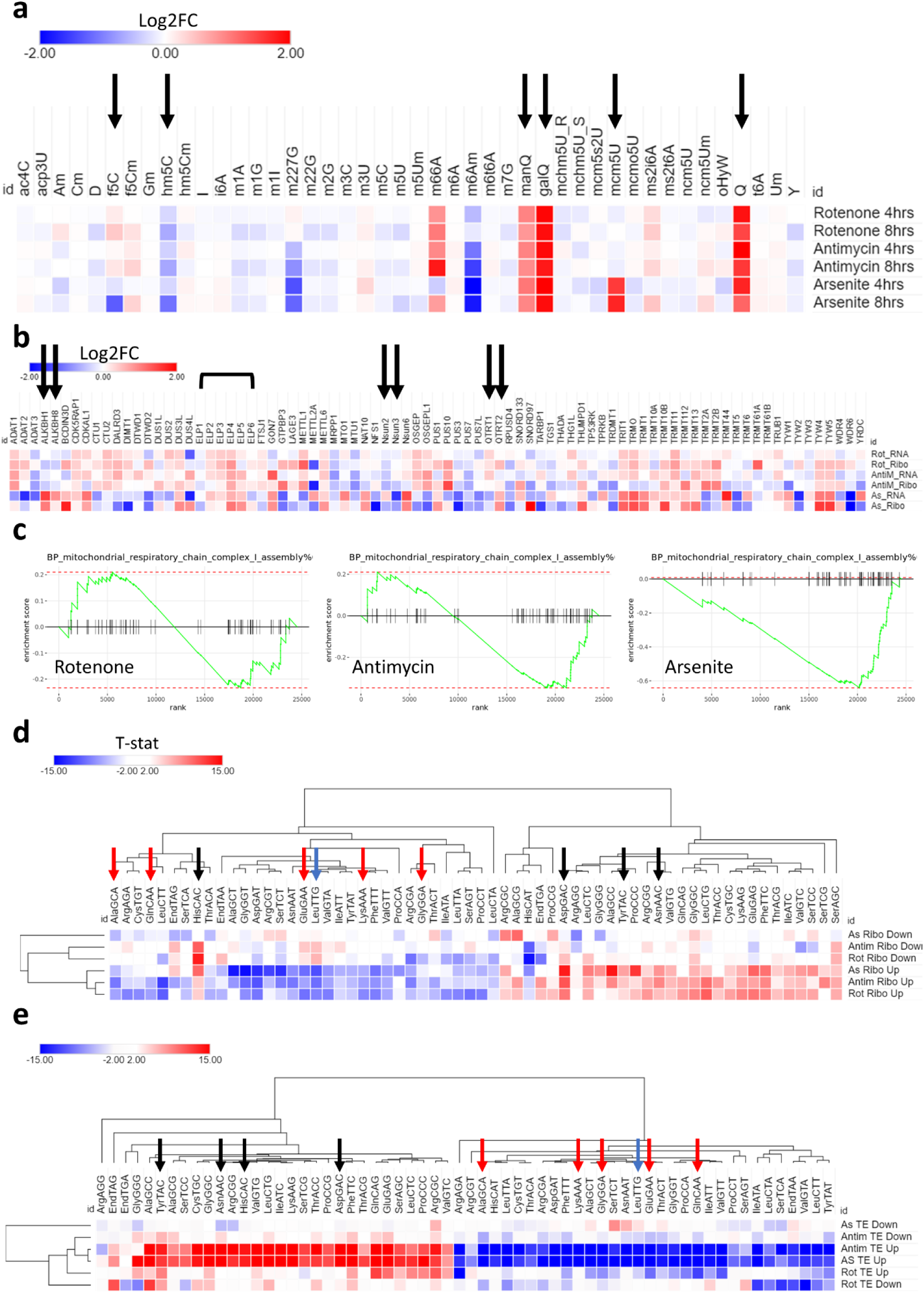
tRNA modifications drive translational changes after mitochondrial stress. **a:** Analysis of tRNA modifications after exposure to 10μM Rotenone, 10μg/ml Antimycin A, or 100μM Arsenite. Black arrow indicates fold change > 1.5 and *p* < 0.05. Data is normalized to controls. **b:** Analysis of all tRNA modifying enzymes from the Ribo-seq and RNA-seq datasets. Black arrows indicate enzymes related to significant modifications in **a**. **c:** GOBP pathway analysis of mitochondrial respiratory chain complex I assembly in all stresses. **d:** Isoacceptors codon frequency analysis of the Ribo-seq datasets in all stresses. **e:** Isoacceptors codon frequency analysis of the translational efficiency (TE) datasets in all stresses.

We directed our attention to the potential biologic or translational impact of the modifications that we observed to be significantly dysregulated. First, we examined f5C and hm5C [Supplementary figure 3a]. Both were downregulated in response to stress. f5C was downregulated only in arsenite after 8 hours of stress, while hm5C was downregulated in all stresses after 8 hours [albeit less than the arbitrary cutoff of 1.5-fold after rotenone and antimycin exposure] [Supplementary figure 3b-c]. f5C is present in mitochondrial methionine tRNA (mt-tRNA-Met) at the wobble position (position 34) and is essential for proper mitochondrial translation(Delaunay et al., 2022; Kawarada et al., 2017). f5C is synthesized from m5C (5-methylcytidine) via ALKBH1, while m5C is synthesized from Cytidine (C) via NSUN3(Kawarada et al., 2017). m5C itself did not show significant changes after stress. Indeed, dilution of any m5C changes in the mitochondrial tRNA-Met by the abundance of m5C in the cytosolic and other mitochondrial tRNAs would obscure any changes occurring in this one tRNA unless a targeted analysis was performed(Kawarada et al., 2017; Suzuki, 2021; Suzuki et al., 2020). hm5C is present in cytosolic tRNA leucine (tRNA-Leu^CAA^) at the wobble position and is synthesized in a multi-step process than involves NSUN2 converting C to m5C, followed by ALKBH1 converting m5C into hm5C and further to f5C(Kawarada et al., 2017). Again, changes in m5C levels cannot inform the changes in hm5C as m5C is abundant in tRNAs in positions other than the anticodon(Suzuki, 2021). There are multiple layers of evidence linking f5C and hm5C to mitochondrial bioenergetics and translation as well as stress response via modulating respiratory complex translation and mitochondrial decoding of transcripts(Delaunay et al., 2022; Kawarada et al., 2017; Rashad et al., 2022). To that end, we examined the expression of mitochondrial respiratory chain complexes and related pathways in our Ribo-seq datasets. GOBP pathway analysis revealed downregulation of respiratory chain complex I assembly pathway only in arsenite stress [Figure 3c]. Further analysis of genes related to respiratory complexes I to V detected in the Ribo-seq analysis revealed a strong downregulation of genes linked to respiratory complexes I, II, III, and V in arsenite [Supplementary figure 3g].

Respiratory complex IV did not show the same level of downregulation as with the other complexes, and the downregulation in Rotenone or antimycin was much less profound. In summary, the analysis shows that the changes in mitochondrial OXPHOS genes in response to arsenite stress could be driven by changes in f5C and hm5C tRNA modifications.

### tRNA modifications drive the mitochondrial stress translational response via altering codon usage and bias

Given the regulatory role of tRNA modifications in codon biased mRNA translation and codon optimality changes, and that the observed significantly changing modifications (mcm5U, Q, f5C, and hm5C) are linked to wobble positions in the anti-codon and their levels are essential in decoding the cognate codon, we examined the codon usage and biases (isoacceptors frequencies(Rashad et al., 2022)) and ribosome stalling at the A-sites(Liu et al., 2020) for known codons linked to the highlighted modifications. Our goal was to study the effect of these modifications on the translational phenomena observed, and how much each modification contributed to the system in question.

First, we examined the ribosome pausing at A-positions [Figure 2e and supplementary figure 5]. In all stresses, we observed changes in ribosome dwelling times at A-sites, indicating changes in codon decoding capacity. There was an evident enhanced translation at the queuosine decoded codons (Q-codons; NAY codons(Suzuki, 2021)) to varying degrees, as well as the NAA ending mcm5U-decoded codons. Importantly, the mcm5U decoded codons were better translated in rotenone and antimycin stresses even in the absence of upregulation of mcm5U modification. Leu-TTG (UUG) codon, which is decoded by hm5C, did not show changes in ribosome dwelling in any stress, including Arsenite.

Next, we analyzed the isoacceptors frequencies (i.e., codon usage and bias), which is defined as the ratio of expression of each codon to its synonymous codons in a given mRNA(Rashad et al., 2022), in the top differentially expressed genes (cut-off: Log2FC >2 or <-2, FDR <0.05) in the Ribo-seq [Figure 3d] and TE [Figure 3e] datasets. In the Ribo-seq dataset [Figure 3d], there was a strong, yet imperfect, A/T versus G/C ending codons bias. The bias in codon usage was stronger in the upregulated (i.e., up-translated) genes than in the downregulated genes (evident by the higher differences in isoacceptors frequencies values). Further, mcm5U codons were less optimal in the upregulated genes in all stresses, despite mcm5U being upregulated after arsenite stress. hm5C codon, Leu^TTG^ was suboptimal in the upregulated genes, in line with the downregulation of the tRNA modification. C-ending Q-codons (NAC codons) were all optimal in the upregulated genes, except for His^CAC^. His^CAC^ also showed increased ribosome pausing in the AS dataset [Supplementary figure 5c]. In the TE dataset [Figure 3e], we again observed the same A/T versus G/C bias. We also observed that upregulated genes (i.e., genes with enhanced translational efficiency) in the Arsenite and Antimycin datasets were the most biased, while Rotenone had a somewhat weaker impact. In summary, our analysis revealed that all the C-ending Q-codons were associated with enhanced translation of mRNAs, while mcm5U and hm5C codons were associated with reduced translational efficiency.

To further reveal how codon patterns correlate between datasets, we conducted correlation analysis for each stress between different methods [Supplementary figure 6a]. In rotenone stress, Ribo-seq showed low correlation between the up and downregulated genes, which was again evident in the translation efficiency dataset, indicating codon biased translation. Genes with enhanced translation efficiency correlated well with the upregulated genes in Ribo-seq, but not with downregulated genes, at the codon level, indicating the changes in TE, driven by tRNA modifications, are a strong driver of translation. In the antimycin dataset, there was a no correlation between the up and down regulated genes in Ribo-seq, as well as with the downregulated genes in Ribo-seq and TE. The TE dataset did not show distinct dichotomy between up and downregulated genes. Both datasets, on the other hand, correlated well with the more translated genes. In the Arsenite dataset, a different pattern of correlation was evident. While up and down regulated genes in Ribo-seq did not correlate, up and down regulated genes in TE had negative correlation. To further understand these patterns, we examined the cognate codons for the modified tRNAs in the TE dataset once again [Figure 3e]. It is apparent that the mcm5U and hm5C codons, both modifications are changing only in arsenite, are contributing to these patterns to an extent. Collectively, these data indicate that tRNA modifications changes are a driving force behind mRNA translation and translation efficiency via altering codon decoding and optimality. We next analyzed the correlation between each stress at the level of each dataset [Supplementary figure 6a]. It was clear that at the level of translation there was a good correlation between the upregulated genes in all stresses. However, at the TE level, antimycin and rotenone stresses did not show good correlation [Supplementary figure 6a].

Next, we directed our attention to analyzing the impact of codon usage and biases and tRNA modifications changes on the expression of various pathways and gene groups after stress induction. First, we analyzed the codon usage and patterns in mitochondrial respiratory complex genes [Supplementary figure 6b]. Above, we showed that arsenite induced a strong downregulation of all respiratory complex genes except for complex IV [Supplementary figure 3d]. Analysis of codon usage and bias for each respiratory complex gene group revealed a strong enrichment of the hm5C codon, Leu^TTG^, in all complexes except complex IV, which had the lowest enrichment. In addition, there were no specific patterns of enrichment of mcm5U or Q-codons, indicating that the changes in translation levels of respiratory complexes are mostly driven by hm5C.

We next examined selenoproteins’ codon usage and biases. Selenoproteins are important in the context of oxidative stress(Alim et al., 2019). Codon bias analysis revealed a clustering pattern that divides selenoproteins into 2 main groups, one that is mostly G/C biased, and the other is mostly A/T biased [Supplementary figure 6c]. In consequence, the enrichment of C-ending queuosine codons and mcm5U codons, which are A ending, showed different patterns of enrichment in both groups. To further evaluate whether this G/C vs A/T bias can inform gene expression of selenoproteins, we extracted the expression data of selenoproteins from the Ribo-seq datasets in the three stresses examined. Only in arsenite did we observe strong changes in selenoproteins translation [Supplementary figure 6d]. In addition, the upregulation of multiple selenoproteins was skewed towards the G/C-ending codon biased ones. Correlation analysis also revealed that the expression of selenoproteins in arsenite is unique compared to rotenone and antimycin [Supplementary figure 6e]. To further clarify the contribution of codons to the expression of selenoproteins, we performed PLS-DA clustering analysis after dividing the proteins into up and downregulated as per the clustering pattern observed in Supplementary figure 6c.

PLS-DA revealed good divergence between up and down regulated selenoproteins in terms of codon usage and bias patterns [Supplementary figure 7a-b]. In addition, analysis of the contribution of each codon to the PLS-DA component 1, which explains most of the variance observed, revealed that the mcm5U codons, and His^CAC^ codon, all contributed to the downregulated genes, while their synonymous codons and other C-ending Q-codons contributed to the upregulated genes [Supplementary figure 7c]. This pattern was also evident when we conducted total codon count analysis, which examines the codon composition across the entire mRNA, contrary to isoacceptors frequency which examines codon bias and optimality of synonymous codons(Ando et al., 2023) [Supplementary figure 7d]. All in all, this indicate that the upregulation of select selenoproteins during Arsenite stress, and their consequent effect on stress response, at the translational level is mainly driven by C-ending Q-codons.

We further expanded our codon analysis to examine the expression of various pathways in the Ribo-seq datasets. Using pre-ranked gene set expression analysis (GSEA) [Supplementary table 2] we extracted the information of the top 5 up and down enriched pathways in the three stresses tested herein. Codon usage and bias revealed subsets of these pathways that are strongly A/T versus G/C biased, and in turn their patterns of enrichment of mcm5U, hm5C, and Q-codons followed accordingly [Supplementary figure 7e]. However, the initial clustering pattern analysis showed some overlap between some up and down enriched pathways, while others had clear correlation with the expression of the corresponding codons and modifications. To evaluate the contribution of the codon optimality patterns to pathway enrichment, we conducted PLS-DA analysis on the up and down enriched pathways. PLS-DA revealed different codon enrichment patterns and contribution to pathway expression [Supplementary figure 7f-g]. Analysis of each codon contribution revealed that C-ending Q-codons contributed to the upregulated pathways, while mcm5U and hm5C codons contributed mostly to the downregulated pathways [Supplementary figure 7h]. We further analyzed the ATF4 pathway, which plays a major role in mitochondrial stress response(Quirós et al., 2017). Given its importance, we asked whether the ATF4 pathways, which is transcriptionally activated during mitochondrial and endoplasmic unfolded protein responses, is optimized for proper translation via the codon sequences of its genes. This question is important in the context of the transcriptional-translational mismatch observed in the stresses tested herein [Figure 2f]. Indeed, ATF4 pathway was upregulated at the transcriptional and translational levels in all stresses tested [Supplementary figure 8a-c]. Next, we looked for genes which are transcriptionally activated via ATF4 in the TRRUST database(Han et al., 2018) and analyzed their codon usage and biases. At the gene level, we observed a large cluster of ATF4 regulated genes that were G/C biased [Supplementary figure 8d]. Indeed, some of the ATF4 genes were also A/T biased in this analysis. However, examining the ATF4 regulated genes as a group, as done for pathway analysis above, we observed a strong global G/C bias vs A/T ending codons [Supplementary figure 8e]. The analysis of the expression of ATF4 genes in our Ribo-seq data revealed a strong correlation between the expression of individual genes and their G/C ending codon preferences [Supplementary figure 8f]. Collectively, our data indicates that the pathway enriched patterns observed at the level of translation during stress are driven, to a large extent by tRNA modifications mediated codon decoding and shifts in codon optimality.

### f5C and hm5C modifications impact mitochondrial respiration and function

Given our findings regarding tRNA modifications, we directed our attention to the important modifications identified in our analysis. f5C and hm5C both share a common synthesis pathway(Kawarada et al., 2017) [Figure 4a]. Given that ALKBH1 is a common enzyme in the synthesis of both modifications(Kawarada et al., 2017), we generated two CRISPR knockout (KO) cell line clones from HEK293T to study the role of these modifications in mitochondrial and oxidative stress [Figure 4b]. It’s notable that ALKBH1 protein levels did not change significantly after 8 hours of stress exposure [Figure 4c], nor did its upstream enzymes (NSUN2 and NSUN3) in this pathway(Kawarada et al., 2017) show changes in their translational levels in the Ribo-seq data [Figure 3b]. Using LC-MS/MS, we validated that ALKBH1 KO abrogated f5C and hm5C and their downstream modifications f5Cm and hm5Cm [Figure 4d and e. Supplementary figure 9a-c, Supplementary table 3]. While ALKBH1 was reported to act as a 1-methyladenosine (m1A) demethylase previously(Arguello et al., 2022; Liu et al., 2016; Rashad et al., 2020a), we did not observe an effect of ALKBH1 KO on m1A levels. We also observed that f5C was not completely abolished in the KO clones, with clone 1 showing more reduction in f5C levels than clone 2 [supplementary figure 9a]. In addition to this effect on known ALKBH1 substrates, we observed indirect effects of ALKBH1 KO on other modifications [Figure 4d]. Namely, there was a significant upregulation of mcm5U in both clones and upregulation of queuosine in clone 2.

**Figure 4:**
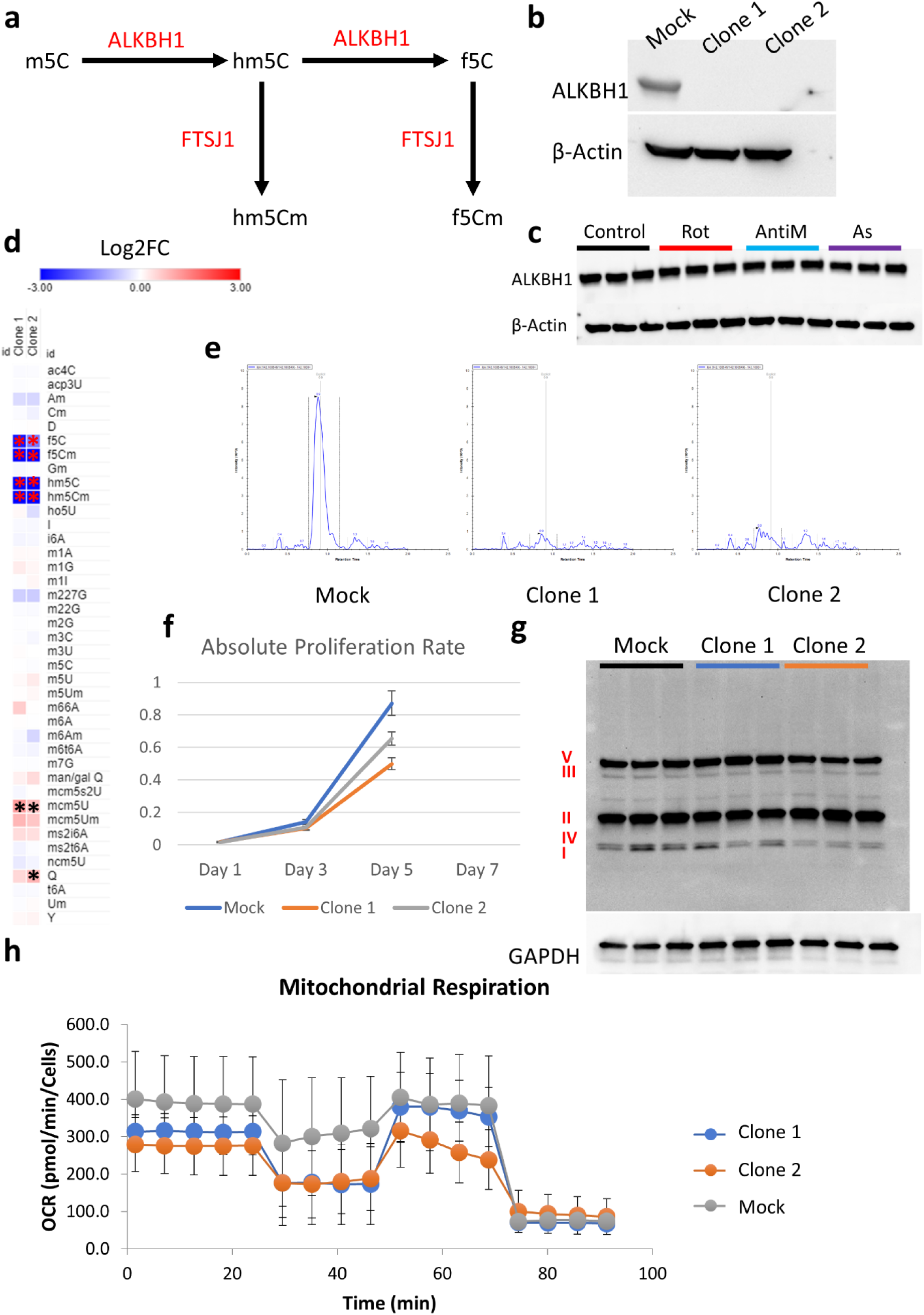
ALKBH1 regulates mitochondrial function through its dioxygenase activity. **a:** Schematic for the synthesis pathway of hm5C and f5C modifications and the responsible enzymes. **b:** Validation of ALKBH1 knockout (KO) in the two generated clones. **c:** Analysis of ALKBH1 expression after 8 hours of stress exposure. **d:** LCMS/MS analysis of tRNA modifications after ALKBH1 KO. Data is normalized to Mock. Asterisk indicates fold change > 1.5 and *p* < 0.05. **e:** Analysis of hm5C peaks showing the complete depletion of hm5C after ALKBH1 KO. **f:** Cell proliferation after ALKBH1 KO. **g:** Analysis of OXPHOS related proteins in all 5 complexes using western blot. **h:** Mitochondrial respiration analysis using Seahorse.

ALKBH1 KO cells grew slower than their Mock KO counterparts [Figure 4g], in contrast to what was previously observed with ALKBH1 shRNA KD(Liu et al., 2016) as well as our observations in ALKBH1 overexpressing glioma cell line(Rashad et al., 2022). It was also noted that clone 1 grew slower than clone 2, in line with the stronger reduction in f5C levels observed in clone 1. We next exposed the KO clones to a stress panel encompassing mitochondrial respiratory complexes inhibitors I to V [Rotenone; I, TTFA; II, Antimycin; III, Potassium cyanide (KCN); IV, and oligomycin; V] in addition to arsenite. We did not observe significant influence of ALKBH1 KO on cell viability after stress, except for a protective effect against TTFA in clone 1 [Supplementary figure 9d]. However, the results showed high variability across different iterations and was not replicated across clones, and so a definitive conclusion could not be achieved regarding this effect. Puromycin assay showed reduced global translation rates after ALKBH1 KO which were more pronounced in clone 2 [supplementary figure 10a]. We further analyzed the markers of ISR and RSR after ALKBH1 KO [Supplementary figure 10b-f]. ALKBH1 KO led to modest downregulation of eIF2α and significant downregulation of its phosphorylated form [Supplementary figure 10b-c]. Phosphorylated P38 was also downregulated to varying degrees in both ALKBH1 KO clones [Supplementary figure d-e]. On the other hand, phosphorylated P70S6K was extensively upregulated in both clones [Supplementary figure 10d & f]. We next validated the impact of ALKBH1 KO on mitochondrial respiratory complexes and activity, and as previously reported(Kawarada et al., 2017), we observed an impact on respiratory complex I levels specifically [Figure 4g]. Seahorse analysis revealed dysfunctional mitochondrial respiration and bioenergetics in both clones [Figure 4h, Supplementary figure 11a-h].

### tRNA-Q depletion impacts global tRNA epitranscriptome and induce translational stress

LC-MS/MS and codon analysis data strongly pointed to tRNA queuosine (tRNA-Q) as a major regulator of the translational response to mitochondrial stress and dysfunction. tRNA-Q is synthesized by tRNA-guanine trans-glycosylase (TGT) complex, which is composed of 2 enzymes: QTRT1 and QTRT2(Behrens et al., 2018; Johannsson et al., 2018; Sebastiani et al., 2022; Sievers et al., 2021). The TGT complex utilizes Queuine to convert Guanosine into Queosine in GUN anticodon, which is essential for the expanding the codon recognition from NAC codons to NAY codons(Dixit et al., 2021; Huber et al., 2022; Sebastiani et al., 2022; Suzuki, 2021; Tuorto et al., 2018) [Figure 5a]. However, QTRT1 and QTRT2 protein levels did not change after 8 hours of stress [Supplementary figure 12a], contrary to the strong upregulation of queuosine levels in tRNA.

**Figure 5:**
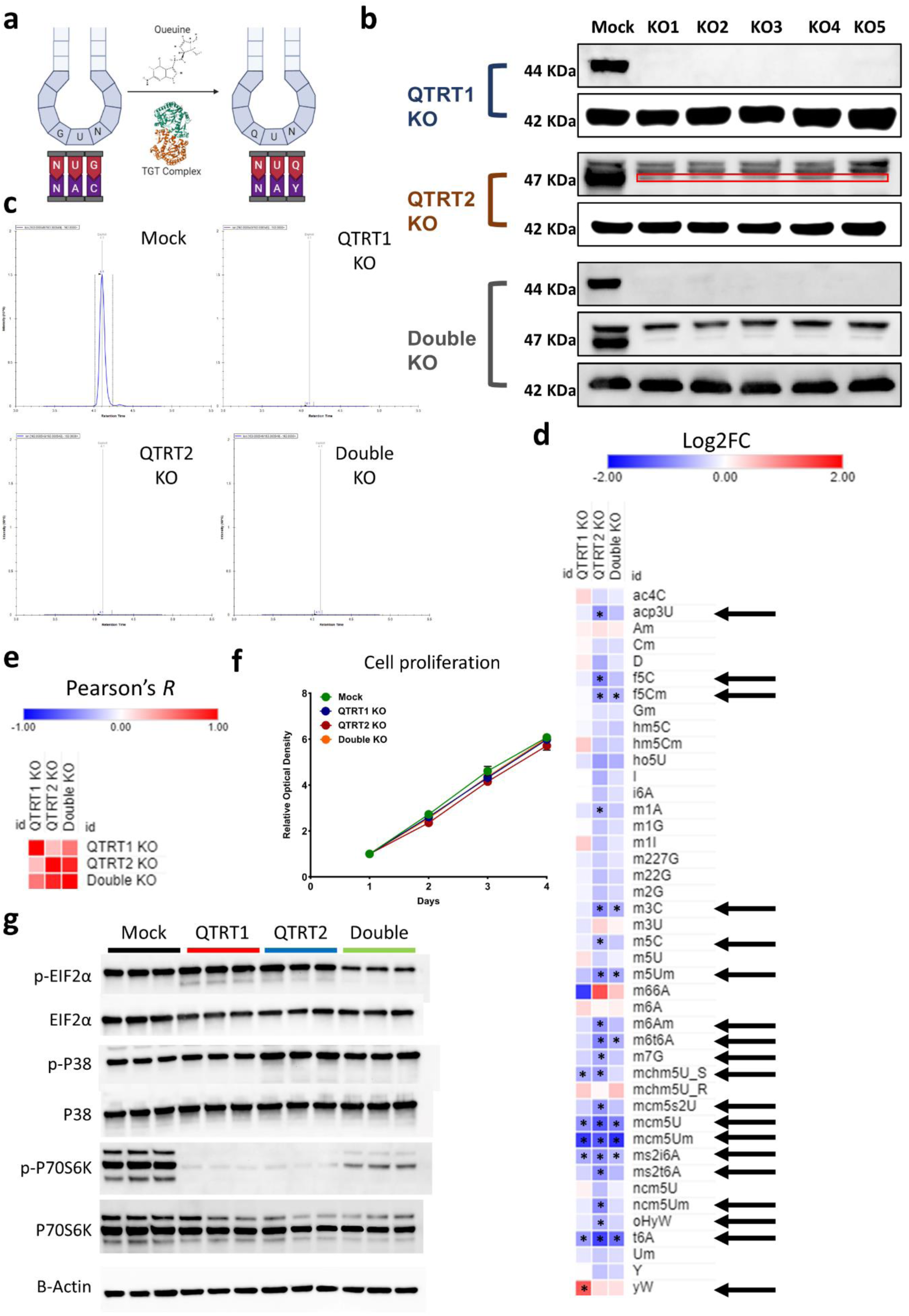
tRNA-Q loss induces translational stress. **a:** Schematic of tRNA-Q mediated codon recognition. **b:** Validation of QTRT1, QTRT2, and double KO using western blots. **c:** Validation of tRNA-Q peak loss using LC-MS/MS. **d:** Analysis of the impact of tRNA-Q loss on other RNA modifications in the three KO cell lines. Arrows: significant on ANOVA analysis. Asterisk: fold change > 1.5, *p* < 0.05. Data normalized to Mock. **e:** Pearson’s correlation analysis using data from **d**. **f:** Proliferation rates of the three KO cell lines. **g:** Analysis of markers of ISR and RSR using western blotting.

To elucidate the impact of tRNA-Q on translation and mitochondrial dysfunction, we generated 3 cell lines, in addition to the Mock cell line, using CRISPR knockout: QTRT1 KO, QTRT2 KO, and a double KO cell line. We validated the KO using western blotting and T7 endonuclease genomic cleavage assay [Figure 5b and supplementary figure 12b]. We also validated the loss of tRNA-Q using LC-MS, which revealed the loss of Q, manQ, and galQ in all KO cell lines [Figure 5c and supplementary figure 12c and d]. We also analyzed other tRNA modifications during our validation in order to identify other indirect effects downstream of tRNA-Q loss that can impact translation [Figure 5d, Supplementary table 3]. Our analysis revealed global alterations in tRNA modifications status that could reflect global epitranscriptional and translational aberrations upon tRNA-Q loss. Notably, several modifications that are linked to mitochondrial function such as f5C, f5Cm, and ms2i6A(Schweizer et al., 2017) were downregulated after tRNA-Q depletion. We also analyzed the correlation between the levels of tRNA modifications after QTRT1 and QTRT2 KO and observed a lower correlation between QTRT1 and QTRT2 KO cell lines than expected (*R* = 0.25) [Figure 5e]. Double KO correlated better with QTRT2 KO (*R =* 0.86) than it did with QTRT1 KO (*R* = 0.53). Proliferation analysis revealed no difference in cell growth between tRNA-Q KO cell lines and Mock cells [Figure 5f]. The Puromycin incorporation assay revealed significant translation repression after QTRT1 KO but not after QTRT2 KO, while the double KO cell line had the strongest translation repression effect [Supplementary figure 12e]. To that end, we analyzed the ISR and RSR markers to evaluate the existence of translational stress [Figure 5g]. p-eIF2α was significantly downregulated in the double KO group but not in the QTRT1 or QTRT2 KO only groups. In addition, eIF2α proteins were mildly downregulated in all groups (between 27∼37% downregulation) [Figure 5g and Supplementary figure 12f]. P38 did not show significant changes in its protein levels, however, P38 phosphorylation was significantly upregulated in the QTRT2 KO cells [Figure 5g and Supplementary figure 12f]. Finally, phosphorylated P70S6K (p-P70S6K) was extremely downregulated in all three KO cell lines, akin to what was observed during pharmacological mitochondrial ETC inhibition [Figure 1e, 5g and Supplementary figure 12f].

We further analyzed the formation of stress granules (SGs) using immunofluorescence staining of the core SG protein G3BP1 [Supplementary figure 13a]. We did not observe apparent spontaneous SG assembly in the KO cells. However, staining against EDC4, a marker for P-bodies, revealed an increased number of P-bodies after QTRT2 KO [Supplementary figure 13b].

### tRNA-Q loss induces transcriptional and translational aberrations and alters NAU codon decoding

To understand the global impact of queuosine loss on mRNA expression and translation, we conducted RNA-seq and Ribo-seq on the QTRT1 and QTRT2 KO cells. RNA sequencing revealed strong transcriptional response to QTRT1 and QTRT2 KO [Figure 6a-b]. GOBP analysis revealed the upregulation of various translational and mRNA processing related pathways in both KO cell lines [Figure 6c-d]. At the translational level, we observed fewer changes related to mRNA translation in both cell lines [Figure 6e-f]. However, GOBP analysis revealed various changes the resonates the transcriptional activation such as the enrichment of cytoplasmic translation [Figure 6g-h]. In both KO cell lines, the top enriched pathway in the Ribo-seq datasets was related to mitochondrial ATP synthesis. On the other hand, we observed downregulation of protein refolding pathways after QTRT2 KO, while they were upregulated after QTRT1 KO.

**Figure 6:**
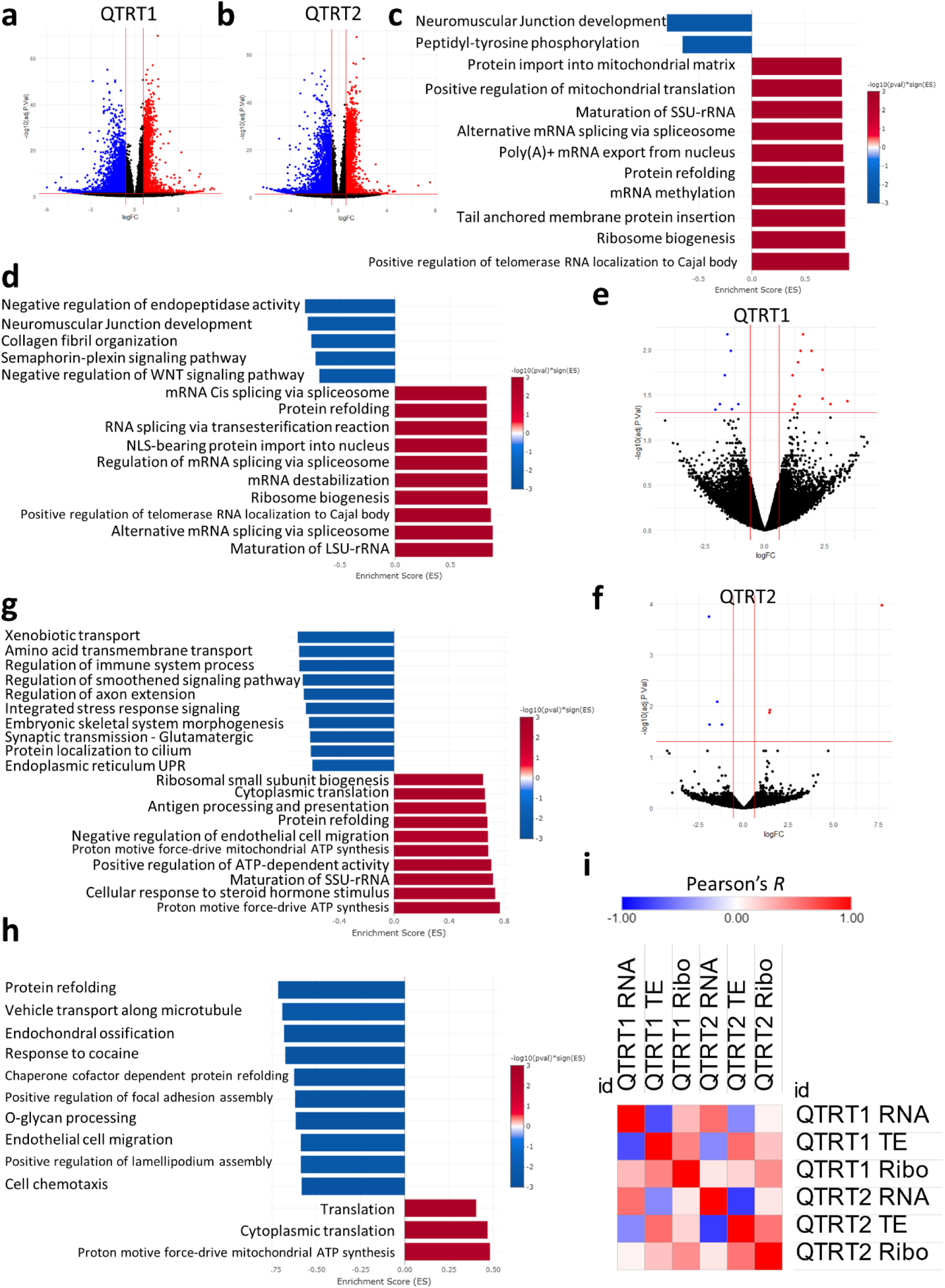
Impact of QTRT1 and QTRT2 KO on mRNA expression and translation. **a-b:** Volcano plots for RNA-seq datasets. **c:** RNA-seq GOBP analysis after QTRT1 KO. **d:** RNA-seq GOBP analysis after QTRT2 KO. **e-f:** Volcano plots for Ribo-seq dataset. **g:** Ribo-seq GOBP analysis after QTRT1 KO. **h:** Ribo-seq GOBP analysis after QTRT2 KO. **i:** Pearson’s correlation analysis between different datasets and cell lines.

Next, we conducted Pearson’s correlation analysis between different datasets, including translational efficiency (TE) and between the two KO cell lines [Figure 6i]. In both cell lines, we observed a strong negative correlation between TE, which can be a proxy for translation initiation(Zhao et al., 2023), and RNA levels (−0.7 in QTRT1 and −0.77 in QTRT2). In addition, we observed very low correlation between RNA levels and RNA translation (i.e., RNA-seq and Ribo-seq) (0.28 in QTRT1 and 0.1 in QTRT2). Both findings can be considered as hallmarks for translational dysregulation. When comparing the two cell lines, we observed a moderate correlation between QTRT1 and QTRT2 KO at the RNA-seq and Ribo-seq datasets levels (0.56 and 0.44 respectively).

In order to explore this observed translational dysregulation, we turned our attention to our Ribo-seq data. First, we observed the distribution of the Ribosome protected fragments (RPFs) across the coding sequence (CDS) [Figure 7a-f]. In both cell lines, there were apparent changes in ribosome occupancy compared to the Mock cells. Further, RPFs tended to accumulate near the start codons rather than the end of the CDS.

**Figure 7:**
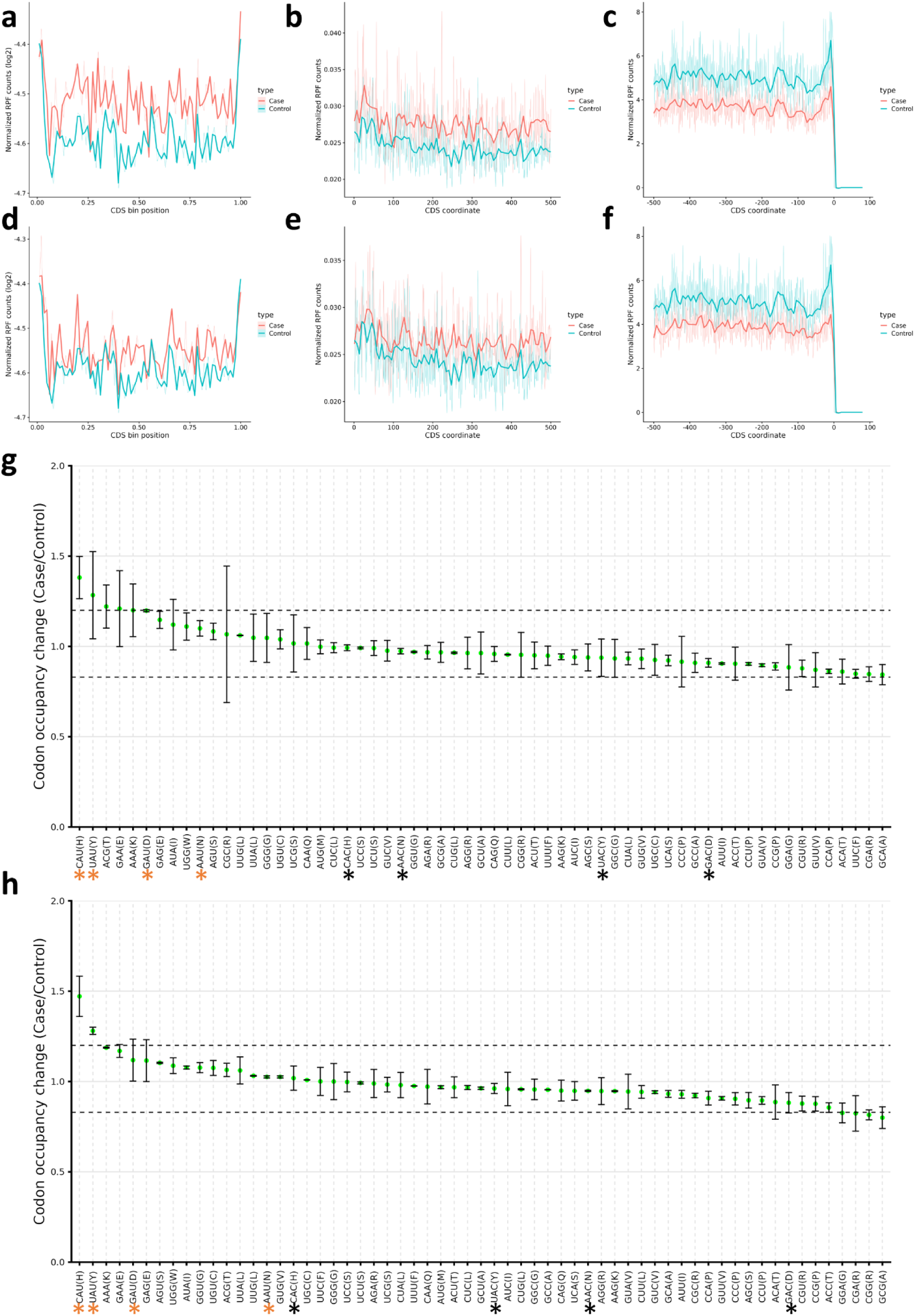
tRNA-Q loss impacts NAU codon decoding. **a-c:** Occupancy metagene plots in QTRT1 KO cells (Across CDS, downstream from start codon, and upstream from end codon respectively). **d-f:** Occupancy metagene plots in QTRT1 KO cells (Across CDS, downstream from start codon, and upstream from end codon respectively). **g:** Ribosome A-site pausing in QTRT1 KO. **h:** Ribosome A-site pausing in QTRT2 KO. Black asterisk: NAC codons, orange asterisk: NAU codons.

We next directed our attention to the codon decoding capacity. A-site pausing revealed significant pausing of NAU Q-codons, while NAC codons decoding was not significantly impacted. These results were consistent between both KO cell lines, with His^CAU^ and Tyr^UAU^ codons being the most impacted [Figure 7g-h]. We further analyzed the isoacceptors and total codon frequencies in both cell lines from the TE and Ribo-seq datasets [Supplementary figure 14]. There was an apparent bias in the codon sequences of translated mRNAs towards His^CAC^ and Tyr^TAC^ (UAC) compared to their synonymous codons [Supplementary figure 14a]. This was not evident when considering total codon frequencies [Supplementary figure 14b]. In addition, the links between codon decoding and TE were not clear [Supplementary figure 14c-d].

### Loss of tRNA-Q leads to mitochondrial dysfunction

Since tRNA-Q modifications exist in the mitochondria and are linked to mitochondrial translation and function(Suzuki, 2021; Suzuki et al., 2020), and given our observation of the enrichment of mitochondrial pathways at the transcriptional and translational levels, we sought to evaluate the mitochondrial function in our KO cell lines. First, we analyzed the expression of different OXPHOS subunits using western blotting, which did not reveal major differences in all three KO cell lines [Supplementary figure 15a-b]. Mito tracker green and red staining did not reveal major changes in mitochondria networks or mitochondrial membrane potential after QTRT1 or 2 KO [Supplementary figure 15c]. We further analyzed the mitochondrial function using seahorse [Supplementary figure 16a-i]. Mitochondrial function in all KO cell lines was reduced compared to the Mock cells. Basal respiration, proton leak, maximal respiration, non-mitochondrial oxygen consumption, and ATP production were all impacted [Supplementary figure 16b-f], while spare respiratory capacity and coupling efficiency were not significantly affected except in QTRT2 KO where the coupling efficiency was increased [Supplementary figure 16g-i]. This data may indicate that the mitochondrial dysfunction observed herein is due to mitochondrial translational dysregulation, rather than changes in nuclear encoded mitochondrial proteins, and the observed changes in the RNA-seq and Ribo-seq datasets were a feedback response to such dysfunction.

### tRNA-Q loss impacts cellular responses to mitochondrial stress

tRNA-Q was previously shown to protect against arsenite induced oxidative stress(Huber et al., 2022), however, previous works have utilized Queuine deprivation as a proxy for tRNA-Q depletion(Huber et al., 2022; Tuorto et al., 2018). Thus, it is unclear whether genetic deletion of the TGT complex will yield similar results. To that end, we designed a stress experiment in which the three KO cell lines were exposed to mitochondrial respiratory complex inhibitors (I-V) in their regular media, in the presence of Queuine, in FBS free media, and in FBS free media supplemented by Queuine [Figure 8a-d, supplementary figure 17]. In regular media, only QTRT1 KO showed an effect which was a protective effect against respiratory complex III inhibition. Addition of Queuine led to lower cell viability after stress across cell lines and stresses and abrogated the protective effect of QTRT1 KO on antimycin stress to some degree. However, Queuine addition led to some modest protection against TTFA and KCN in QTRT1 KO cells. On the other hand, QTRT2 KO cells were the most sensitive to Queuine addition especially with TTFA and antimycin stresses. FBS depletion sensitized all cell lines to stress to varying degrees. Queuine supplementation to FBS depleted cells had stress and cell line specific effects. For example, Queuine protected Mock cells, QTRT2 and double KO cells from rotenone stress, but let to more cell death in QTRT1 KO cells. Queuine also sensitized cells to antimycin stress. The sensitization to arsenite and TTFA stresses was most pronounced in QTRT1 KO cells. From these results, it is clear that the influence of Queuine supplementation, FBS depletion, or tRNA-Q depletion are context specific. Given that Queuine showed an effect in the KO cell lines, it became imperative to validate whether tRNA-Q modifications are rescued upon Queuine addition in our KO cells. We conducted LC-MS validation that showed that Queuine had no impact on tRNA-Q levels in all KO cell lines, which remained undetectable [Figure 8e, Supplementary table 4]. In addition, Mock cells showed robust upregulation of tRNA-Q after Queuine addition (at 1μM concentration) in regular media and in FBS free media, with strong depletion of tRNA-Q after FBS depletion, similar to previously published works(Huber et al., 2022; Tuorto et al., 2018). We also evaluated whether supplementation with Queuine would rescue the FBS induced proliferation suppression. In all cell lines, Queuine itself, or Queuine addition to FBS depleted cells did not impact the proliferation rate [Figure 8f-i].

**Figure 8:**
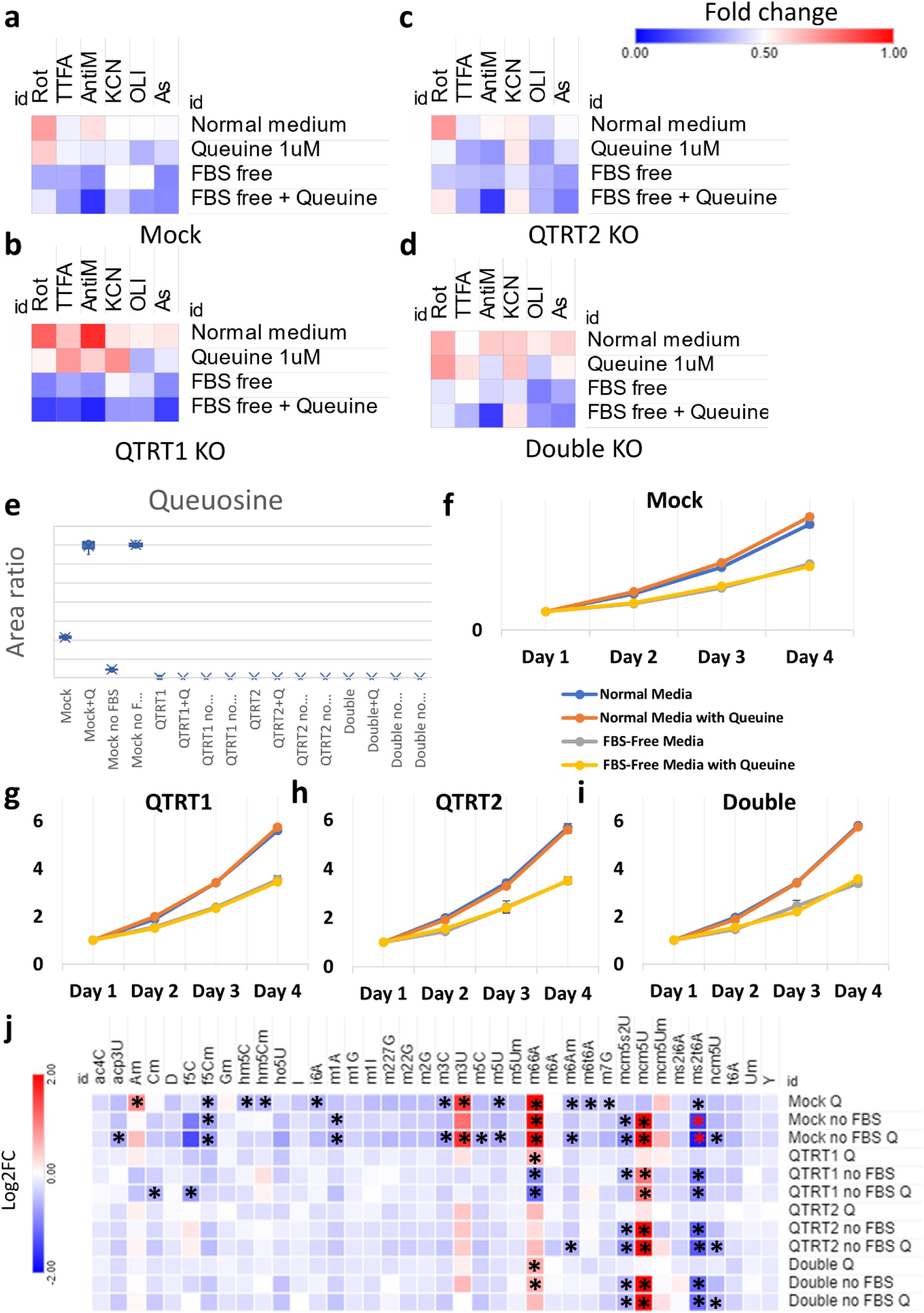
tRNA-Q loss as well as Queuine availability influence stress response and the epitranscriptome. **a-d:** Heatmaps of cell viability analysis after exposure to mitochondrial ETC inhibitors for 4 hours (Rotenone 80μM, TTFA 1.5mM, Antimycin A 50μg/ml, Potassium Cyanide (KCN) 15mM, and Oligomycin 20μM) as well as Arsenite 600μM, in normal media, in serum (FBS) free media, and with Queuine supplementation (1μM in either media). Data were normalized to controls for each cell line and expressed as fold change. **e:** Validation of tRNA-Q levels after serum deprivation and Queuine supplementation in all cell lines. **f-i:** Proliferation rates of all cell lines in different media used. **j:** Analysis of the epitranscriptional changes of serum deprivation and/or Queuine supplementation in all cell lines using LC-MS/MS. Asterisk: fold change > 1.5, *p* < 0.05. Data normalized to normal media values for each cell line and presented as Log2FC.

We further analyzed different tRNA modifications in the different media conditions to which the cells were exposed to evaluate any indirect impact of Queuine or FBS depletion on tRNA modifications that may alter the cellular response to stress [Figure 8j]. In the mock cells, both Queuine supplementation and FBS depletion had strong effects on various tRNA modifications, which were not accounted for in previous studies. Some of these effects were specific, such as upregulation of mcm5U and the downregulation of mcm5s2U and m1A after FBS depletion, or the upregulation of Am and m3U or the downregulation of m3C, i6A, hm5C, hm5Cm, and m5U with Queuine supplementation. In addition, FBS depletion and Queuine supplementation had some added effects on some modifications such as ncm5U and acp3U. Given that many of these modifications are known to be related to stress responses, it becomes clear that using either Queuine supplementation or FBS depletion as proxies to study the role of tRNA-Q on cellular stress responses are not ideal approaches. Comparing these changes to the tRNA modifications changes after TGT complex KO [Figure 5d], it becomes clear that these changes are not due to tRNA-Q upregulation secondary to Queuine supplementation, as many of these changes were in the directions observed in the KO cells. Rather, it appears that the changes observed in the KO cells might be, at least in part, attributed to unknown non-canonical activities of free Queuine that does not get incorporated into the tRNA.

We further looked into these effects in the KO cell lines. Evidently, many of the changes at the level of tRNA modifications that were attributed to Queuine supplementation were abrogated after tRNA-Q depletion. However, this might be due to the fact that these modifications were already altered after tRNA-Q and the impact of Queuine addition was minimal (note that the comparison was for each cell line’s normal media cultured cells and not between cell lines). It was consistently apparent, on the other hand, that mcm5U, ms2t6A, and mcm5s2U are all related to FBS depletion. While ncm5U showed tendency to be downregulated when Queuine was added to FBS depleted cells.

Finally, we examined how the cells epitranscriptionally respond to various stresses under normal conditions (i.e., normal media) at the level of tRNA modifications. Cells were exposed to 4 hours to various stress inducers (mitochondrial respiratory complex inhibitors + arsenite) and analyzed using LCMS [Supplementary figure 18, Supplementary table 5]. In the Mock cells, we observed significant upregulation of mcm5U only after TTFA treatment (respiratory complex II inhibitor). In QTRT1 KO cells, we observed downregulation of f5C after TTFA and potassium cyanide (KCN) treatment and upregulation of mcm5U after TTFA, KCN, and arsenite treatment. in QTRT2 KO cells, we observed upregulation of mcm5U after TTFA and arenite treatment, mcm5Um and ms2i6A after TTFA treatment, ncm5Um after arsenite treatment, and Wybutosine (yW) after TTFA treatment. In Double KO cell line, we observed mcm5U upregulation after TTFA and KCN treatment, and ms2i6A after TTFA treatment. Overall, KO cell lines’ response to stress deviated from the mock cells. There were also variations between QTRT1 & QTRT2 KO cell lines responses.

### tRNA-Q loss alters the cellular metabolome

Given the observed effects of tRNA-Q depletion on translation, mitochondrial function, and stress response, and the observed differences between QTRT1 and QTRT2 KO, we aimed next to evaluate the metabolic impact of tRNA-Q depletion, which was not previously evaluated. We employed three strategies to explore various metabolic changes in the three KO cell lines tested: targeted metabolomics/lipidomic LC-MS/MS approach [Supplementary figure 19a-b, Supplementary table 6], untargeted gas chromatography tandem mass spectrometry (GC-MS/MS) metabolomic approach [Supplementary figure 19c-d, Supplementary table 7], and a targeted LC-MS/MS analysis of glutathione and the transsulfuration pathway [Supplementary figure 19e, Supplementary table 8]. Metabolomic analysis revealed significant changes in large numbers of metabolites in tRNA-Q depleted cells. For example, Serotonin was downregulated in all three KO cell lines. The downregulation of serotonin could be linked to the tetrahydrobiopterin (BH4) pathway, which is essential for the generation of serotonin and other neurotransmitters and was shown to be linked to tRNA-Q previously(Fanet et al., 2021a; Rakovich et al., 2011). Another observation was the downregulation of 7-methylguanine, a degradation product of nucleic acids(Shapiro, 1968). 7-methylguanine, which is not related to RNA m7G modifications, is known as an inhibitor of the TGT complex and queuosine modifications(Kirsanov et al., 2022). This downregulation could be reflective of changes in global cellular RNA turnover dynamics(Kirsanov et al., 2022), or it could be a feedback response to the depletion of tRNA-Q. We also observed the downregulation of several amino acids. Most notably, we observed the downregulation of Taurine, which is itself linked to mitochondrial taurine tRNA modifications and mitochondrial function(Asano et al., 2018; Chen et al., 2016), and threonine, which is the precursor for t6A modifications(Swinehart et al., 2020; Zhou et al., 2020) which are downregulated after tRNA-Q depletion, and also linked to proper mitochondrial function. Such changes could be the cause, or an effect of the mitochondrial defects observed in the KO cells. We also observed the upregulation of glutathione and glutathione oxide in the KO cells, as well as s-adenosylhomocysteine. Such changes could impact the buffering capacity of the cells against free radicals and may in part explain the resistance to stress. However, these changes in glutathione do not explain the specificity of the stress response. Pearson’s correlation analysis using the metabolic datasets showed that QTRT1 and QTRT2 KO had moderate correlation (*R* = 0.51), while the double KO cells had lower correlation with both cell lines [Supplementary figure 19f].

Next, we utilized the entire metabolic profile of each cell line to analyze the metabolic pathways enriched using the small molecular pathway database (SMPDB) and the Kyoto encyclopedia of genes and genomes (KEGG) database using MetaboAnalyst software(Chong et al., 2018). QTRT1 KO [supplementary figure 20] led to dysregulation of phosphatidylinositol metabolism, and various mitochondrial metabolic pathways such as beta-oxidation of fatty acids and glycolysis/gluconeogenesis. QTRT2 KO [Supplementary figure 21] revealed the same tendencies as QTRT1, with some differences in the enrichment scores of various pathways, but with preservation of the overall trends. In the double KO cells [Supplementary figure 22], similar trends of dysregulation of fatty acid metabolism and mitochondrial metabolism were observed. However, the top enriched pathway was plasmalogen synthesis, which was not enriched in either QTRT1 or QTRT2 KO cells.

Overall, our metabolic profiling reveals a basis for mitochondrial dysfunction after tRNA-Q depletion as well as subtle differences between different KO cell lines that cannot be explained on the basis of tRNA-Q loss only.

## Discussion

In this work, we comprehensively analyzed the role of tRNA modifications in driving the translational shifts during mitochondrial and oxidative stress by altering codon decoding and optimality patterns. Our data revealed stress specific changes in tRNA modifications that were associated with distinct stress phenotype, such as the under-translation of mitochondrial respiratory complexes in arsenite stress due to downregulation of f5C and hm5C. Further, tRNA-Q appears to be a major player in dictating cellular responses to oxidative stress and in maintaining mitochondrial function and cellular homeostasis, in line with previous research(Huber et al., 2022; Tuorto et al., 2018). However, we observed various differences from previous studies that will be discussed herein.

In the first part of this study, we show that during mitochondrial and oxidative stresses, mRNA translation is dysregulated, and mRNA expression does not correspond to mRNA translation. This has important implications in the quest of understanding oxidative stress, as most studies have employed RNA sequencing techniques without considering the actual mRNA translation or protein profiling in studying acute stress. At the translational level, it became clear that changes in tRNA modifications levels were the main driver of mRNA translation by impacting ribosome codon decoding and optimality patterns, in line with previous studies(Chan et al., 2012; Chionh et al., 2016; Huber et al., 2022; Torrent et al., 2018). We also show that the changes in codon decoding and optimality were a driving factor behind the enrichment of many pathways and the translation of various gene sets such as respiratory chain complexes, selenoproteins, and ATF4 transcribed genes. While previous studies have shown the impact of ribosome stalling on translation of specific genes(Li et al., 2022) and how changes in codon optimality can impact specific pathways(Huber et al., 2022), these links were not studied to the extent described herein.

In order to validate the role of f5C and hm5C in the stress response, we selected ALKBH1 as a gene target(Kawarada et al., 2017). ALKBH1 itself has been a subject of controversy in the literature. Initially, ALKBH1 was reported to be a histone dioxygenase(Ougland et al., 2012). It was later reported to demethylate N1-methyladenosine (m1A) at position 58 in the tRNA(Liu et al., 2016). This demethylase activity was later shown to occur specifically during oxidative stress and to impact tRNA stability and cleavage(Rashad et al., 2020a). The impact of ALKBH1 on m1A was further validated by an independent group(Arguello et al., 2022). ALKBH1 was also shown to deoxygenate m5C into hm5C and f5C(Kawarada et al., 2017). Surprisingly, there was a report on the potential role of ALKBH1 in demethylating N6-methyladenosine in the DNA(Xie et al., 2018). However, this report was thoroughly refuted by Douvlataniotis et al(Douvlataniotis et al., 2020) where they showed that the m6A signals detected in DNA are due to potential contamination from mycobacteria in the mammalian culture. Another report from Lyu et al(Lyu et al., 2022) elegantly demonstrated that m6A gets mis-incorporated into DNA from fragmented mRNA and that ALKBH1 does not have an effect on this system. Here, our data agrees with the dioxygenase activity of ALKBH1 on m5C to generate hm5C and f5C. We did not observe an impact of ALKBH1 KO on m1A levels in the cells. It’s also notable that we could not abrogate f5C completely from the two clones generated. On the other hand, f5Cm was undetected after ALKBH1 KO. In the cytosol, hm5C and f5C are 2’O ribose methylated to f5Cm via FTSJ1 [Figure 4a]. This reaction does not occur in the mitochondria, thus f5C is the terminal product of the dioxygenation reaction catalyzed by ALKBH1(Kawarada et al., 2017). The failure to abolish f5C, while simultaneously removing f5Cm, hm5C, and hm5Cm after ALKBH1 might indicate the presence of another mitochondrial dioxygenase that could play a role in generating f5C(Arguello et al., 2022). Another possibility could be that f5C is essential for cell viability in the HEK293T cell line, and the clones that had complete depletion of f5C simply did not grow. Indeed, the rate of growth of the two colonies was tied to their respective f5C levels. We also observed that ALKBH1 KO impacted mitochondrial function as reported previously(Kawarada et al., 2017) and as predicted by our codon analysis of the respiratory complexes. ALKBH1 KO led to reduction in protein translation rates, contrary to previous reports(Liu et al., 2016) including data from our group using a different cell line (9L gliosarcoma cell line) with ALKBH1 overexpression(Rashad et al., 2022). Such discrepancies point out to a potential cell context specific role of ALKBH1 that needs to be further characterized(Rashad et al., 2020a). Interestingly, ALKBH1 KO led to the upregulation of mcm5U and tRNA-Q, mimicking what was observed in the stressed wild-type cells exposed to arsenite. Further, loss of ALKBH1, and in turn cytosolic hm5C and f5C, led to translational stress, in the form of repressed protein translation and activation of the mTOR pathway, evident by the extensive phosphorylation of P70S6K. Collectively, the data presented herein provide evidence that ALKBH1 is essential for maintaining cellular mRNA translation and mitochondrial function via its dioxygenase activity on tRNA.

Queuosine tRNA modifications (tRNA-Q) have been a subject of increasing interest in recent years. tRNA-Q was shown to be essential for protein translation(Dixit et al., 2021; Tuorto et al., 2018) and stress response to arsenite induced oxidative stress(Huber et al., 2022). QTRT1 KO mice were shown to have cognitive and memory aberrations in a sex dependent manner(Cirzi et al., 2023). We recently showed that tRNA-Q is most enriched in the brain and its levels dictate codon optimality patterns across different tissues(Ando et al., 2023). The synthesis pathway of tRNA-Q is different in mammalian cells from prokaryotes. In mammals, Queuine, the precursor for tRNA-Q, is acquired from the gut microbiome and diet(Dixit et al., 2021; Rakovich et al., 2011; Reyniers et al., 1981; Tuorto et al., 2018). The TGT complex, composed of QTRT1 and QTRT2(Behrens et al., 2018), modifies guanosine in GUN anticodons in cytosolic and mitochondrial tRNAs into queuosine(Suzuki, 2021; Zhao et al., 2023). tRNA-Q is associated with four specific amino acids; Asp (asparagine), Asn (aspartate), His (histidine), and Tyr (tyrosine). tRNA-Q is further glycosylated in mannosyl and galactosyl queuosine (manQ and galQ respectively) in cytosolic tRNAs only through two recently identified enzymes in the seminal work by Zhao et al(Zhao et al., 2023). tRNA-Q expands the codon recognition of GUN codons from NAC to NAC and NAU(Suzuki, 2021). As demonstrated here, the loss of tRNA-Q loss significantly impacted the decoding of NAU codons while NAC codons were not significantly affected.

In this work, we show that tRNA-Q dynamically change to regulate codon decoding during mitochondrial and oxidative stress, in line with previous results(Huber et al., 2022). We also show that the upregulation of tRNA-Q was essential in dictating mRNA translational shifts towards stress response genes and pathways. However, while previous works have opted for Queuine supplementation as a proxy to study tRNA-Q either in cells grown in distilled FBS containing media(Huber et al., 2022) or serum depleted media(Tuorto et al., 2018), we opted for a genetic approach. Given that our main focus was mitochondrial stress, altering serum composition could inadvertently induce confounding factors by altering metabolites that may impact mitochondrial function or stress response. Indeed, in our analysis we observed differences in how the KO cells respond to stress in normal media and also when FBS is depleted and Queuine is supplemented. Interestingly, supplementing Queuine had a deleterious effect on the cells either in normal media or in serum free media, contrary to previous report(Huber et al., 2022). However, tRNA-Q depletion had clear impact on mRNA translation, induced translational stress, and impacted mitochondrial function in line with previous reports(Hayes et al., 2020; Tuorto et al., 2018). Despite these obvious effects of tRNA-Q on cellular bioenergetics, we did not observe an impact on cellular proliferation per se. Again, in line with the recently published report using QTRT1 KO mice that had no apparent phenotype except for cognitive dysfunction(Cirzi et al., 2023).

While previous studies have opted to validate tRNA-Q depletion only via low throughput methods such as APM-gel electrophoresis or LC-MS/MS targeted for tRNA-Q only(Cirzi et al., 2023; Dixit et al., 2021; Huber et al., 2022; Tuorto et al., 2018), we analyzed all tRNA modifications in our quest to validate tRNA-Q loss after TGT KO. This approach allowed us to study the domino effect of tRNA-Q loss on the tRNA epitranscriptome. In our LC-MS/MS analysis, we observed alterations in the levels of various tRNA modifications. Interestingly, these changes were different between QTRT1 and QTRT2 KO cell lines, alluding to potential interactions at the protein levels that impact tRNA modifications. The changes in tRNA modifications levels could also partially explain the mitochondrial and translational dysfunction observed in the tRNA-Q depleted cells, such as mitochondrial dysfunction secondary to f5C, ms2i6A, t6A, or ms2t6A downregulation(Kawarada et al., 2017; Miwa et al., 2023; Zhou et al., 2020). Interestingly, supplying Queuine to Mock cells replicated some of these changes, raising the question as to whether such domino effect is the result of tRNA-Q depletion or the presence of free Queuine in the cell that cannot be incorporated into tRNA due to TGT complex KO. Another important observation that should be considered is the effect of FBS depletion and/or Queuine supplementation on tRNA modifications other than tRNA-Q. This is essential to be considered when such approaches are used to study protein translation or oxidative stress. For example, mcm5U, which is essential in stress response and for the translation of selenoproteins such as GPX4(Deng et al., 2015; Endres et al., 2015) is upregulated after FBS depletion. In addition, essential modifications such as m7G, m3C, m5C, m5U, and others were downregulated after Queuine supplementation. While we anticipate such effects to vary from cell line to cell line, taking them into account, and proper investigation of such confounding effects, should be a standard when FBS depletion or other metabolic approaches are used to study mRNA translation and the tRNA epitranscriptome.

While the translational effects of tRNA-Q and its glycosylated derivatives on codon decoding are well documented and replicated herein(Ando et al., 2023; Tuorto et al., 2018; Zhao et al., 2023), the metabolic impact of tRNA-Q loss was not studied previously. Here, we utilized multiple approaches to probe the metabolic impact of tRNA-Q loss. Our findings indicate a wide range of metabolic aberrations that impact the mitochondria as well as other cellular processes. For example, we observed downregulation of Taurine after tRNA-Q loss. Taurine itself is associated with taurine tRNA modifications, and it was shown to be important for mitochondrial function and mammalian healthy aging(Asano et al., 2018; Singh et al., 2023). L-threonine is associated with t6A modifications that are also essential for mitochondrial translation(Suzuki, 2021). Dysregulation of metabolic pathways such as beta-oxidation of fatty acids, phosphatidylinositol signaling, and glycolysis/gluconeogenesis indicate dysregulation of cellular bioenergetics and could be a consequence of translational mitochondrial dysfunction. Loss of tRNA-Q also led to the downregulation of serotonin, possibly through the tetrahydrobiopterin (BH4) pathway(Choi et al., 2006; Fanet et al., 2021a; Rakovich et al., 2011). Dysregulation of the BH4 pathway can impact mitochondrial function(Choi et al., 2006) and was linked to neurodegenerative diseases(Aziz et al., 1983; Choi et al., 2006; Fanet et al., 2021b; Foxton et al., 2007). Such metabolic connection between BH4 pathway and tRNA-Q could be a potential explanation of the phenotype observed in the work by Cirzi et al(Cirzi et al., 2023) and can be an interesting link between tRNA-Q and neurodegenerative diseases.

An important question that was not addressed in this work is how did tRNA modifications change? Is it a function of the changes in tRNA expression levels that occur after stress(Torrent et al., 2018)? Or is it due to changes in the stoichiometric levels of tRNA modifications(Pichot et al., 2023)? These questions become apparent when we consider the absence of gross changes in the transcription, translation, and protein levels of tRNA modifying enzymes responsible for the changes observed in our work after stress exposure. In addition, are the changes in tRNA modifications part of the ISR relayed from the mitochondria after ETC inhibition via the OMAI1-DELE1-HRI pathway(Fessler et al., 2020; Guo et al., 2020)? Or is there a separate sensing mechanism that led to the changes in tRNA modifications and hence translational responses to stress? Such questions are beyond the scope of the work but answering them would greatly enhance our understanding of mitochondrial stress and dysfunction, tRNA modifications’ regulation, and their disease relevance.

In conclusion, we provide a comprehensive analysis of the translational changes occurring during mitochondrial stress. We show that tRNA modifications, especially tRNA-Q, are major determinants of stress induced translational changes. We also provide comprehensive multi-omics analysis of tRNA-Q and its regulatory functions in the cell that reveals the importance of this tRNA modification in multiple processes and pathways and its potential diseases relevance.

## Methods

### Cell culture, proliferation, and stress induction

HEK293T cells were cultured in high-glucose (4.5g/l) DMEM medium supplemented with 10% fetal bovine serum (FBS) and 1% sodium pyruvate. Stress experiments were conducted in 96 well plates for cell viability assay by MTT or in 6 well plates or 10cm dishes when samples were collected for western blotting or sequencing. Cells were seeded 24 hours before the stress experiment and cultured either in regular media, or in serum free media with or without 1μM Queuine supplementation (Toronto research, Cat# Q525000) Cells were stressed by Arsenite (sodium metaArsenite), Rotenone (Respiratory complex I inhibitor), Thenoyltrifluoroacetone (TTFA; Respiratory complex II inhibitor), Antimycin A (Respiratory complex III inhibitor), potassium cyanide (KCN; Respiratory complex IV inhibitor), or Oligomycin (Respiratory complex V inhibitor) for the indicated doses and duration. Cell proliferation was conducted by passaging the cells at low concentrations into 96 wells or 6 wells plates (10,000 cells per well) and analyzing the cells over time using MTT or by cell counting by Trypan blue respectively.

### Generation of CRISPR KO cell lines

Lentivirus CRISPR sgRNAs targeting ALKBH1 gene (F: CACCGGAAACCTAATGTATGTAACC), QTRT1 gene (F: GGAGTGGCCACTGTCCCATG), QTRT2 gene (F: GGGGCTGACTGGATCGTGCA), and a non-targeting control (F: GTTCCGCGTTACATAACTTA) were designed using the BROAD Institute CRISPick design tool. They were incorporated into lentiCRISPR-v2 (Addgene, Cat# 52961) following the GeCKO protocol with minor adjustments(Sanjana et al., 2014). Briefly, the lentiCRISPR-v2 vector underwent BsmBI-v2 (NEB, Cat# R0739L) digestion and Quick CIP (NEB, Cat# M0525S) dephosphorylation. Subsequently, synthesized phosphorylated oligos were annealed in a thermocycler (Thermo Fisher Scientific) and ligated into the digested lentiCRISPRv2 backbone using the DNA ligation kit (Mighty Mix) (Takara, Cat# 6023). The ligated DNA was then transformed into One Shot Stabl3 competent cells (Thermo Fisher Scientific, Cat# C7373-03) and selected on LB agar plates with 100 µg/mL Ampicillin sodium (Fujifilm, Cat# 012-23303) to confirm the correct incorporation of the sgRNA target sequence. Following this, HEK293T cells were transfected with the respective constructs, along with envelope plasmid (pMD2.G) (Addgene, Cat# 12259), and packaging plasmid (psPAX2) (Addgene, 12260) using Lipofectamine 3000 per the manufacturer’s protocol. The 293T media containing lentivirus was collected, filtered through a 0.45 µm Millex-HV filter (Merck, Cat# SLHV033RS), and used to infect 293T cells with 8 µg/ml polybrene (Sigma Aldrich, Cat# TR-1003-G) for 24 hours. Stably infected cells were harvested and isolated through limiting dilution with 2 µg/mL Puromycin (Sigma Aldrich, Cat# P8833) over a week. Finally, single cells were expanded, and the knockout efficiency was assessed through western blotting, T7 endonuclease I assay kit following the manual’s protocol (GeneCopoeia, Cat# IC005), and LC-MS/MS.

Primers used for T7 endonuclease I assay were:

QTRT1: F: CAAATCCGTTGCCATGGTCC R: GCTGTCCTAAGGGAAACCCC

QTRT2: F: GCCCCTAGGGACCCATGATA R: TCCTTAGGCAGCCCTACTCT

### RNA sequencing

Cells were lysed in Qiazol & RNA was extracted using Qiagen’s miRNeasy mini kit with DNase digestion step. RNA concentration and purity was determined using nanodrop 1. RNA integrity was analyzed using Agilent Bioanalyzer 2100 and Agilent’s RNA 6000 nano kit, all samples had RIN > 8. mRNA sequencing was performed using NEBNext Ultra II Directional RNA Library Prep kit as per manufacturer’s instructions after mRNA enrichment by poly-A selection. Quality control of libraries was performed using Agilent’s DNA 1000 kit. NEBNext Library Quant Kit was used for quantifying libraries. Libraries were pooled and sequenced on Hiseq-X ten by Macrogen Japan (150 bp x 2). All sequencing experiments were performed with 2 or 3 biological replicates per group.

### RNA-seq data analysis

Quality control for Raw fastq files was performed using FastQC. Reads were trimmed using Trimmomatic to remove adaptor sequences and low-quality reads. Reads were aligned to hg38 assembly with NCBI Refseq annotation (Downloaded from USCS) using Hisat2. FeatureCounts was used for read counting, differential gene expression was performed using Limma-voom package. RNA-seq analysis was done on a local instance of Galaxy(Afgan et al., 2018). Pathway analysis and visualization was conducted using eVitta easyGSEA and Omics Playground.

### Ribosome profiling

To collect ribosome protected fragments (RPFs), cells were treated with the indicated drugs or collected without treatment (in case of knock-out cell analysis) by washing the cells in ice-cold PBS containing 100μg/ml cycloheximide. Cells were then scrapped and centrifuged at 4°C and lysed in 300μl polysome buffer (20mM Tris-CL (pH 7.4), 150mM NaCl, 5mM MgCl2, 1mM DTT, 100ug/ml Cycloheximide, and 1% Triton-X 100 in RNase free water). 2.5μl RNase If (100 U/μl) were added for each 100μl sample and incubated for 45 min at room temperature on a rotator mixer. Next, Qiazol was added, and RNA extracted using miRNeasy mini kit with a DNase digestion step followed by concentrating the sample using Speedvac. Samples were reconstituted in 7μl RNase free water and loaded onto 10∼15% TBE-Urea gel followed by SYBR gold staining. Gels were visualized using Chemidoc and fragments between 27∼34 nucleotides extracted and purified using Zymo small RNA PAGE recovery kit. RPFs were end rRNA depleted using NEBNext rRNA depletion kit v2 and end-repaired using T4 PNK. Libraries were then prepared using NEBNext small RNA library prep kit and sequenced as above.

### Ribo-seq data analysis

Following quality check using FastQC, libraries were trimmed and paired end reads collapsed using Seqprep. Next, bowtie2 was used to map the libraries against a reference of rRNA and tRNA to remove contaminants followed by mapping against the hg38 reference genome using Hisat2, reads counted using FeaturCounts, and differential gene expression using Limma-voom. Translation efficiency (TE) was calculated from RNA-seq and Ribo-seq data using riborex *R* package (https://github.com/smithlabcode/riborex). Metagene plots and ribosome dwelling times were analyzed using RiboToolkit(Liu et al., 2020). Pathway analysis and visualization was conducted with RNA-seq analysis.

### Codon usage and bias analysis

Isoacceptors codon frequency and total codon usage were calculated as previously reported(Rashad et al., 2022; Tumu et al., 2012). A cut-off based on the Log2 fold change and FDR values was used to select genes for analysis as indicated. For analysis of specific pathways, the genes pertaining to such pathways or gene groups were extracted from gene ontology database or relevant databases and used for analysis. Z-scores of isoacceptors frequencies were used to analyze and visualize individual genes, while T-stats representing the isoacceptors frequencies of a pathway or group of genes compared to the genome average were used to analyze and visualize gene groups. Heatmaps and correlation analysis was prepared using Morpheus. PLS-DA was conducted using mixOmics *R* package (https://www.bioconductor.org/packages/release/bioc/html/mixOmics.html).

### LC-MS/MS analysis of tRNA modifications

Mass spectrometry (LC-MS/MS) analysis of tRNA modifications was performed on two triple quadrupole platforms: Agilent 6495 and Shimadzu 8050. The analysis on Agilent platform was conducted as reported previously(Su et al., 2014) with several changes. Namely, an ultra-short (10min total time), high throughput method was used to analyze approximately 60 transitions in a given sample. First, isolated small RNAs (containing >90% tRNA) were digested in RNA digestion buffer (MgCl2 2.5mM, Tris 5mM (pH 8), Coformycin 0.1μg/ml, deferoxamine 0.1mM, Butylated hydroxytoluene (BHT) 0.1mM, Benzonase 0.25 U/μl, Calf Intestinal Alkaline Phosphatase (CIAP) 0.1U/μl, and phosphodiesterase I (PDE I) 0.003 U/μl) for 6 hours at 37°C. Digested RNAs were then injected through a Waters BEH C18 column (50 × 2.1 mm, 1.7 µm) coupled to an Agilent 1290 HPLC system and an Agilent 6495 triple-quad mass spectrometer. The LC system was conducted at 25 °C and a flow rate of 0.35 mL/min. Buffer A was composed of 0.02% formic acid (FA) in DDW. Buffer B was composed of 70% Acetonitrile + 0.02% FA. Buffer gradient program was: 2min 0% B, 4min 5.7% B, 5.9min 72& B, 6min 100% B, 6.7min 100% B, 6.75min 0% B, 10min 0% B. The UPLC column was coupled to an Agilent 6495 triple quad mass spectrometer with an electrospray ionization source in positive mode with the following parameters: gas temperature: 200 °C; gas flow: 11 L/min; nebulizer: 20 psi, sheath gas temperature: 300°C, sheath gas flow: 12 L/min, capillary voltage: 3000V and, nozzle voltage: 0V. A standard mix containing was injected to determine the retention time (RT) of the available nucleoside standards. Identification of each modification was based on its transition (either one or more), retention time, and validation by knockout cell lines if available for modifications for which standards were not available (f5Cm for example).

Collision energy was optimized beforehand for each transition. The transition list is presented in supplementary table 9. A linear UV detector, as part of the UPLC, was used to determine the UV intensity of the canonical nucleosides (U, C, G, and A) and the sum of this signal was used for normalization of the peak area. Agilent Mass hunter was used for data analysis and extraction of peak information. Data presented in supplementary tables 1,

The analysis on the Shimadzu platform was conducted similarly with several changes. First, Pentostatin was used instead of Coformycin in the digestion mix. After digestion, the digested RNAs were injected through a Waters BEH C18 column (50 × 2.1 mm, 1.7 µm) coupled to a Shimadzu Nextra Ultra-HPLC, with in-line UV detector, and a Shimadzu 8050 triple-quad mass spectrometer. Oven temperature and buffer gradients were the same as above. The following source parameters were used: Nebulizing gas flow, 2.5 L/min, heating gas flow: 10 L/min, Interface temperature: 300°C, DL temperature: 150°C, heat block temperature: 400°C, Drying gas flow: 4 L/min, and interface voltage: 1kV. Loop time was set to 0.468sec, with a dwell time between 10 and 26 msec. The transition list is presented in supplementary table 10. The signals were normalized using the UV intensity of the canonical nucleosides as above and a standard mix was used to determine the retention time and validate the results. All transitions were optimized beforehand for their collision energies and Q1 and Q3 biases.

All data presented in the heatmaps are log2 fold change to appropriate controls. Data in the graphs are presented as normalized peak areas (Mass peak area normalized by UV signal). All experiments were conducted with 3 or more biological replicates.

LCMS data in figures 3, 5, and supplementary figure 17 were conducted on the Agilent platform. LCMS data in figures 4 and 8 were conducted on the Shimadzu platform.

### Western blotting

Western blotting was conducted as previously reported(Rashad et al., 2022; Rashad et al., 2020a). Briefly, for the analysis of puromycin incorporation assay, cells were incubated in a complete growth medium with 10 μg/ml puromycin then homogenized in T-PER™ tissue protein extraction reagent (Thermo Fisher Scientific, Cat# 78510) with Triton(R) X-100 (Nacalai Tesque, Cat# 35501-15) and cOmplete™ protease inhibitor cocktail (Roche, Cat# 4693116001). Proteins were isolated from the supernatant, quantified using the Brachidonic-acid assay kit (BCA) (Thermo Fisher Scientific, Cat# 23227), separated on Mini-PROTEAN TGX gel (Bio-Rad, Cat# 4561096), and transferred to Polyvinylidene difluoride membrane (PVDF) (Bio-Rad, Cat# 1704156). Membranes were blocked in 5% Skim milk powder (Fujifilm, Cat# 190-12865) with Phosphate buffered saline with tween® (PBS-T) (Takara, Cat# T9183). The primary antibody was incubated overnight at 4°C, followed by room temperature incubation with the secondary antibody (IgG detector solution v2). Signal detection was performed using Pierce ECL substrate reagent (Thermo Fisher Scientific, Cat# 32106) on a ChemiDoc machine (Bio-Rad). To normalize protein expression, membranes were stripped with Restore™ stripping buffer (Thermo Fisher Scientific, Cat# 21059) and reprobed with Anti-beta actin antibody.

List of antibodies used:

- Beta actin antibody (1:2000 dilution, Cell signaling, Cat# 4970S).
- EIF2a antibody (1:1000 dilution, Cell signaling, Cat# 9722).
- IgG detector solution V2 (1:2000 dilution, Takara, Cat# T7122A-1).
- Mouse IgG (1:1000 dilution, Cell signaling, Cat# 7076S).
- Puromycin antibody (1:2000 dilution, Millipore, Cat# MABE343).
- P38 antibody (1:1000 dilution, Cell signaling, Cat# 9212).
- p-P38 antibody (1:1000 dilution, Cell signaling, Cat# 4511s).
- p-EIF2a antibody (1:1000 dilution, Cell signaling, Cat# 3398).
- P70S6K antibody (1:1000 dilution, Cell signaling, Cat# 9202).
- p-P70S6K antibody (1:1000 dilution, Cell signaling, Cat# 9234).
- QTRT1 antibody (1:1000 dilution, GeneTex, Cat# GTX118778).
- QTRT2 antibody (1:1000 dilution, GeneTex, Cat# GTX123016).
- Total OXPHOS Human antibody cocktail (1:1000 dilution, abcam, Cat# ab110411).

### Immunofluorescence staining

Cultured on Nunc™ Lab-Tek™ II Chamber Slide™ (Thermo Fisher Scientific, Cat# 154534PK) coated with collagen (Koken, Cat# IPC-50), 293T cells were fixed with 4% paraformaldehyde (Fujifilm, Cat# 162-16065) in Dulbecco’s phosphate-buffered saline (D-PBS) (Nacalai Tesque, Cat# 14249-24) for 10 minutes post-exposure to stressors. D-PBS was employed for cell washing before and after each step. Subsequently, cells were permeabilized with 0.1% Triton X-100 for 20 minutes at room temperature and blocked with 2% Bovine serum albumin (BSA) (Fujifilm, Cat# 010-25783) in D-PBS for 1 hour at room temperature. G3BP1 (Invitrogen, Cat# PA5-29455) or EDC4 antibodies (Santa Cruz Biotechnology, Cat# sc-374211) were used with 1% BSA and 0.1% Triton X-100 in D-PBS for an overnight incubation at 4°C, followed by a secondary incubation with Alexa Fluor 568 antibody (Invitrogen, Cat# A11036) or Alexa Fluor 488 antibody (Invitrogen, Cat# A11029) for 2 hours at room temperature. Cell nuclei were stained with 4’,6-Diamidino-2-Phenylindole, Dihydrochloride (DAPI) (1:500 dilution, Thermo Fisher Scientific, Cat# D1306) for 2 minutes at room temperature. Finally, cells were mounted with ProLong™ Diamond Antifade Mountant (Thermo Fisher Scientific, Cat# P36961). Live cell imaging utilized MitoTracker™ Green FM (Invitrogen, Cat# M7514) and MitoTracker™ Deep Red FM (Invitrogen, Cat# M22426) in a glass bottom dish (Matsunami, Cat# D11131H). Confocal microscopy with Fluoroview v6 software (Olympus FV3000) was employed for image acquisition.

### Cell lysis for widely targeted metabolomics by GC-MS/MS and UHPLC-MS/MS

We employed widely targeted metabolomics by GC-MS/MS and UHPLC-MS/MS on cell lysate from following previously established protocols(Saigusa et al., 2021b). The frozen cell pellet (50 µl / 1 × 10^6 cells) was thawed on ice and suspended in ethanol-phosphate buffer (ethanol/0.01 M phosphate buffer (pH 7.5), 85/15, v/v%). The sample was sonicated for 3 min in ice bath and snap frozen in liquid nitrogen for 30 sec, and the process was twice repeated. Afterwards the lysates were centrifuged for 5 min at 2 °C at 30,030 × *g*, and the supernatant was transferred to a 1.5-ml sample tube and stored at −80 °C.

### Widely targeted metabolomics by GC-MS/MS

GC-MS/MS analysis was performed using a previously described method(Yamauchi et al., 2021). Briefly, cell lysate (50 μl) was mixed with the extraction solution (260 μl, water/methanol/chloroform, 1/2.5/1 (v/v/v)) containing 10 µl of 2-isopropylmalic acid (0.5 mg/ml, Sigma-Aldrich, Tokyo, Japan) as an internal standard (IS), and the sample was incubated for 30 min at 37°C with shaking at 1,200 rpm by BioShaker (M-BP-220U, TYTECH, Kyoto, Japan). After centrifuged at 16,000 × *g* for 3 min at 4°C, the upper phase (225 µl) was transferred to another 1.5-ml tube, and Mili-Q water (200 µl) was added to the tube and mixed for 30 sec. The sample was centrifuged at 16,000 × *g* for 3 min at 4°C, and the supernatant (280 µl) was transferred to another 1,5-ml tube, and then dried in a vacuum centrifuge system before being lyophilized using a freeze dryer. For the first step of the derivatization (oximation), the methoxyamine hydrochloride (80 μl, 20 mg/ml, Sigma-Aldrich, Tokyo, Japan) dissolved in pyridine (TCI, Tokyo, Japan) was added to the lyophilized sample. Then the sample was sonicated for 10 min with the ultrasonic bath and incubated for 90 min at 30°C with shaking at 1,200 rpm by ThermoMixer C (Eppendorf,

Tokyo, Japan). Next, *N*-methyl-*N*-trimethylsilyl-trifluoroacetamide (40 µl, GL Science, Tokyo, Japan) was added to the sample for the second step of the derivatization. The sample was incubated for 30 min at 37°C 1,200 rpm with shaking at 1,200 rpm by ThermoMixer C. After centrifuged at 16,000 × *g* for 5 min at 4°C, the supernatant was transferred to the sample vial for subjecting to GC-MS/MS system. The GC-MS/MS analysis was performed on a TQ8040 (Shimadzu Co., Kyoto Japan) equipped with a fused silica capillary column (BPX-5; 30 m × 0.25 mm inner diameter, film thickness: 0.25 μm; Shimadzu Co.). The front inlet temperature was 250°C. The flow rate of helium gas through the column was 39 cm/sec. The column temperature was held at 60°C for 2 min and then raised by 15°C/min to 330°C and held there for 3 min. The interface and ion-source temperatures were 280°C and 200°C, respectively. The metabolites were detected and annotated by performing the software (Smart Metabolites Database, Shimadzu, Co.), which contained the relevant multiple reaction monitoring (MRM) condition with and optimal GC analytical conditions, MRM parameters, and retention index employed for each metabolite. All data was imported to the software (Traverse MS, Reifycs Inc., Tokyo, Japan) for peak picking. The area ratio calculated with the IS of all metabolites were exported and MetaboAnalyst 5.0 (https://www.metaboanalyst.ca/) software was used for the prior multivariate analysis.

### Sample preparation for UHPLC-MS/MS for glutathione (GSH) analysis

Sample was suspended in of methanol (100 µl) containing 0.1% formic acid and the ISs (0.1 µg/mL S-adenosylmethionine-^13^C_5_^15^N and 0.1 µg/mL GSH-^13^C ^15^N), and mixed for 30 sec followed by sonication for 5 min. After centrifugation at 16,400 x *g* for 10 min at 4°C, the supernatant was transferred to a vial, and the sample (3 µl) was subjected to the ultra-high-performance liquid chromatography triple quadrupole mass spectrometry (UHPLC-MS/MS).

### UHPLC-MS/MS for GSH analysis

The UHPLC-MS/MS analysis was performed with Acquity™ UPLC I-class system equipped with Xevo TQ-XS MS/MS system (Waters Corp. Milford, UK). The transitions for multiple reaction monitoring were previously reported(Nishizawa et al., 2020). The capillary voltages of electrospray ionization for positive and negative ion mode were 4.0 kV and 2.5 kV, respectively. The cone voltage, cone gas (nitrogen) flow rate, desolvation temperature, desolvation gas flow, collision gas flow and nebulization gas flow were 64 V, 150 L/hr, 600°C, 1000 L/hr, 0.15 ml/min 7.00 bar, respectively. The UHPLC condition was modified with the previous publication(Saigusa et al., 2016). LC separation was performed with an Acquity UPLC BEH Amide column (1.7 μm, 2.1 × 150 mm, Waters Corp.) kept at 20 °C with a gradient elution using solvent A (10 mmol/l NH_4_HCO_3_, adjusted to pH 9.2 using ammonia solution) and B (acetonitrile) at 0.2 ml/min. All data was analyzed by to the software (Traverse MS).

### Widely targeted metabolomics by UHPLC-MS/MS

The sample preparation and UHPLC-MS/MS analysis were followed by the MxP® Quant 500 kit (Biocrates Life Sciences AG, Innsbruck, Austria). The methods were followed by the kit protocols previously described(Saigusa et al., 2021a). In brief, the blank, calibration standard, quality control, and cell lysate samples (10 μl, each) were added to predefined 24-well of the 96-well plate. The plate was dried under the vacuumed pressure manifold (Positive Pressure-96 Processor, Waters), and 5% phenyl isothiocyanate was added for derivatization. The plate was dried again, and the analytes were eluted by the solution of methanol containing 5 mmol/l ammonium acetate. The plate was duplicated into two plates for LC and flow injection analysis mode. The UHPLC-MS/MS analysis was performed by the same system described above. The optimal UHPLC-MS/MS conditions were automatically set using the method in the kit. The data were collected by MassLynx 4.2 software (Waters). The concentration of metabolites was calculated using the manufacturer’s protocol using the software (MetIDQ Oxygen).

### Seahorse analysis

We employed Seahorse XF96 analyzer (Agilent, CA, USA) and Mito-stress test kit (Agilent, Cat# 103015-100) along with Seahorse XFe96/XF Pro FluxPak (Agilent, Cat# 103792-100) and Seahorse XF DMEM medium (without Phenol Red/pH 7.4/with HEPES/500 mL, Agilent, Cat# 103575-100) to quantify Oxygen Consumption Rate (OCR) and Extracellular Acidification Rate (ECAR). Initially, 35,000 cells/well were seeded in 96-well plates on day one, while the XFe96 sensor cartridge hydrated in 200 µL calibration medium. On the next day, the cell medium was replaced with DMEM, and cells were incubated at 37℃ in a CO₂-free incubator for 1 hour. During this time, 1 µM Oligomycin, 0.5 µM FCCP, Antimycin A, and Rotenone were loaded into the drug ports of the cartridge. After loading, the sensor plate was calibrated in the analyzer. Following calibration, the cell culture plate was loaded, and the analysis initiated with the program: Set: Mixture 2 minutes, Measure 3 minutes, 4 sets in all steps. Ports: (A: Oligomycin / B: FCCP / C: Antimycin/Rotenone).

## Statistical analysis

Statistical analysis was performed using SPSS v20 (IBM Corp.). Student’s t-test or ANOVA with Turkey’s post-hoc were conducted based on the number of groups in the comparison.

Levene’s test was used to analyze the homogeneity of variance of samples. *P* value less than 0.05 and fold change of more than 1.5 were used as cutoffs for statistical significance on multiple comparisons.

## Data availability

The raw sequencing data were deposited in the sequencing read archive (PRJNA1061814, PRJNA1063528, and PRJNA1076587). Mass spectrometry data (RNA modifications and metabolomics) are provided in the supplementary data as normalized (corrected) peak areas.

## Author contributions

**S.R.:** Conception, Study design, Protocol development, LC-MS/MS analysis of RNA modifications, Formal analysis, Bioinformatics analysis, Administration, Writing. **S.R., S.A, A.M.:** Conducted experiments. **S.R., S.A., Y.Z.:** Sequencing library preparation. **S.R., G.S., J.S.:** LC-MS/MS methods development for RNA modifications analysis. **T.F. Y.I. D.S.:** Metabolomics analysis. **S.R., J.X., S.B**.: Codon usage bias analysis. **T.J.B.:** Codon counting algorithm development. **All authors:** Critically revised the manuscript. **S.R., D.S., K.N.:** Funding.

## Funding sources

This work was supported by the Japan society for promotion of science (JSPS) KAKENHI grants number 20K16323, 20KK0338, and 23H02741 for **S.R**., by JST Moonshot R&D project number JPMJPS2023 for **K.N**., and partially by JSPS KAKENHI grants Number JP20H03374, JP23H02625 and Japan Agency for Medical Research and Development (AMED) grant Number 23ek0210168h0002 for **D.S**. **S.R.B** was supported by NIEHS Training Grant in Environmental Toxicology (T32-ES007020). **J.X.** was supported by MIT UROP program.

## Supporting information

Suppmenetary tables

## Acknowledgment

The authors report no conflict of interest nor any ethical adherences regarding this work.

## Notes

### Competing Interest Statement

The authors have declared no competing interest.

