## Supplementary figures for "Translational response to mitochondrial stresses is orchestrated by tRNA modifications"

### Supplementary tables information

**Supplementary table 1:** Normalized peak areas of RNA modifications after stress exposure.

**Supplementary table 2:** Pathways used for codon analysis. Related to supplementary figure 7e.

**Supplementary table 3:** Normalized peak area of RNA modifications after ALKBH1 KO.

**Supplementary table 4:** Normalized peak areas of RNA modifications after serum deprivation or Queuine supplementation.

**Supplementary table 5:** Normalized peak area of RNA modifications after exposing QTRT1 (Q1), QTRT2 (Q2), or Double (Q3) KO cells to stress. AM: antimycin A. C: control. KCN: potassium cyanide. OLI: Oligomycin. Ro: Rotenone. TF: TTFA. As: Arsenite.

**Supplementary table 6:** Global targeted LC-MS/MS metabolomics analysis in tRNA-Q KO cells. Data presented as normalized peak areas.

**Supplementary table 7:** Global untargeted GC-MS/MS metabolomics analysis in tRNA-Q KO cells. Data presented as normalized peak areas.

**Supplementary table 8:** Targeted glutathione and transsulfuration pathway analysis via LC-MS/MS in tRNA-Q KO cells. Data presented as normalized peak areas.

**Supplementary table 9:** The transition list used in Agilent 6495 LC-MS/MS analysis of RNA modifications.

**Supplementary table 10:** The transition list used in Shimadzu 8050 LC-MS/MS analysis of RNA modifications.

### Supplementary figures

**Supplementary figure 1: Mitochondrial stress induces transcriptional and translational dysregulation (Supplementary to figure 2): a:** Gene UMAP from RNA-seq data. **b:** GOBP activation matrix from RNA-seq data. **c-e:** Volcano plot for differentially expressed genes in the Ribo-seq analysis. **f:** Cluster heatmap for the Ribo-seq data. **g:** GOBP activation matrix from the Ribo-seq data.

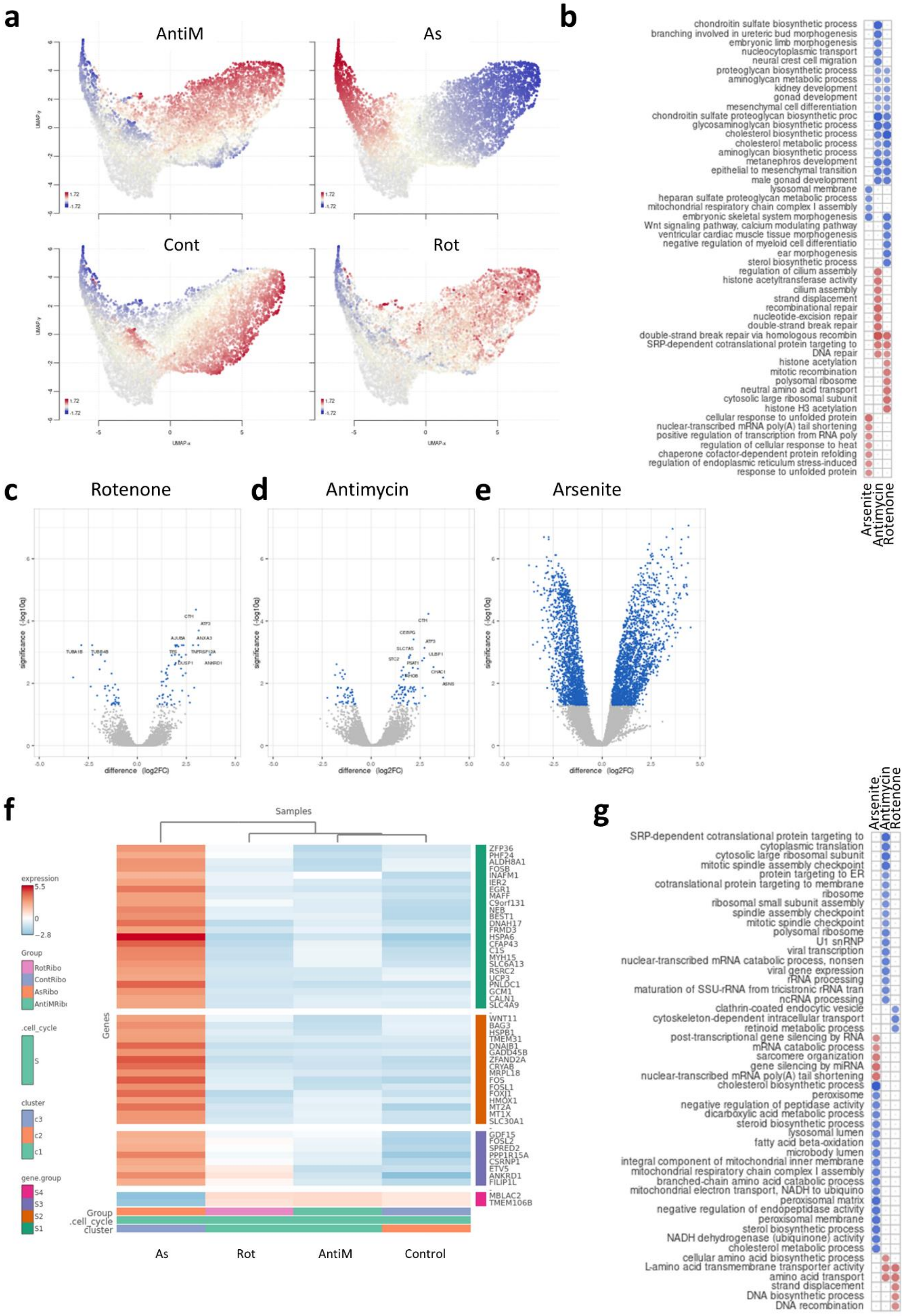
a
AntiM
As
Cont
Rot
UMAP_x
UMAP_y
b
chondroitin sulfate biosynthetic process
branching involved in ureteric bud morphogenesis
embryonic limb morphogenesis
nucleocytoplasmic transport
neural crest cell migration
proteoglycan biosynthetic process
aminoglycan metabolic process
kidney development
gonad development
mesenchymal cell differentiation
chondroitin sulfate proteoglycan biosynthetic proc
glycosaminoglycan biosynthetic process
cholesterol biosynthetic process
cholesterol metabolic process
aminoglycan biosynthetic process
metanephros development
epithelial to mesenchymal transition
male gonad development
lysosomal membrane
heparan sulfate proteoglycan metabolic process
mitochondrial respiratory chain complex I assembly
embryonic skeletal system morphogenesis
Wnt signaling pathway, calcium modulating pathway
ventricular cardiac muscle tissue morphogenesis
negative regulation of myeloid cell differentiatio
ear morphogenesis
sterol biosynthetic process
regulation of cilium assembly
histone acetyltransferase activity
cilium assembly
strand displacement
recombinational repair
nucleotide-excision repair
double-strand break repair
double-strand break repair via homologous recombin
SRP-dependent cotranslational protein targeting to
DNA repair
histone acetylation
mitotic recombination
polysomal ribosome
neutral amino acid transport
cytosolic large ribosomal subunit
histone H3 acetylation
cellular response to unfolded protein
nuclear-transcribed mRNA poly(A) tail shortening
positive regulation of transcription from RNA poly
regulation of cellular response to heat
chaperone cofactor-dependent protein refolding
regulation of endoplasmic reticulum stress-induced
response to unfolded protein
Arsenite
Antimycin
Rotenone
c
Rotenone
d
Antimycin
e
Arsenite
significance (-log10q)
difference (log2FC)
f
Samples
expression
5.5
0
−2.8
Group
RotRibo
ContRibo
AsRibo
AntiMRibo
.cell_cycle
S
cluster
c3
c2
c1
gene.group
S4
S3
S2
S1
Genes
ZFP36
PHF24
ALDH8A1
FOSB
INAFM1
IER2
EGR1
MAFF
C9orf131
NEB
BEST1
DNAH17
FRMD3
HSPA6
CFAP43
C1S
MYH15
SLC6A13
RSRC2
UCP3
PNLDC1
GCM1
CALN1
SLC4A9
WNT11
BAG3
HSPB1
TMEM31
DNAJB1
GADD45B
ZFAND2A
CRYAB
MRPL18
FOS
FOSL1
FOXJ1
HMOX1
MT2A
MT1X
SLC30A1
GDF15
FOSL2
SPRED2
PPP1R15A
CSRNP1
ETV5
ANKRD1
FILIP1L
MBLAC2
TMEM106B
Group
.cell_cycle
cluster
As
Rot
AntiM
Control
g
Arsenite
Antimycin
Rotenone
SRP-dependent cotranslational protein targeting to
cytoplasmic translation
cytosolic large ribosomal subunit
mitotic spindle assembly checkpoint
protein targeting to ER
cotranslational protein targeting to membrane
ribosome
ribosomal small subunit assembly
spindle assembly checkpoint
mitotic spindle checkpoint
polysomal ribosome
U1 snRNP
viral transcription
nuclear-transcribed mRNA catabolic process, nonsen
viral gene expression
rRNA processing
maturation of SSU-rRNA from tricistronic rRNA tran
ncRNA processing
clathrin-coated endocytic vesicle
cytoskeleton-dependent intracellular transport
retinoid metabolic process
post-transcriptional gene silencing by RNA
mRNA catabolic process
sarcomere organization
gene silencing by miRNA
nuclear-transcribed mRNA poly(A) tail shortening
cholesterol biosynthetic process
peroxisome
negative regulation of peptidase activity
dicarboxylic acid metabolic process
steroid biosynthetic process
lysosomal lumen
fatty acid beta-oxidation
microbody lumen
integral component of mitochondrial inner membrane
mitochondrial respiratory chain complex I assembly
branched-chain amino acid catabolic process
mitochondrial electron transport, NADH to ubiquino
peroxisomal matrix
negative regulation of endopeptidase activity
peroxisomal membrane
sterol biosynthetic process
NADH dehydrogenase (ubiquinone) activity
cholesterol metabolic process
cellular amino acid biosynthetic process
L-amino acid transmembrane transporter activity
amino acid transport
strand displacement
DNA biosynthetic process
DNA recombination

**Supplementary figure 2: Occupancy metagene plots after stress exposure. a-c:** Across CDS. **d-f:** Downstream from the start codon. **g-i:** Upstream from the stop codon.

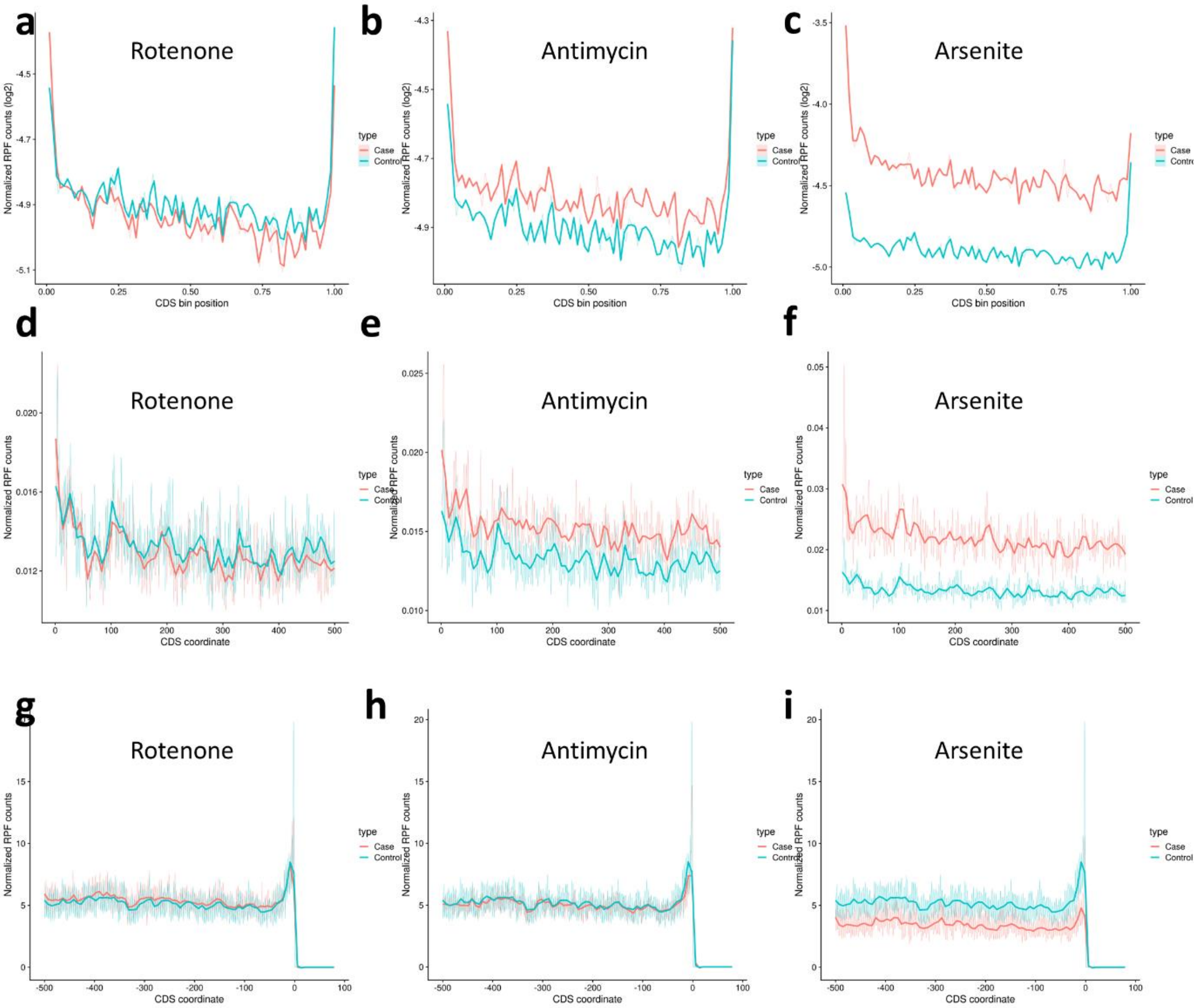

a
Rotenone
b
Antimycin
c
Arsenite
d
Rotenone
e
Antimycin
f
Arsenite
g
Rotenone
h
Antimycin
i
Arsenite
Normalized RPF counts (log2)
CDS bin position
Normalized RPF counts
CDS coordinate
type
Case
Control

**Supplementary figure 3: a:** Chemical structure for f5C and hm5C. **b:** expression of f5C in different stresses. Data presented as normalized peak areas. **c:** Expression of hm5C in different stresses. Asterisk: fold change > 1.5 and $p < 0.05$. **d:** Expression of various genes related to different respiratory chain complexes in the Ribo-seq dataset in different stresses.

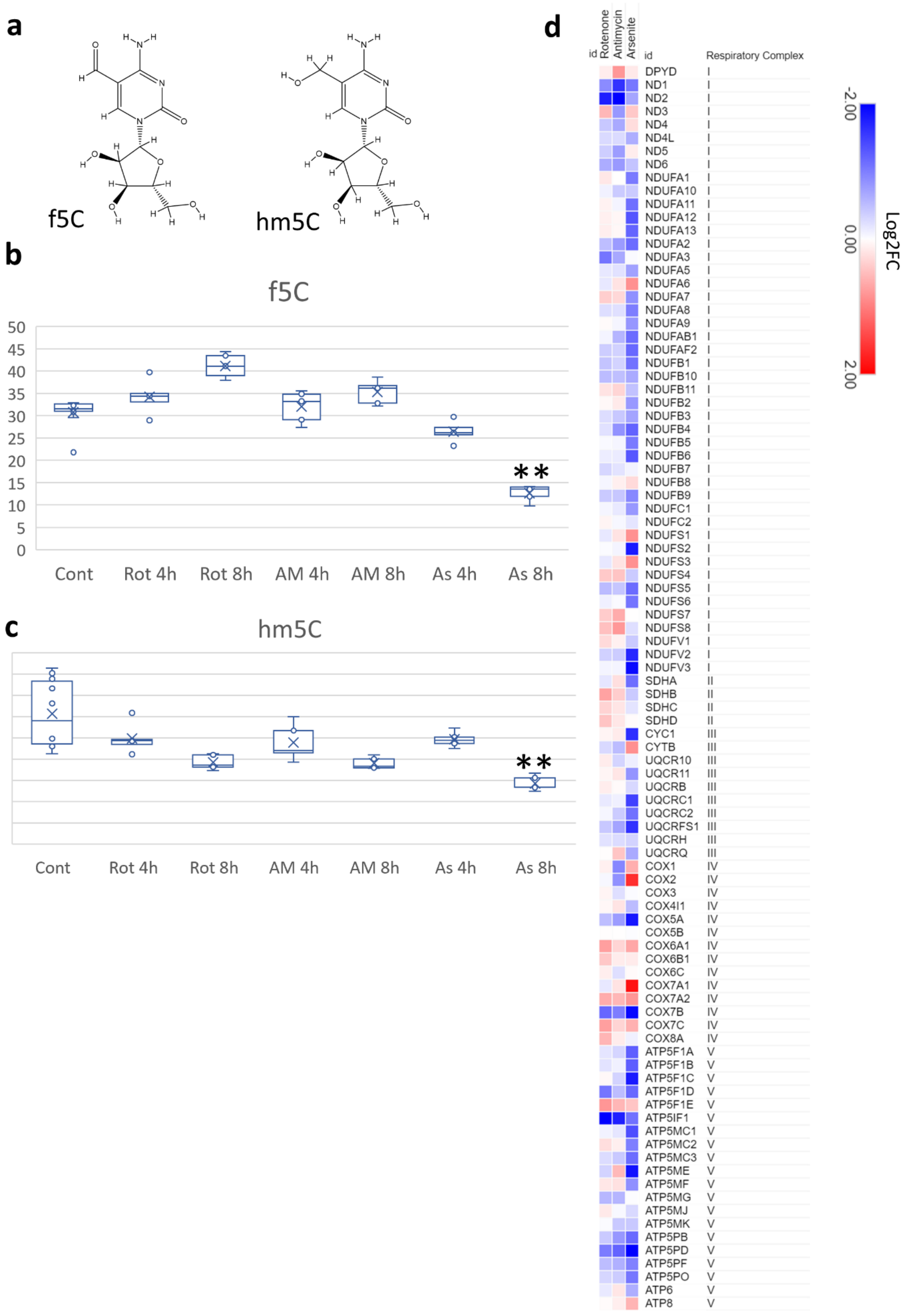

a
f5C
hm5C
b
f5C
50
45
40
35
30
25
20
15
10
5
0
**
Cont
Rot 4h
Rot 8h
AM 4h
AM 8h
As 4h
As 8h
c
hm5C
**
Cont
Rot 4h
Rot 8h
AM 4h
AM 8h
As 4h
As 8h
d
id
Rotenone
Antimycin
Arsenite
id
Respiratory Complex
DPYD I
ND1 I
ND2 I
ND3 I
ND4 I
ND4L I
ND5 I
ND6 I
NDUFA1 I
NDUFA10 I
NDUFA11 I
NDUFA12 I
NDUFA13 I
NDUFA2 I
NDUFA3 I
NDUFA5 I
NDUFA6 I
NDUFA7 I
NDUFA8 I
NDUFA9 I
NDUFAB1 I
NDUFAF2 I
NDUFB1 I
NDUFB10 I
NDUFB11 I
NDUFB2 I
NDUFB3 I
NDUFB4 I
NDUFB5 I
NDUFB6 I
NDUFB7 I
NDUFB8 I
NDUFB9 I
NDUFC1 I
NDUFC2 I
NDUFS1 I
NDUFS2 I
NDUFS3 I
NDUFS4 I
NDUFS5 I
NDUFS6 I
NDUFS7 I
NDUFS8 I
NDUFV1 I
NDUFV2 I
NDUFV3 I
SDHA II
SDHB II
SDHC II
SDHD II
CYC1 III
CYTB III
UQCR10 III
UQCR11 III
UQCRB III
UQCRC1 III
UQCRC2 III
UQCRFS1 III
UQCRH III
UQCRQ III
COX1 IV
COX2 IV
COX3 IV
COX4I1 IV
COX5A IV
COX5B IV
COX6A1 IV
COX6B1 IV
COX6C IV
COX7A1 IV
COX7A2 IV
COX7B IV
COX7C IV
COX8A IV
ATP5F1A V
ATP5F1B V
ATP5F1C V
ATP5F1D V
ATP5F1E V
ATP5IF1 V
ATP5MC1 V
ATP5MC2 V
ATP5MC3 V
ATP5ME V
ATP5MF V
ATP5MG V
ATP5MJ V
ATP5MK V
ATP5PB V
ATP5PD V
ATP5PF V
ATP5PO V
ATP6 V
ATP8 V
-2.00
0.00
2.00
Log2FC

**Supplementary figure 4:** Structure and expression of tRNA modifications after stress exposure. **a-b:** mcm5U, **c-d:** Queuosine (tRNA-Q), **e-f:** manQ, and **g-h:** galQ. Data presented as normalized peak areas. Asterisk: fold change > 1.5 and $p < 0.05$.

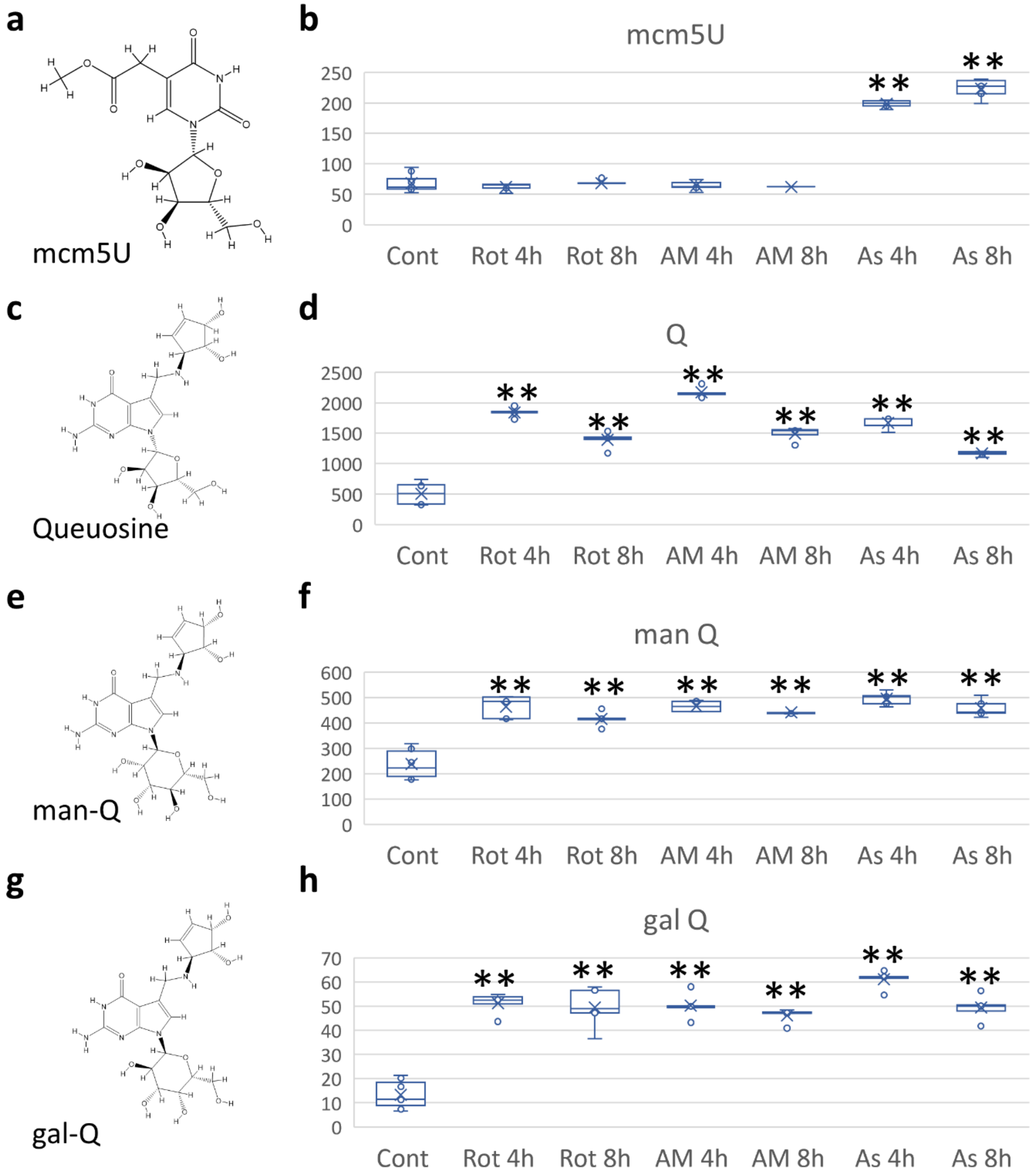
a
mcm5U
b
mcm5U
250
200
150
100
50
0
Cont
Rot 4h
Rot 8h
AM 4h
AM 8h
As 4h
As 8h
c
Queuosine
d
Q
2500
2000
1500
1000
500
0
e
man-Q
f
man Q
600
500
400
300
200
100
0
g
gal-Q
h
gal Q
70
60
50
40
30
20
10
0

**Supplementary figure 5:** A-site pausing after stress exposure. Red asterisk: mcm5U codons. Black: NAC Q-codons. Orange: NAU Q-codons. Blue: hm5C codon.

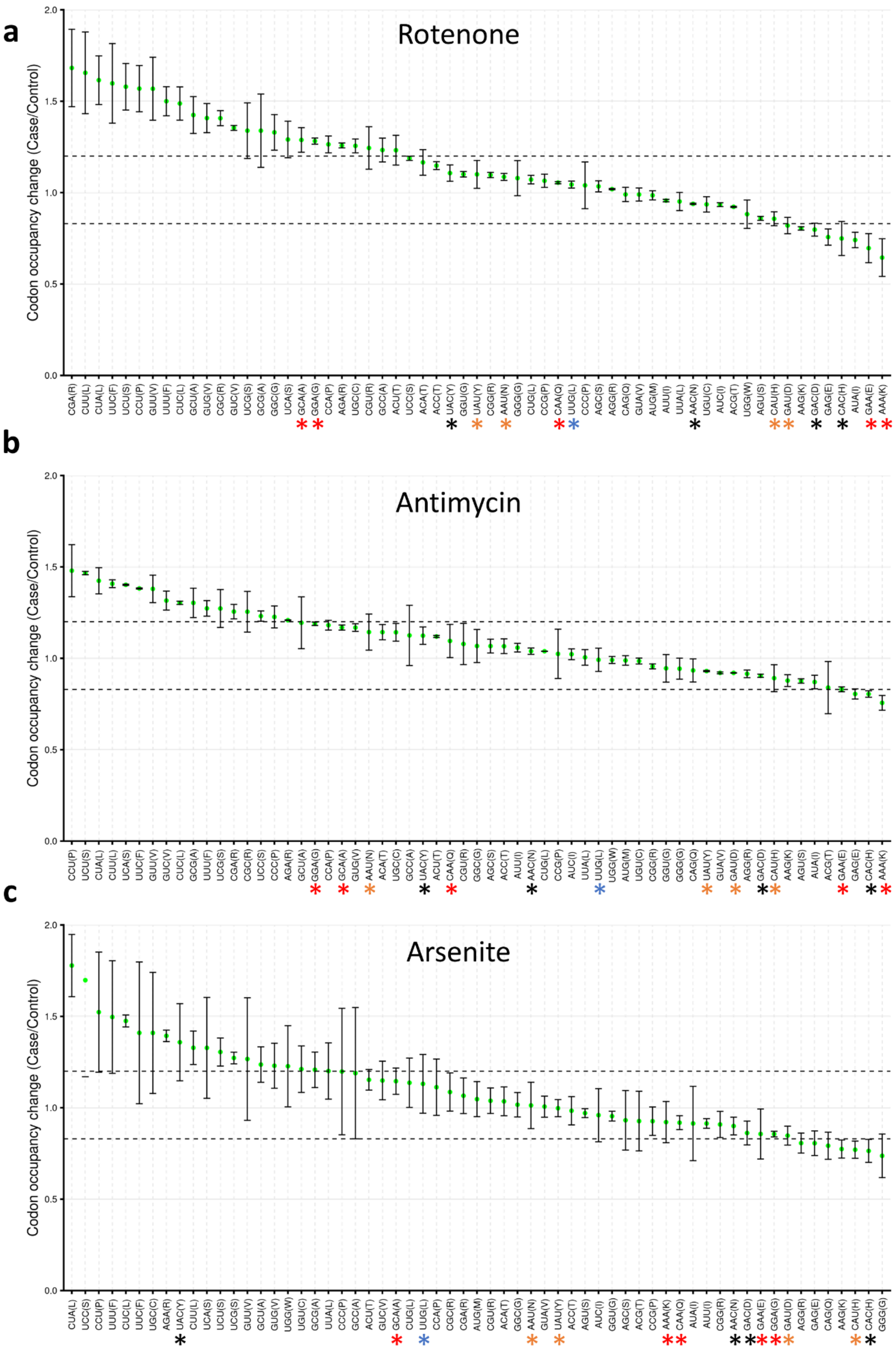
a
Rotenone
b
Antimycin
c
Arsenite
Codon occupancy change (Case/Control)
2.0
1.5
1.0
0.5
0.0

**Supplementary figure 6: Analysis of codon changes after stress. a:** Pearson's correlation analysis across different datasets using isoacceptors codon frequencies as input. **b:** Isoacceptors **c**odon frequency analysis of respiratory chain complex genes. **c:** Isoacceptors **c**odon frequency analysis of selenoproteins detected in the sequencing analysis. **d:** Expression of selenoproteins in the Ribo-seq datasets. **e:** Pearson's correlation analysis using the expression of selenoproteins as input.

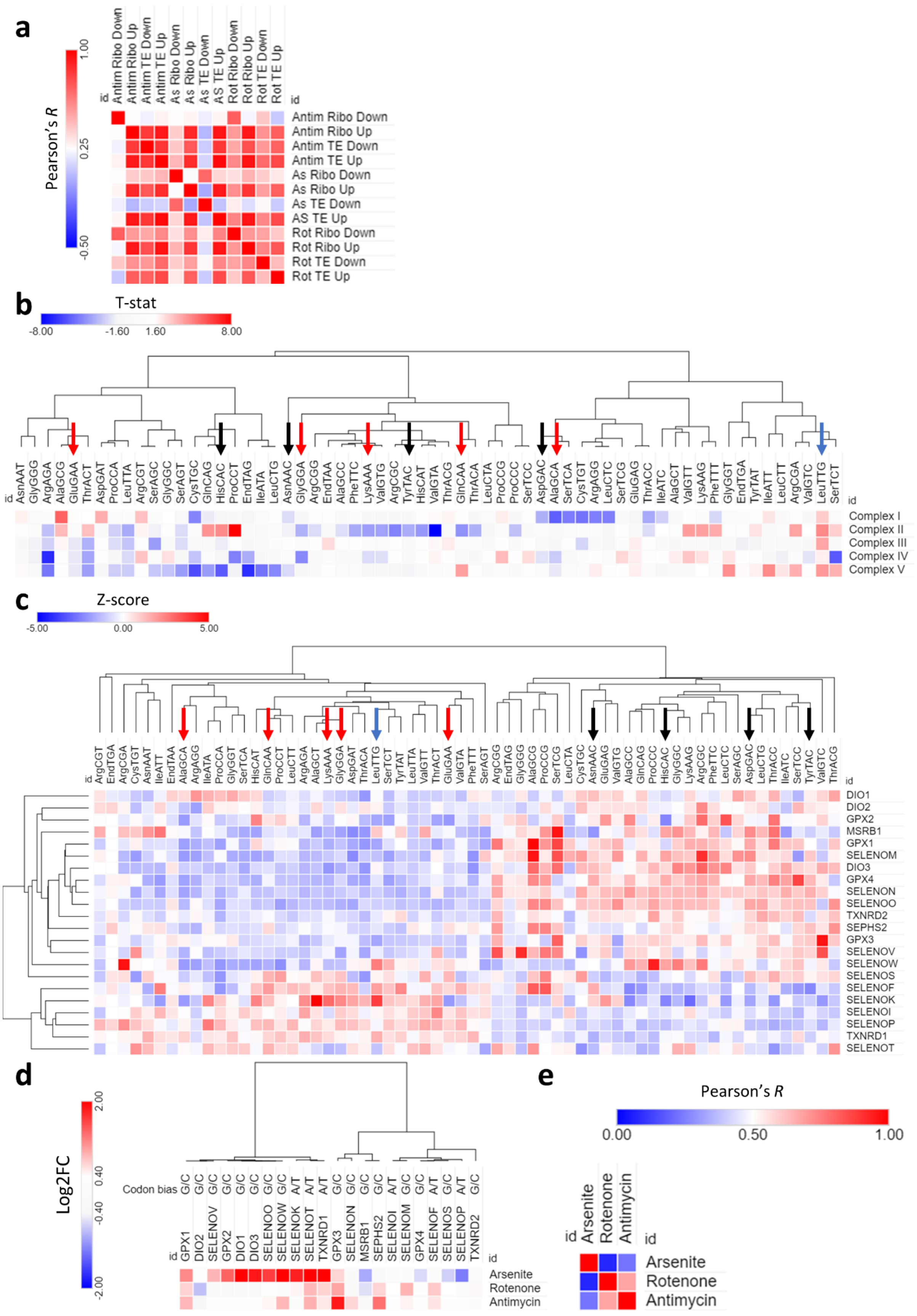
a
Pearson's R
1.00
0.25
-0.50
id
Antim Ribo Down
Antim Ribo Up
Antim TE Down
Antim TE Up
As Ribo Down
As Ribo Up
As TE Down
AS TE Up
Rot Ribo Down
Rot Ribo Up
Rot TE Down
Rot TE Up
b
T-stat
-8.00
-1.60
1.60
8.00
id
AsnAAT GlyGGG ArgAGA AlaGCG GluGAA ThrACT AspGAT ProCCA LeuTTA ArgCGT SerAGC GlyGGC SerAGT CysTGC GlnCAG HisCAC ProCCT EndTAG IleATA LeuCTG AsnAAC GlyGGA ArgCGG EndTAA AlaGCC PheTTC LysAAA ValGTG ArgCGC TyrTAC HisCAT ValGTA ThrACG GlnCAA ThrACA LeuCTA ProCCG ProCCC SerTCC AspGAC AlaGCA SerTCA CysTGT ArgAGG LeuCTC SerTCG GluGAG ThrACC IleATC AlaGCT ValGTT LysAAG PheTTT GlyGGT EndTGA TyrTAT IleATT LeuCTT ArgCGA ValGTC LeuTTG SerTCT
Complex I
Complex II
Complex III
Complex IV
Complex V
c
Z-score
-5.00
0.00
5.00
id
ArgCGT EndTGA ArgCGA CysTGT AsnAAT IleATT EndTAA AlaGCA ArgAGG IleATA ProCCA GlyGGT SerTCA HisCAT GlnCAA ProCCT LeuCTT ArgAGA AlaGCT LysAAA GlyGGA AspGAT ThrACA LeuTTG SerTCT TyrTAT LeuTTA ValGTT ThrACT GluGAA ValGTA PheTTT SerAGT ArgCGG EndTAG GlyGGG AlaGCG ProCCG SerTCG LeuCTA CysTGC AsnAAC GluGAG ValGTG AlaGCC GlnCAG ProCCC HisCAC GlyGGC LysAAG ArgCGC PheTTC LeuCTC SerAGC AspGAC LeuCTG ThrACC IleATC SerTCC TyrTAC ValGTC ThrACG
DIO1
DIO2
GPX2
MSRB1
GPX1
SELENOM
DIO3
GPX4
SELENON
SELENOO
TXNRD2
SEPHS2
GPX3
SELENOV
SELENOW
SELENOS
SELENOF
SELENOK
SELENOI
SELENOP
TXNRD1
SELENOT
d
Log2FC
2.00
0.40
-0.40
-2.00
Codon bias G/C G/C G/C G/C G/C G/C G/C G/C A/T A/T A/T G/C G/C G/C A/T G/C G/C A/T G/C G/C A/T G/C
id GPX1 DIO2 SELENOV GPX2 DIO1 DIO3 SELENOO SELENOW SELENOK SELENOT TXNRD1 GPX3 SELENON MSRB1 SEPHS2 SELENOI SELENOM GPX4 SELENOF SELENOS SELENOP TXNRD2
Arsenite
Rotenone
Antimycin
e
Pearson's R
0.00
0.50
1.00
id
Arsenite
Rotenone
Antimycin

**Supplementary figure 7: Analysis of codon changes after stress (Cont.). a:** PLS-DA on selenoproteins using their expression in the arsenite Ribo-seq dataset after dividing them into up and down regulated and using isoacceptors frequencies as input. **b:** PLS-DA of the variables (codons) impacting selenoproteins expression. **c:** VIP analysis of codons contribution to selenoproteins expression using isoacceptors frequencies as input. **d:** VIP analysis of codons contribution to selenoproteins expression using total codon frequencies as input. **e:** Isoacceptors codon frequencies analysis of enriched pathways using the Ribo-seq datasets as input. **f:** PLS-DA on the enriched pathways using the isoacceptors codon frequencies as input. **g:** PLS-DA of the variables (codons) impacting pathway enrichment. **h:** VIP analysis of codons contribution to pathways enrichment using isoacceptors frequencies as input.

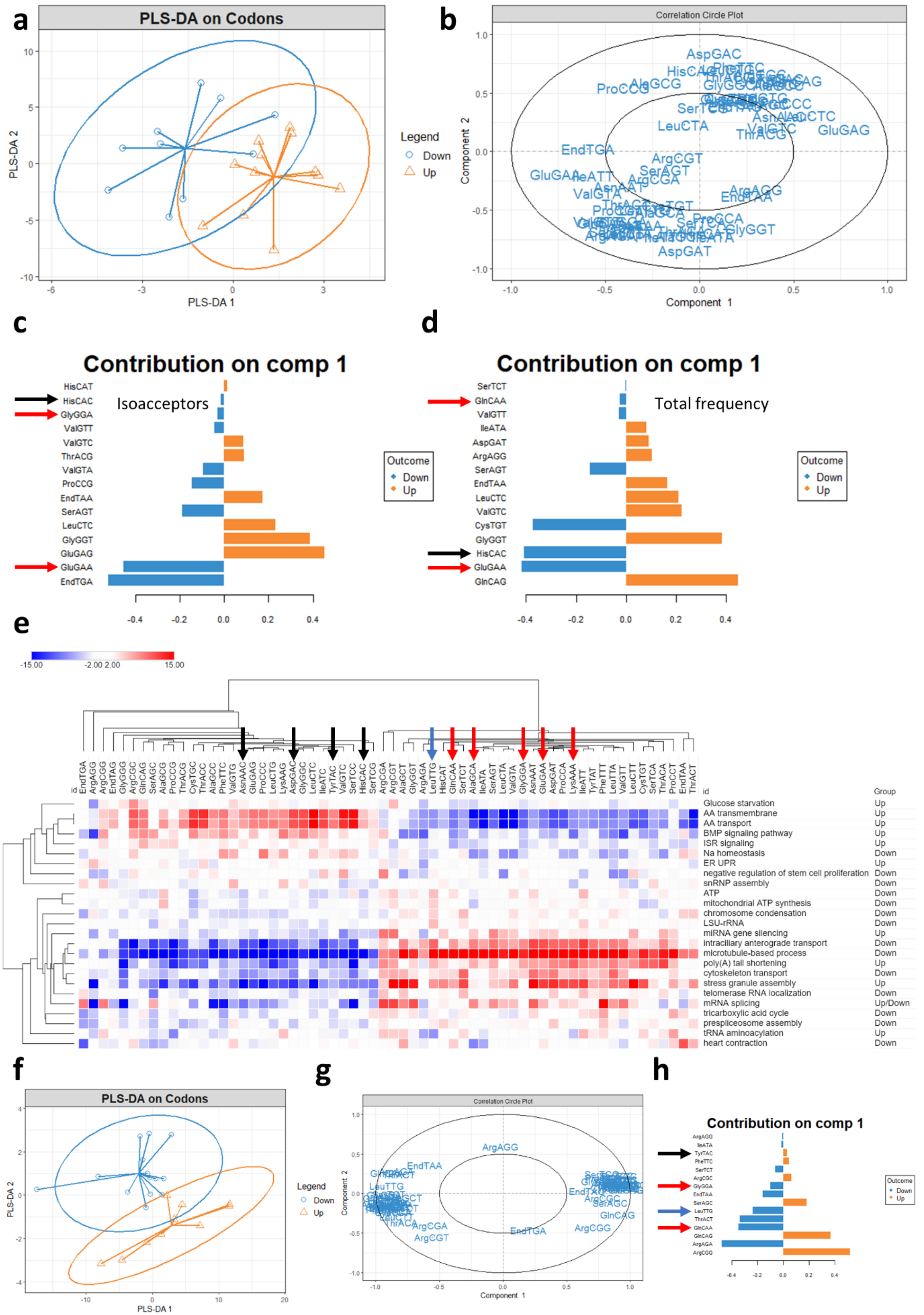

a
PLS-DA on Codons
PLS-DA 1
PLS-DA 2
Legend
Down
Up
b
Correlation Circle Plot
Component 1
Component 2
AspGAC
ProCCG
LeuCTA
GluGAG
EndTGA
ArgCGT
SerAGT
ArgCGA
ValGTA
ArgAGG
AspGAT
c
Contribution on comp 1
Isoacceptors
HisCAT
HisCAC
GlyGGA
ValGTT
ValGTC
ThrACG
ValGTA
ProCCG
EndTAA
SerAGT
LeuCTC
GlyGGT
GluGAG
GluGAA
EndTGA
Outcome
Down
Up
d
Contribution on comp 1
Total frequency
SerTCT
GlnCAA
ValGTT
IleATA
AspGAT
ArgAGG
SerAGT
EndTAA
LeuCTC
ValGTC
CysTGT
GlyGGT
HisCAC
GluGAA
GlnCAG
Outcome
Down
Up
e
-15.00
-2.00 2.00
15.00
id
Group
Glucose starvation
Up
AA transmembrane
Up
AA transport
Up
BMP signaling pathway
Up
ISR signaling
Up
Na homeostasis
Down
ER UPR
Up
negative regulation of stem cell proliferation
Down
snRNP assembly
Down
ATP
Down
mitochondrial ATP synthesis
Down
chromosome condensation
Down
LSU-rRNA
Down
miRNA gene silencing
Up
intracilliary anterograde transport
Down
microtubule-based process
Down
poly(A) tail shortening
Up
cytoskeleton transport
Down
stress granule assembly
Up
telomerase RNA localization
Down
mRNA splicing
Up/Down
tricarboxylic acid cycle
Down
prespliceosome assembly
Down
tRNA aminoacylation
Up
heart contraction
Down
f
PLS-DA on Codons
PLS-DA 1
PLS-DA 2
Legend
Down
Up
g
Correlation Circle Plot
Component 1
Component 2
ArgAGG
EndTAA
LeuTTG
EndTGA
ArgCGA
ArgCGT
ArgCGG
GlnCAG
SerAGC
h
Contribution on comp 1
Outcome
Down
Up

**Supplementary figure 8: ATF4 transcribed genes are G/C biased.** Enrichment of ATF4 pathway after Rotenone (**a**), antimycin A (**b**), or arsenite (**c**) exposure. **d:** Isoacceptors codon frequencies analysis of ATF4 transcribed genes retrieved from TRRUST database. **e:** Collected analysis of the ATF4 pathway represented as T-stat versus the genome average. **f:** Expression of different ATF4 transcribed genes in the Ribo-seq datasets.

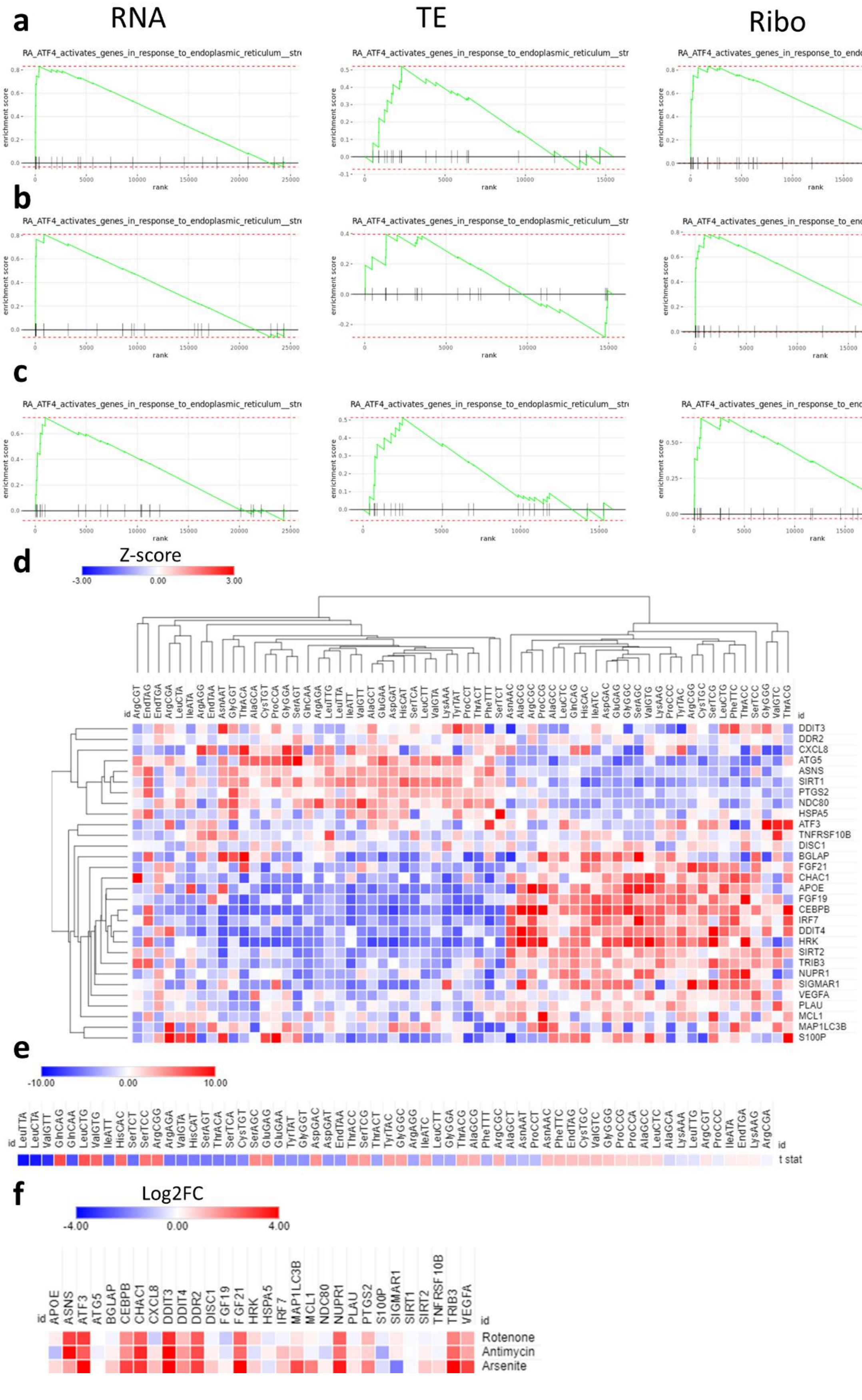

a
RNA
TE
Ribo
RA_ATF4_activates_genes_in_response_to_endoplasmic_reticulum__str
enrichment score
rank
b
c
d
Z-score
-3.00
0.00
3.00
id
ArgCGT EndTAG EndTGA ArgCGA LeuCTA IleATA ArgAGG EndTAA AsnAAT GlyGGT ThrACA AlaGCA CysTGT ProCCA GlyGGA SerAGT GlnCAA ArgAGA LeuTTG LeuTTA IleATT ValGTT AlaGCT GluGAA AspGAT HisCAT SerTCA LeuCTT ValGTA LysAAA TyrTAT ProCCT ThrACT PheTTT SerTCT AsnAAC AlaGCG ArgCGC ProCCG AlaGCC LeuCTC GlnCAG HisCAC IleATC AspGAC GluGAG GlyGGC SerAGC ValGTG LysAAG ProCCC TyrTAC ArgCGG CysTGC SerTCG LeuCTG PheTTC ThrACC SerTCC GlyGGG ValGTC ThrACG
DDIT3 DDR2 CXCL8 ATG5 ASNS SIRT1 PTGS2 NDC80 HSPA5 ATF3 TNFRSF10B DISC1 BGLAP FGF21 CHAC1 APOE FGF19 CEBPB IRF7 DDIT4 HRK SIRT2 TRIB3 NUPR1 SIGMAR1 VEGFA PLAU MCL1 MAP1LC3B S100P
e
-10.00
0.00
10.00
LeuTTA LeuCTA ValGTT GlnCAG GlnCAA LeuCTG ValGTG IleATT HisCAC SerTCT SerTCC ArgCGG ArgAGA ValGTA HisCAT SerAGT ThrACA SerTCA CysTGT SerAGC GluGAG GluGAA TyrTAT GlyGGT AspGAC AspGAT EndTAA ThrACC SerTCG ThrACT TyrTAC GlyGGC ArgAGG IleATC LeuCTT GlyGGA ThrACG AlaGCG PheTTT ArgCGC AlaGCT AsnAAT ProCCT AsnAAC PheTTC EndTAG CysTGC ValGTC GlyGGG ProCCG ProCCA AlaGCC LeuCTC AlaGCA LysAAA LeuTTG ArgCGT ProCCC IleATA EndTGA LysAAG ArgCGA
t stat
f
Log2FC
-4.00
0.00
4.00
APOE ASNS ATF3 ATG5 BGLAP CEBPB CHAC1 CXCL8 DDIT3 DDIT4 DDR2 DISC1 FGF19 FGF21 HRK HSPA5 IRF7 MAP1LC3B MCL1 NDC80 NUPR1 PLAU PTGS2 S100P SIGMAR1 SIRT1 SIRT2 TNFRSF10B TRIB3 VEGFA
Rotenone
Antimycin
Arsenite

**Supplementary figure 9: Impact of ALKBH1 on tRNA modifications.** LC-MS/MS spectral peaks after ALKBH1 KO (**a:** f5C, **b:** hm5Cm, and **c:** f5Cm). **d:** Cell viability after ALKBH1 KO and exposure to different ETC inhibitors for 4 hours (Rotenone 80µM, TTFA 1.5mM, Antimycin A 50µg/ml, Potassium Cyanide (KCN) 15mM, and Oligomycin 20µM) as well as Arsenite 600µM.

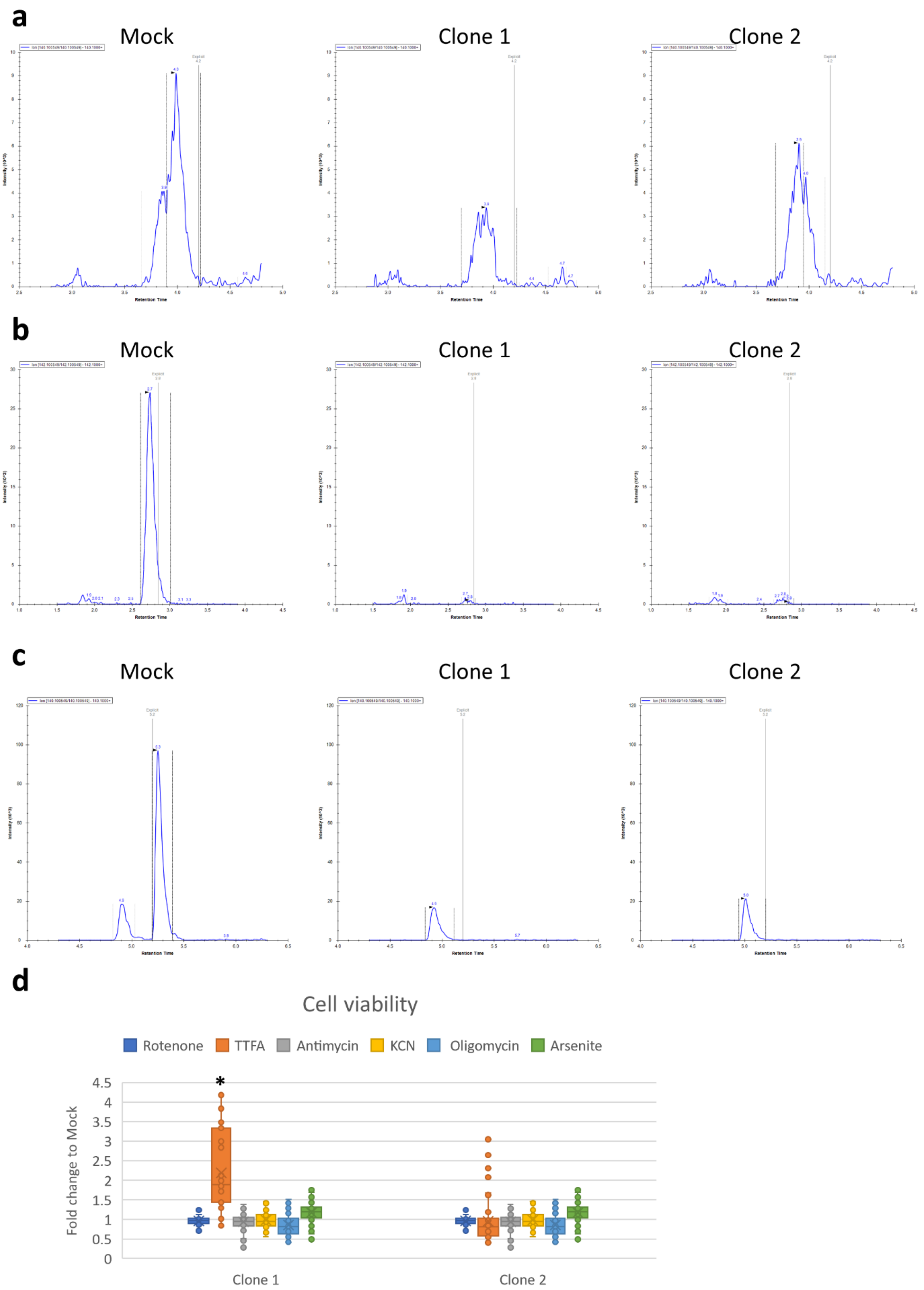
a
Mock
Clone 1
Clone 2
b
Mock
Clone 1
Clone 2
c
Mock
Clone 1
Clone 2
d
Cell viability
Rotenone
TTFA
Antimycin
KCN
Oligomycin
Arsenite
Fold change to Mock
4.5
4
3.5
3
2.5
2
1.5
1
0.5
0
*
Clone 1
Clone 2

**Supplementary figure 10: ALKBH1 KO leads to translational stress. a:** Puromycin incorporation assay after ALKBH1 KO with quantification graph. **b:** Western blot of eIF2α and its phosphorylated form (p- eIF2α). **c:** Quantification of eIF2α and p- eIF2α protein expression. **d:** Western blot of P38 and P70S6K and their phosphorylated forms (p-P38 and p-P70S6K). **e:** Quantification of P38 and p-P38 protein expression. **f:** Quantification of P70S6K and p-P70S6K protein expression. Asterisk: Fold change > 1.5 and $p < 0.05$.

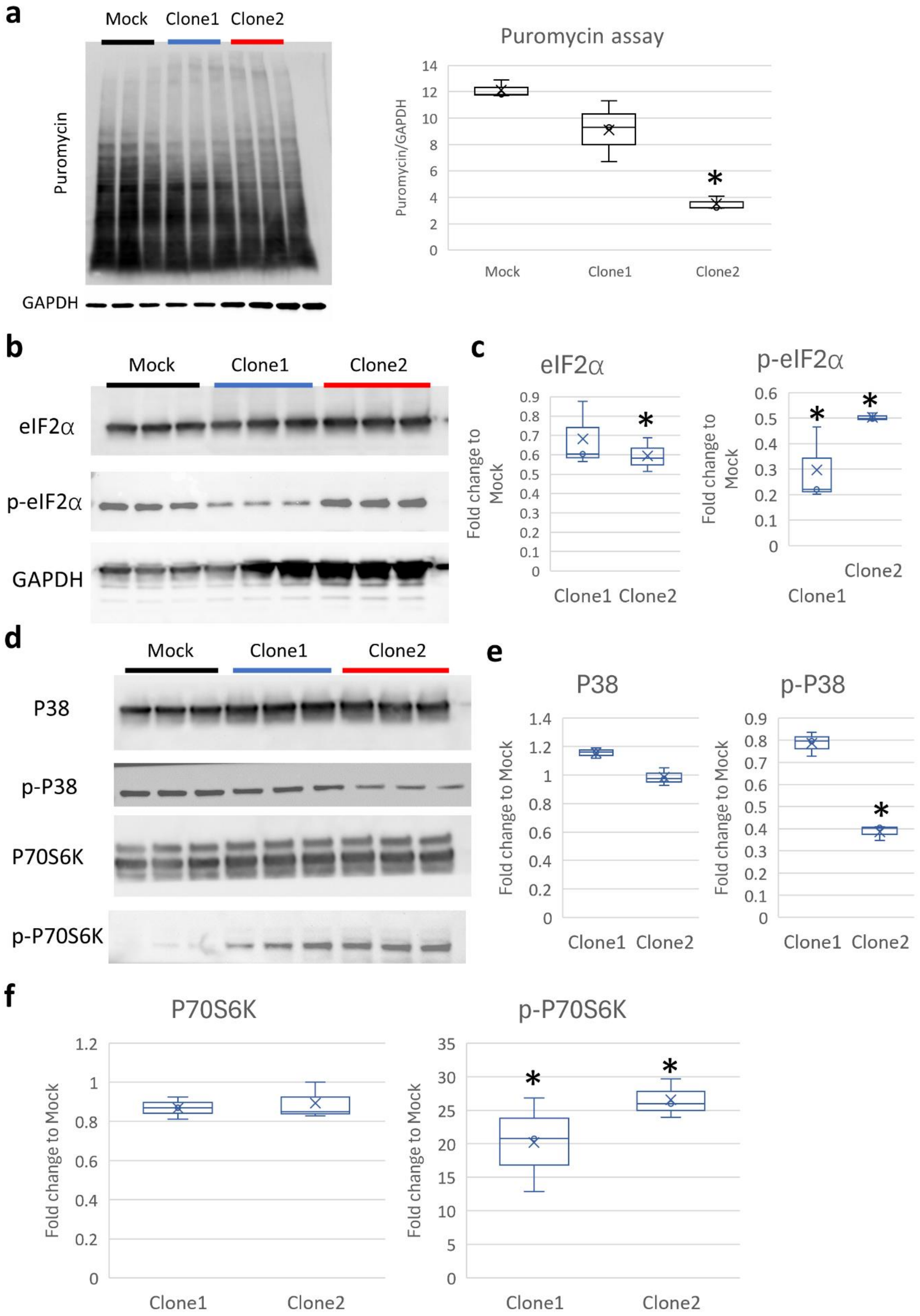
a
Mock
Clone1
Clone2
Puromycin
GAPDH
Puromycin assay
Puromycin/GAPDH
Mock
Clone1
Clone2
b
Mock
Clone1
Clone2
eIF2α
p-eIF2α
GAPDH
c
eIF2α
Fold change to Mock
Clone1
Clone2
p-eIF2α
Fold change to Mock
Clone1
Clone2
d
Mock
Clone1
Clone2
P38
p-P38
P70S6K
p-P70S6K
e
P38
Fold change to Mock
Clone1
Clone2
p-P38
Fold change to Mock
Clone1
Clone2
f
P70S6K
Fold change to Mock
Clone1
Clone2
p-P70S6K
Fold change to Mock
Clone1
Clone2

**Supplementary figure 11: ALKBH1 KO leads to mitochondrial dysfunction. a-h:** Different measurements derived from Seahorse assay. Asterisk: $p < 0.05$.

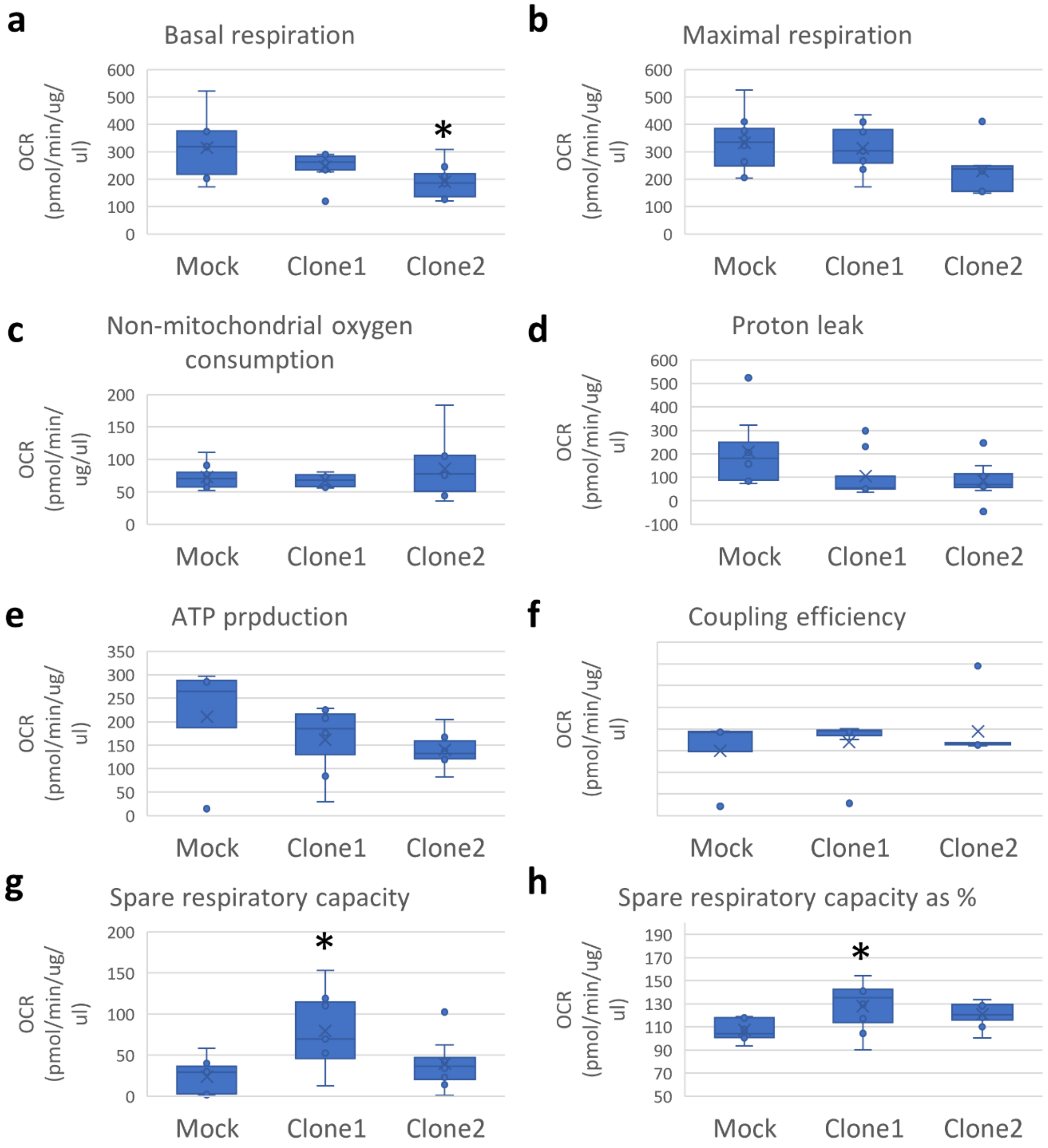
a
Basal respiration
OCR (pmol/min/ug/ul)
600
500
400
300
200
100
0
*
Mock
Clone1
Clone2
b
Maximal respiration
OCR (pmol/min/ug/ul)
600
500
400
300
200
100
0
Mock
Clone1
Clone2
c
Non-mitochondrial oxygen consumption
OCR (pmol/min/ug/ul)
200
150
100
50
0
Mock
Clone1
Clone2
d
Proton leak
OCR (pmol/min/ug/ul)
600
500
400
300
200
100
0
-100
Mock
Clone1
Clone2
e
ATP prpduction
OCR (pmol/min/ug/ul)
350
300
250
200
150
100
50
0
Mock
Clone1
Clone2
f
Coupling efficiency
OCR (pmol/min/ug/ul)
Mock
Clone1
Clone2
g
Spare respiratory capacity
OCR (pmol/min/ug/ul)
200
150
100
50
0
*
Mock
Clone1
Clone2
h
Spare respiratory capacity as %
OCR (pmol/min/ug/ul)
190
170
150
130
110
90
70
50
*
Mock
Clone1
Clone2

**Supplementary figure 12: tRNA-Q KO impacts tRNA modifications. a:** Protein expression of QTRT1 and QTRT2 after 8 hours of stress exposure in wild-type cells. **b:** T7 endonuclease assay to validate DNA cleavage after CRISPR KO induction. Lanes (from left): Mock, QTRT1 KO, QTRT2 KO, Double KO (QTRT1 primer), Double KO (QTRT2 primer). **c-d:** Mass spectrometry signals for manQ and galQ after QTRT1, QTRT2, or double KO. **e:** Puromycin incorporation assay. **f:** Quantification of western blot presented in figure 5g.

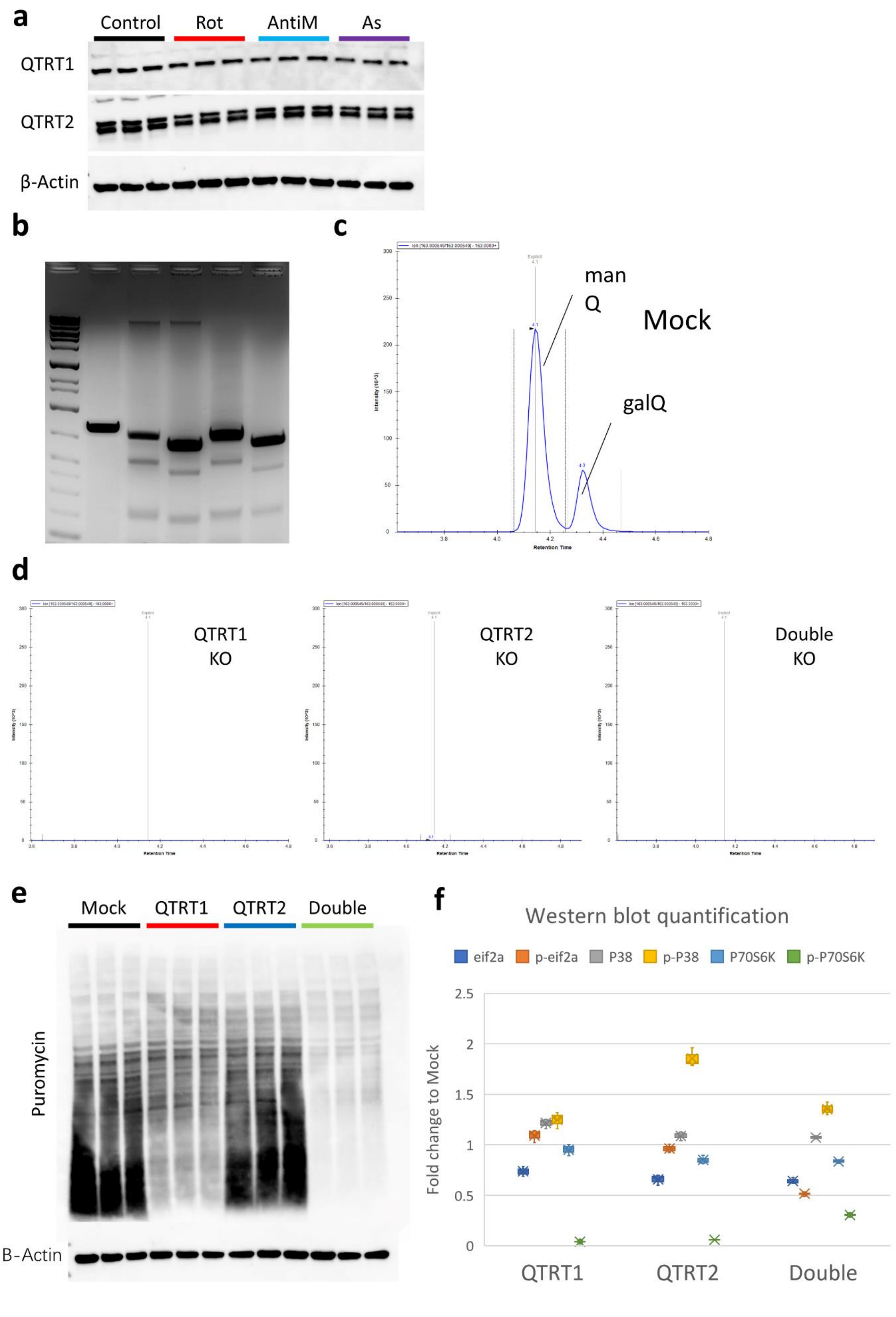
a
Control
Rot
AntiM
As
QTRT1
QTRT2
β-Actin
b
c
man
Q
Mock
galQ
Retention Time
d
QTRT1
KO
QTRT2
KO
Double
KO
e
Mock
QTRT1
QTRT2
Double
Puromycin
B-Actin
f
Western blot quantification
eif2a
p-eif2a
P38
p-P38
P70S6K
p-P70S6K
Fold change to Mock
2.5
2
1.5
1
0.5
0
QTRT1
QTRT2
Double

**Supplementary figure 13:** **a:** G3BP1 (a marker for SG) staining. **b:** EDC4 (a marker for P-bodies) staining and its quantification. Asterisk: fold change > 1.5 and $p < 0.05$.

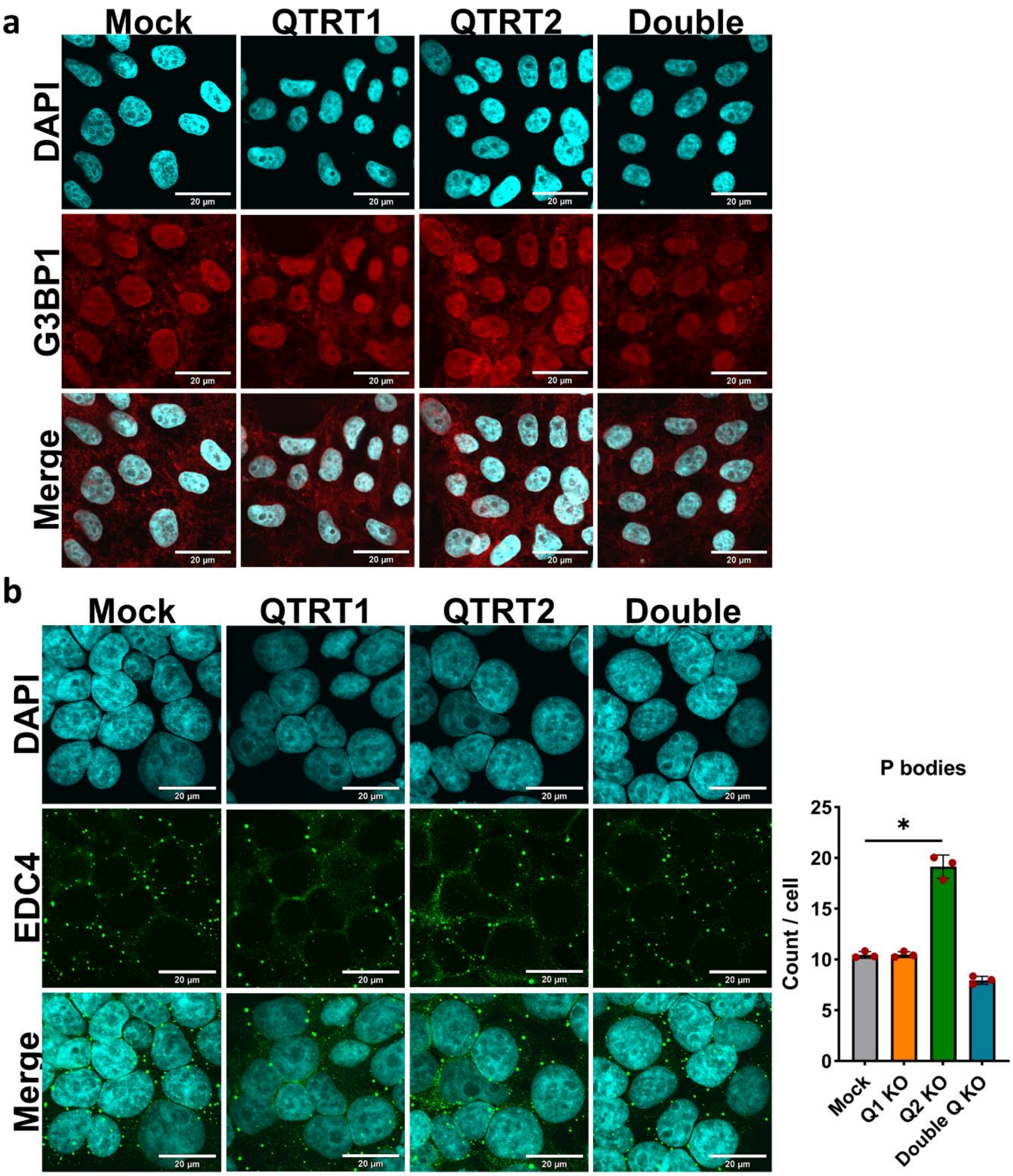
a
Mock
QTRT1
QTRT2
Double
DAPI
G3BP1
Merge
20 μm
b
Mock
QTRT1
QTRT2
Double
DAPI
EDC4
Merge
20 μm
P bodies
Count / cell
0
5
10
15
20
25
*
Mock
Q1 KO
Q2 KO
Double Q KO

**Supplementary figure 14: a:** Isoacceptors codon frequencies in the Ribo-seq dataset. **b:** Total codon frequencies in the Ribo-seq dataset. **c:** Isoacceptors codon frequencies in the TE dataset. **d:** Total codon frequencies in the TE dataset.

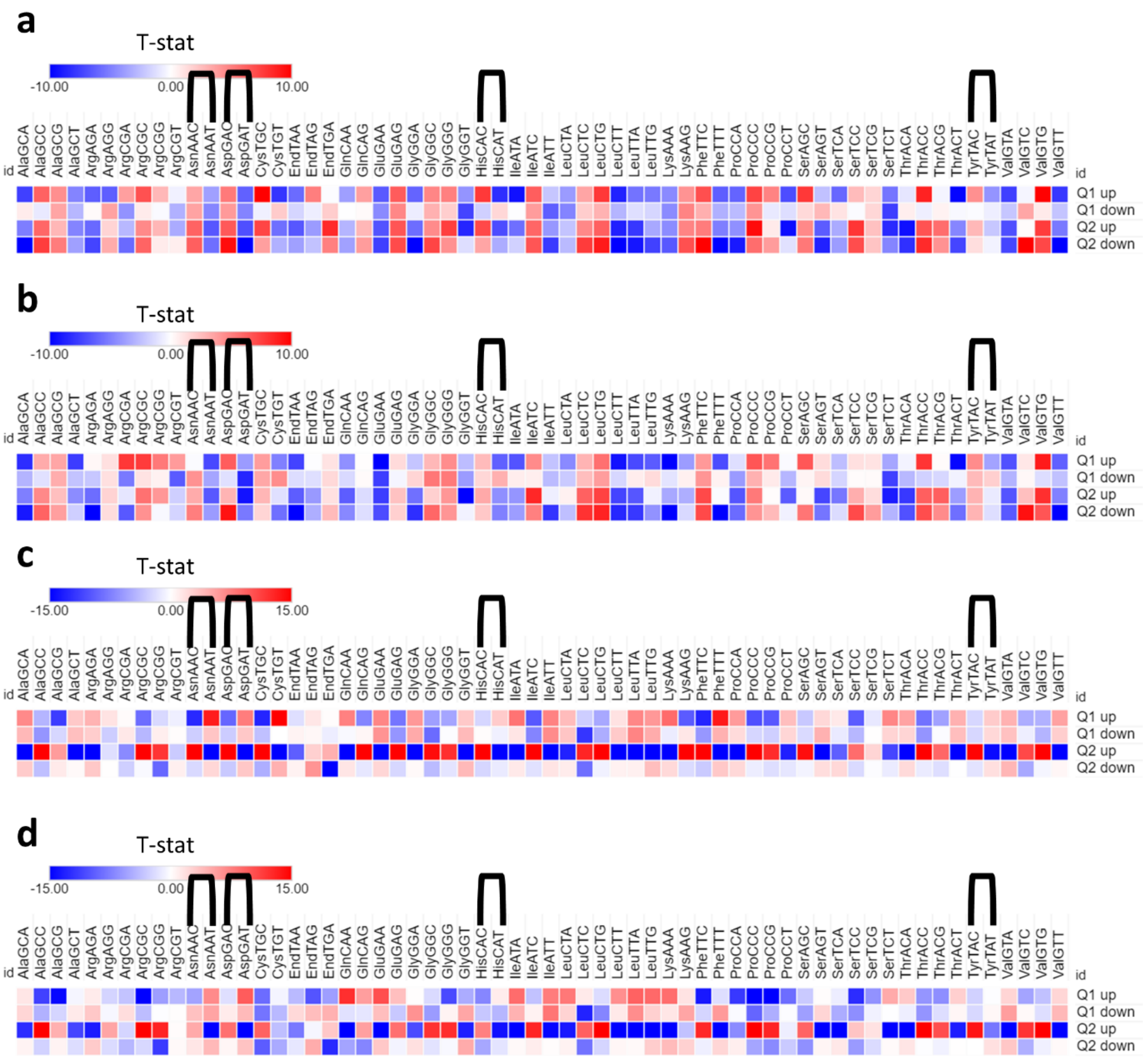

a
T-stat
-10.00
0.00
10.00
id AlaGCA AlaGCC AlaGCG AlaGCT ArgAGA ArgAGG ArgCGA ArgCGC ArgCGG ArgCGT AsnAAC AsnAAT AspGAC AspGAT CysTGC CysTGT EndTAA EndTAG EndTGA GlnCAA GlnCAG GluGAA GluGAG GlyGGA GlyGGC GlyGGG GlyGGT HisCAC HisCAT IleATA IleATC IleATT LeuCTA LeuCTC LeuCTG LeuCTT LeuTTA LeuTTG LysAAA LysAAG PheTTC PheTTT ProCCA ProCCC ProCCG ProCCT SerAGC SerAGT SerTCA SerTCC SerTCG SerTCT ThrACA ThrACC ThrACG ThrACT TyrTAC TyrTAT ValGTA ValGTC ValGTG ValGTT id
Q1 up
Q1 down
Q2 up
Q2 down
b
T-stat
-10.00
0.00
10.00
id AlaGCA AlaGCC AlaGCG AlaGCT ArgAGA ArgAGG ArgCGA ArgCGC ArgCGG ArgCGT AsnAAC AsnAAT AspGAC AspGAT CysTGC CysTGT EndTAA EndTAG EndTGA GlnCAA GlnCAG GluGAA GluGAG GlyGGA GlyGGC GlyGGG GlyGGT HisCAC HisCAT IleATA IleATC IleATT LeuCTA LeuCTC LeuCTG LeuCTT LeuTTA LeuTTG LysAAA LysAAG PheTTC PheTTT ProCCA ProCCC ProCCG ProCCT SerAGC SerAGT SerTCA SerTCC SerTCG SerTCT ThrACA ThrACC ThrACG ThrACT TyrTAC TyrTAT ValGTA ValGTC ValGTG ValGTT id
Q1 up
Q1 down
Q2 up
Q2 down
c
T-stat
-15.00
0.00
15.00
id AlaGCA AlaGCC AlaGCG AlaGCT ArgAGA ArgAGG ArgCGA ArgCGC ArgCGG ArgCGT AsnAAC AsnAAT AspGAC AspGAT CysTGC CysTGT EndTAA EndTAG EndTGA GlnCAA GlnCAG GluGAA GluGAG GlyGGA GlyGGC GlyGGG GlyGGT HisCAC HisCAT IleATA IleATC IleATT LeuCTA LeuCTC LeuCTG LeuCTT LeuTTA LeuTTG LysAAA LysAAG PheTTC PheTTT ProCCA ProCCC ProCCG ProCCT SerAGC SerAGT SerTCA SerTCC SerTCG SerTCT ThrACA ThrACC ThrACG ThrACT TyrTAC TyrTAT ValGTA ValGTC ValGTG ValGTT id
Q1 up
Q1 down
Q2 up
Q2 down
d
T-stat
-15.00
0.00
15.00
id AlaGCA AlaGCC AlaGCG AlaGCT ArgAGA ArgAGG ArgCGA ArgCGC ArgCGG ArgCGT AsnAAC AsnAAT AspGAC AspGAT CysTGC CysTGT EndTAA EndTAG EndTGA GlnCAA GlnCAG GluGAA GluGAG GlyGGA GlyGGC GlyGGG GlyGGT HisCAC HisCAT IleATA IleATC IleATT LeuCTA LeuCTC LeuCTG LeuCTT LeuTTA LeuTTG LysAAA LysAAG PheTTC PheTTT ProCCA ProCCC ProCCG ProCCT SerAGC SerAGT SerTCA SerTCC SerTCG SerTCT ThrACA ThrACC ThrACG ThrACT TyrTAC TyrTAT ValGTA ValGTC ValGTG ValGTT id
Q1 up
Q1 down
Q2 up
Q2 down

**Supplementary figure 15: tRNA-Q loss impacts mitochondrial function. a-b:** Western blot and quantification of different respiratory complex proteins in the KO cell lines. **c:** Live cell confocal imaging of Mito tracker red and green in the KO cell lines.

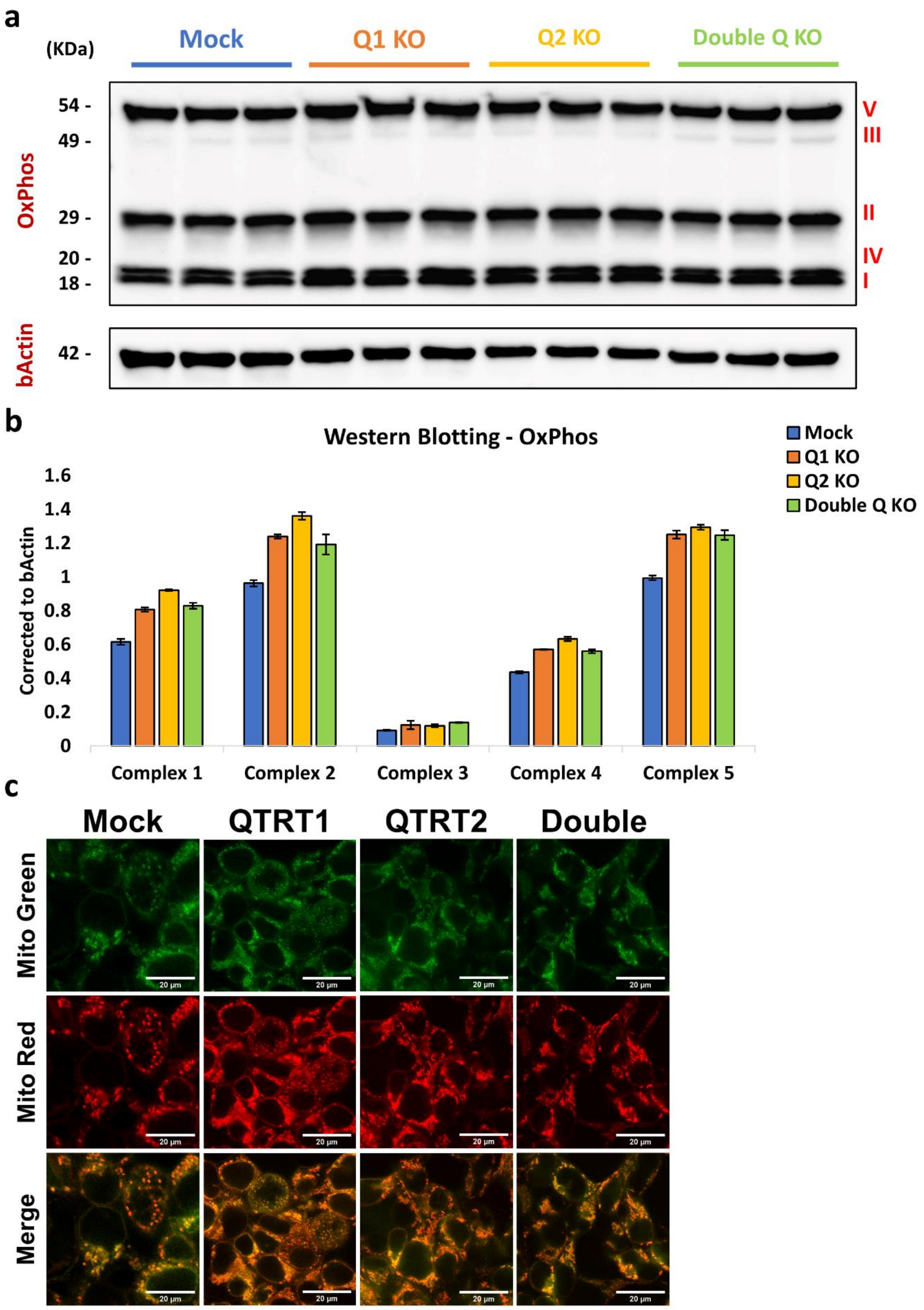
a
(KDa)
Mock
Q1 KO
Q2 KO
Double Q KO
54 -
49 -
29 -
20 -
18 -
V
III
II
IV
I
OxPhos
bActin
42 -
b
Western Blotting - OxPhos
Mock
Q1 KO
Q2 KO
Double Q KO
Corrected to bActin
1.6
1.4
1.2
1
0.8
0.6
0.4
0.2
0
Complex 1
Complex 2
Complex 3
Complex 4
Complex 5
c
Mock
QTRT1
QTRT2
Double
Mito Green
Mito Red
Merge

**Supplementary figure 16: tRNA-Q loss impacts mitochondrial function (Cont.) a:** Mitochondrial respiration (oxygen consumption rate) analysis from Seahorse assay. **b-i:** Various readouts from the seahorse assay. Asterisk: $p < 0.05$.

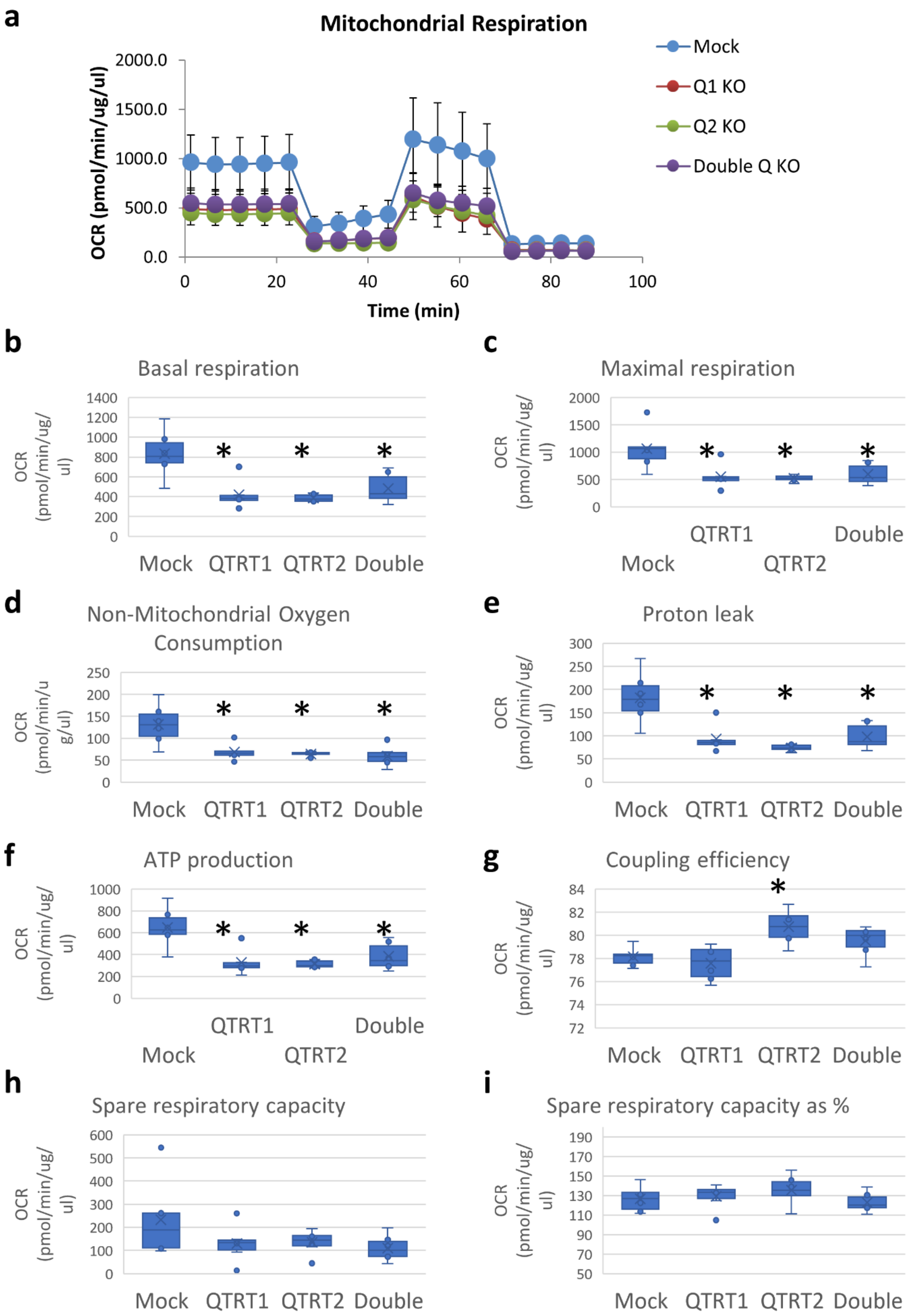

a
Mitochondrial Respiration
Mock
Q1 KO
Q2 KO
Double Q KO
OCR (pmol/min/ug/ul)
Time (min)
b
Basal respiration
c
Maximal respiration
d
Non-Mitochondrial Oxygen Consumption
e
Proton leak
f
ATP production
g
Coupling efficiency
h
Spare respiratory capacity
i
Spare respiratory capacity as %
OCR (pmol/min/ug/ul)
Mock
QTRT1
QTRT2
Double

**Supplementary figure 17:** Cell viability analysis after stress exposure of tRNA-Q KO cell lines in different media. This data is detailed format for the heatmaps presented in figure 8a-d.

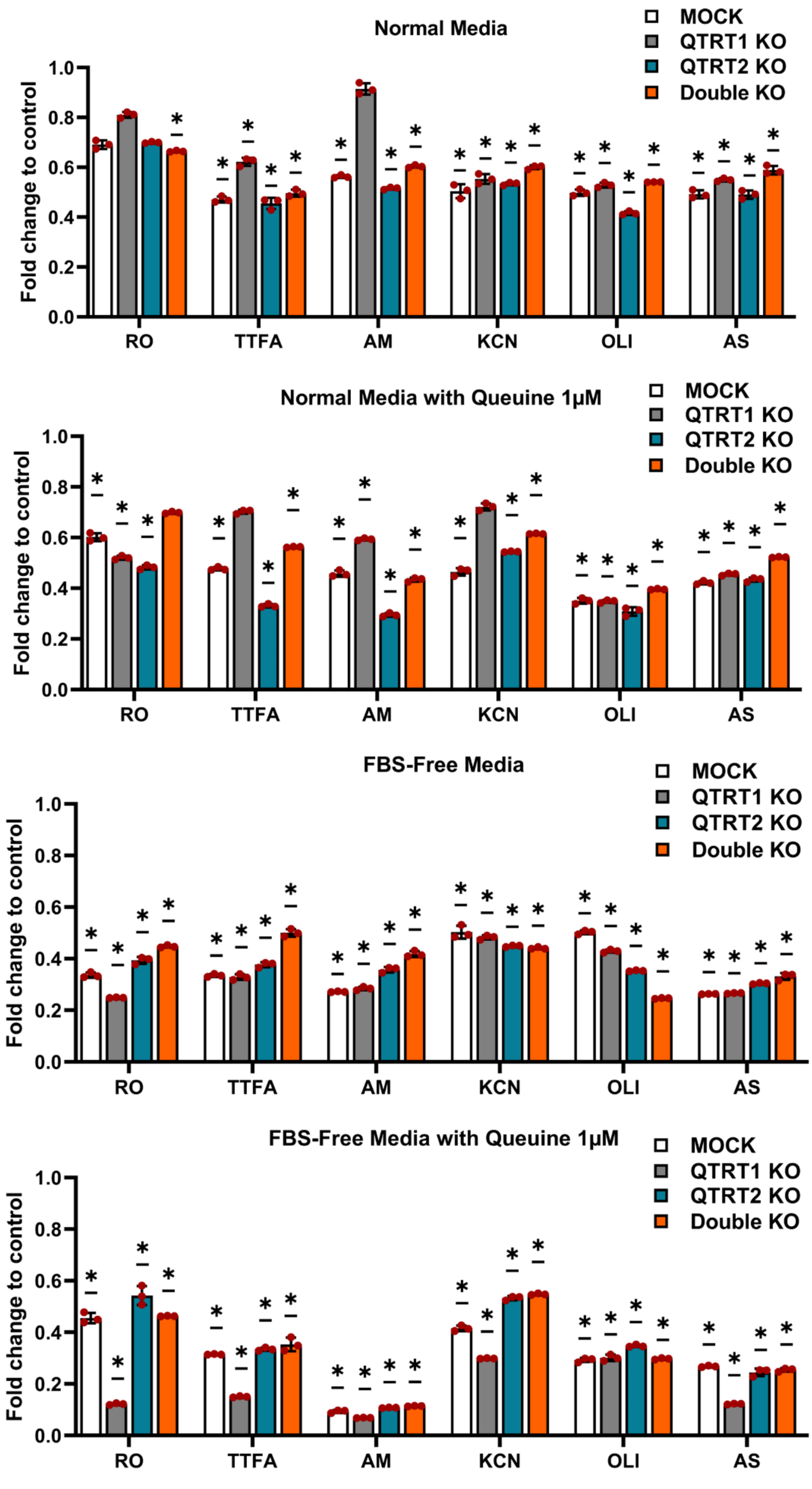
Normal Media
MOCK
QTRT1 KO
QTRT2 KO
Double KO
Fold change to control
RO
TTFA
AM
KCN
OLI
AS
Normal Media with Queuine 1μM
FBS-Free Media
FBS-Free Media with Queuine 1μM

**Supplementary figure 18:** Mass spectrometry analysis of tRNA modifications after 4 hours of stress exposure of the tRNA-Q and Mock KO cell lines to ETC inhibitors (Rotenone 80µM, TTFA 1.5mM, Antimycin A 50µg/ml, Potassium Cyanide (KCN) 15mM, and Oligomycin 20µM) as well as Arsenite 600µM. Data represented as log2 fold changes to the controls of each cell line. Asterisk: fold change > 1.5 and *p* < 0.05.

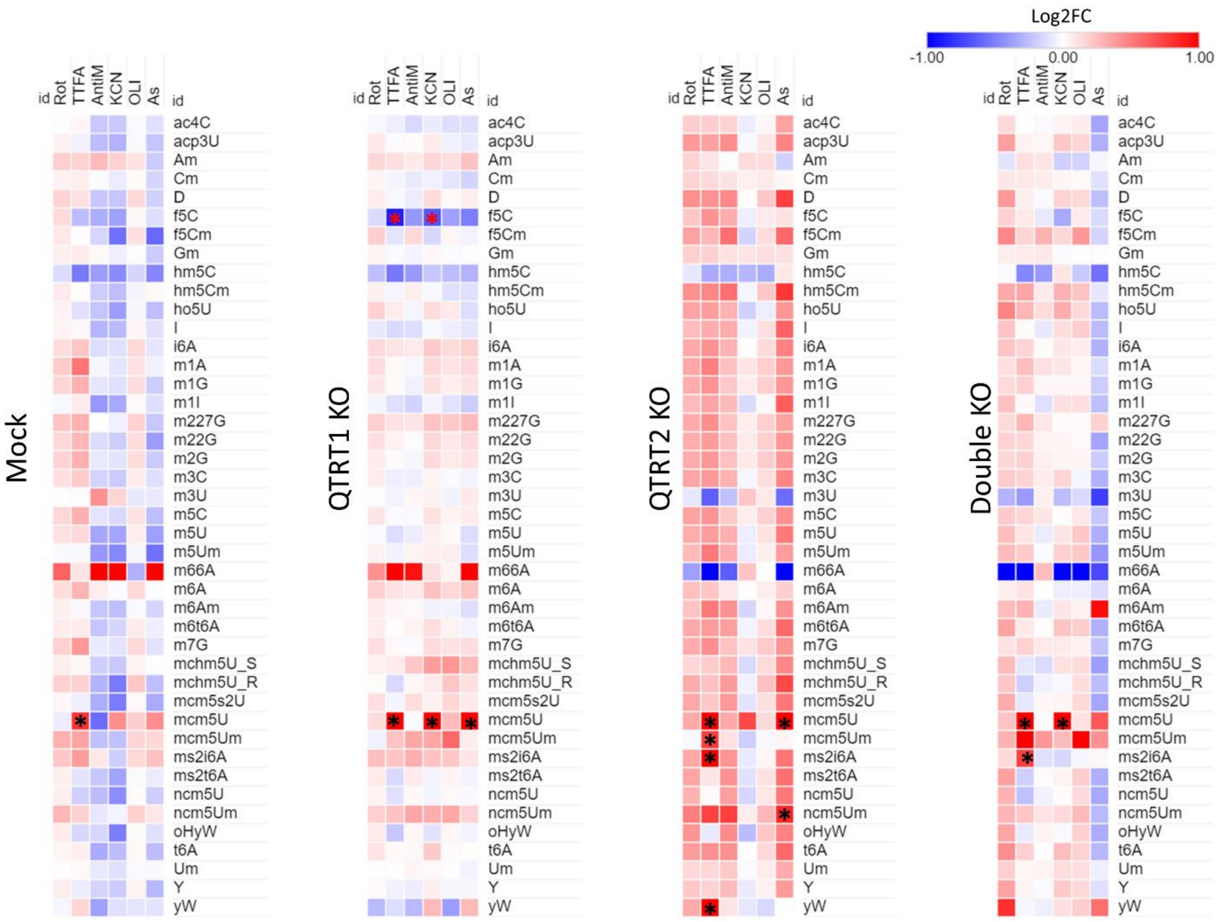

**Supplementary figure 19: Metabolomic analysis after tRNA-Q depletion. a:** LC-MS/MS based Targeted metabolomics. **b:** statistically significant metabolites on Anova (asterisk: FC > 1.5, $p < 0.05$ with Turkey's post hoc. **c:** GC-MS/MS based untargeted metabolomics. **d:** significant metabolites on ANOVA. Asterisk: FC > 1.5, $p < 0.05$ with Turkey's post hoc. **e:** GSH transsulfuration pathway analysis. Asterisk: FC > 1.5, $p < 0.05$ with Turkey's post hoc. **f:** Correlation analysis using all the metabolomics data.

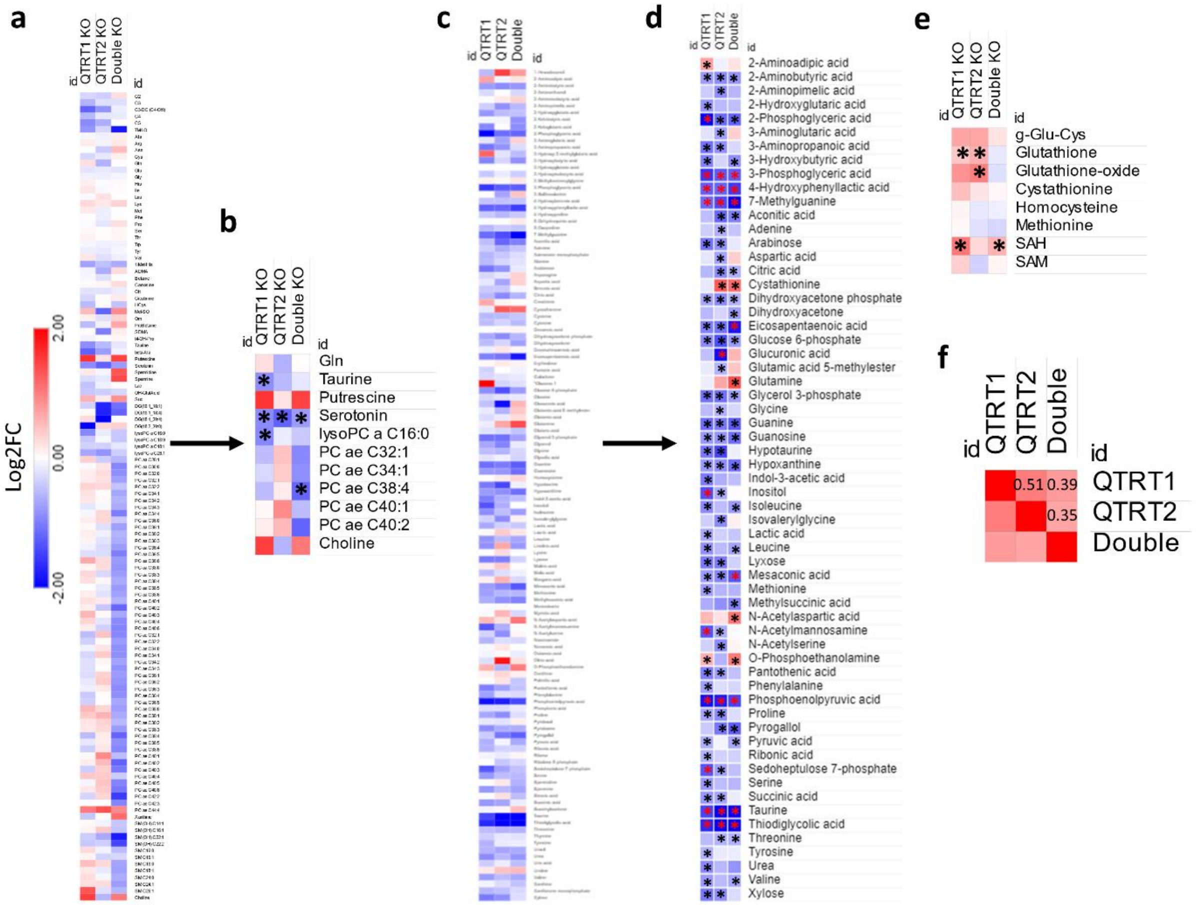

**Supplementary figure 20:** Metabolic pathway analysis in QTRT1 KO cells. **a:** Small molecules database (SMDB) pathway analysis. **b:** Kegg pathway analysis. Statistically significant metabolites used in the enrichment analysis were globally downregulated.

**a**

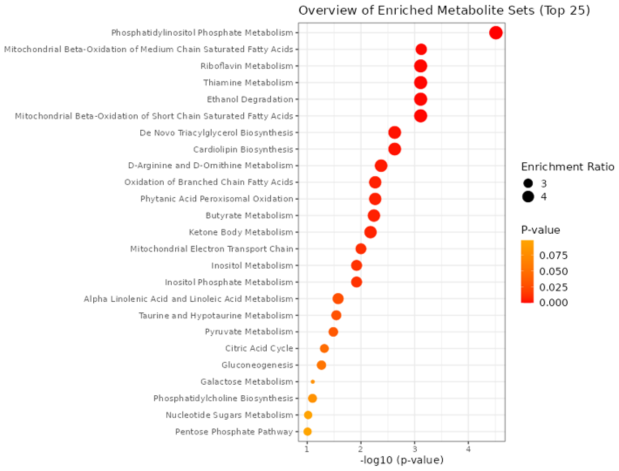


**b**

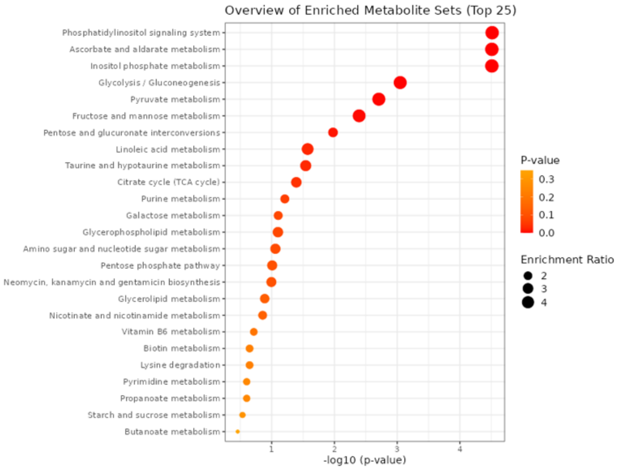

**Supplementary figure 21:** Metabolic pathway analysis in QTRT2 KO cells. **a:** Small molecules database (SMDB) pathway analysis. **b:** Kegg pathway analysis. Statistically significant metabolites used in the enrichment analysis were globally downregulated.

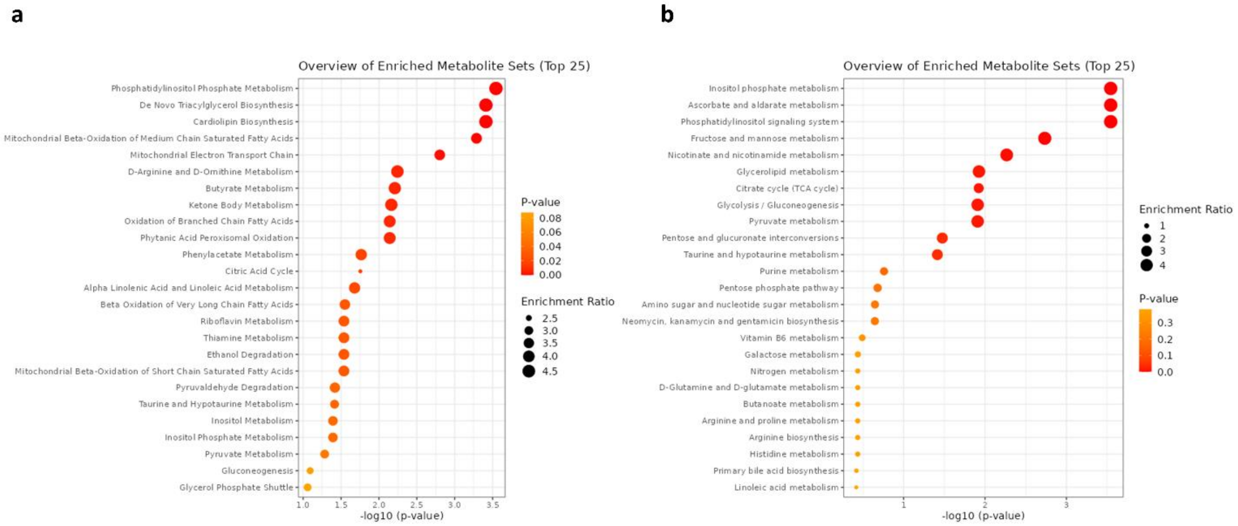

**Supplementary figure 22:** Metabolic pathway analysis in Double KO cells. **a:** Small molecules database (SMDB) pathway analysis. **b:** Kegg pathway analysis. Statistically significant metabolites used in the enrichment analysis were globally downregulated.

a

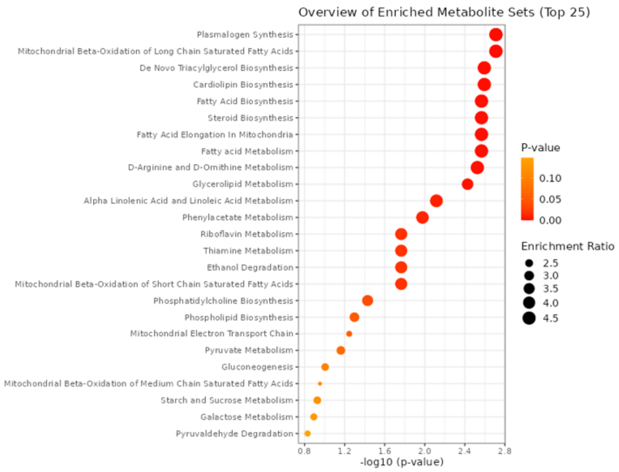


b

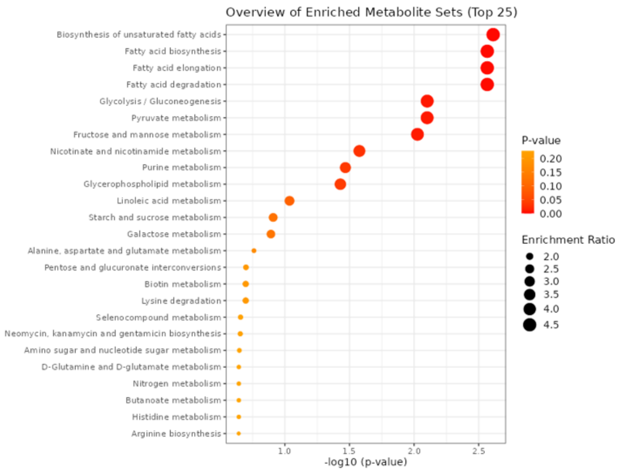
